# Molecular basis of interchain disulfide-bond formation in BMP-9 and BMP-10

**DOI:** 10.1101/2024.10.14.618187

**Authors:** Tristin A. Schwartze, Stefanie A. Morosky, Teresa L. Rosato, Amy Henrickson, Guowu Lin, Cynthia S. Hinck, Alexander B. Taylor, Shaun K. Olsen, Guillermo Calero, Borries Demeler, Beth L. Roman, Andrew P. Hinck

## Abstract

BMP-9 and BMP-10 are TGF-β family signaling ligands naturally secreted into blood. They act on endothelial cells and are required for proper development and maintenance of the vasculature. In hereditary hemorrhagic telangiectasia, regulation is disrupted due to mutations in the BMP-9/10 pathway, namely in the type I receptor ALK1 or the co-receptor endoglin. It has been demonstrated that BMP-9/10 heterodimers are the most abundant signaling species in the blood, but it is unclear how they form. Unlike other ligands of the TGF-β family, BMP-9 and -10 are secreted as a mixture of monomers and disulfide-linked dimers. Here, we show that the monomers are secreted in a cysteinylated form that crystallizes as a noncovalent dimer. Despite this, monomers do not self-associate at micromolar or lower concentrations and have reduced signaling potency compared to dimers. We further show using protein crystallography that the interchain disulfide of the BMP-9 homodimer adopts a highly strained syn-periplanar conformation. Hence, geometric strain across the interchain disulfide is responsible for the reduced propensity to dimerize, not the cysteinylation. Additionally, we show that the dimerization propensity of BMP-9 is lower than BMP-10 and these propensities can be reversed by swapping residues near the interchain disulfide that form attractive interactions with the opposing monomer. Finally, we discuss the implications of these observations on BMP-9/10 heterodimer formation.

## Introduction

Bone morphogenetic protein nine and ten (BMP-9 and BMP-10) are TGF-β family growth factors synthesized by hepatic stellate cells (1) and for BMP-10 also cardiomyocytes (2,3). They are secreted into the blood and signal on endothelial cells. This signaling is essential for the proper development and maintenance of the vasculature (3–6). In disease, dysfunction of the BMP-9/10 signaling pathway causes hereditary hemorrhagic telangiectasia (HHT) (7–11). HHT is characterized by the development of arteriovenous malformations (AVMs), direct connections between arteries and veins without intervening capillaries. AVMs can form in small vessels on the skin, nose, and gastrointestinal tract, where they are known as telangiectasias as well as in larger vessels in the lung, liver, and brain and can lead to epistaxis, internal bleeding, anemia, stroke, brain abscess, and high-output heart failure. Understanding the BMP-9/-10 signaling pathway will assist in the development of first-in-class targeted treatments for HHT patients.

BMP-9 and -10 are synthesized as pro-proteins and processed by furin to release growth factors comprised of two cystine-knotted monomers that homo- or hetero-dimerize (**Fig. S1**). In the blood, growth factors encounter their type-I receptor ALK1 (12) and co-receptor endoglin (13) on vascular endothelial cells. ALK1 binds across the dimer interface with extensive contacts on the interface helix resulting in high affinity and specificity for BMP-9 and -10 (14–16). Endoglin binds the convex surface of the β-strands with high specificity for BMP-9 and -10 (14,15,17), but it is displaced by a type-II receptor: ActRIIA, ActRIIB, or BMPRII. The resulting signaling complex consists of one dimeric ligand, two type-I receptors (ALK1), and two type-II receptors. Signaling complex formation allows the constitutively active type-II receptor kinase to phosphorylate and activate the ALK1 kinase, which phosphorylates the downstream transcriptional effector molecules, SMAD1, SMAD5, and SMAD9. In 85 - 96% of HHT cases, mutations in genes encoding the non-redundant signaling receptor and co-receptor of the pathway, *ALK1* and *ENG,* cause the disease (10,11).

BMP-9 and -10 are found as homodimers in the blood, but the predominant circulating form and signaling species is a BMP-9/10 heterodimer (18,19). The molecular basis for heterodimer formation is unknown. One mechanism is prodomain selection where swapped prodomains select for an opposing growth factor domain (20). Another mechanism is growth factor complementarity where the interface of one growth factor complements the interface of another.

BMP-9 and -10 are unique in the TGF-β family for recombinantly overexpressing as a mixture of monomers and disulfide-linked dimers (14,16,17,21,22). The BMP-9 and BMP-10 monomers have been reported to have a strong propensity to self-associate and form noncovalent dimers, which are resistant to disruption even in 1 M guanidinium hydrochloride (23). Most structures of BMP-9 and BMP-10 derive from crystals formed from monomer-dimer mixtures (14,16,17,21,22), and the resulting electron densities reflect contributions from both monomers and dimers. The densities are therefore difficult to interpret at the cysteines that form the interchain disulfide bond. These cysteines are modeled with two conformations **(Fig. 1A)**: one with the interchain disulfide and the other with free sulfhydryls where the side-chain χ_1_ dihedrals are rotated by 120° relative to the presumed disulfide-bonded conformation (17,21,22). In these models, there is no mechanism to prevent rotation about χ_1_ and covalent dimer formation by spontaneous oxidation. Hence, there is no mechanistic basis for poor covalent dimer formation in BMP-9 and BMP-10. It has been suggested that the interchain disulfide of BMP-9 is more readily reduced compared to the interchain disulfide of BMP-6 based on redox titrations with oxidized and reduced glutathione (21). While evidence of a more positive reduction potential could be consistent with the synthesis and secretion of monomers, it provides no structural basis for the failure of monomers to dimerize.

**Figure 1:**
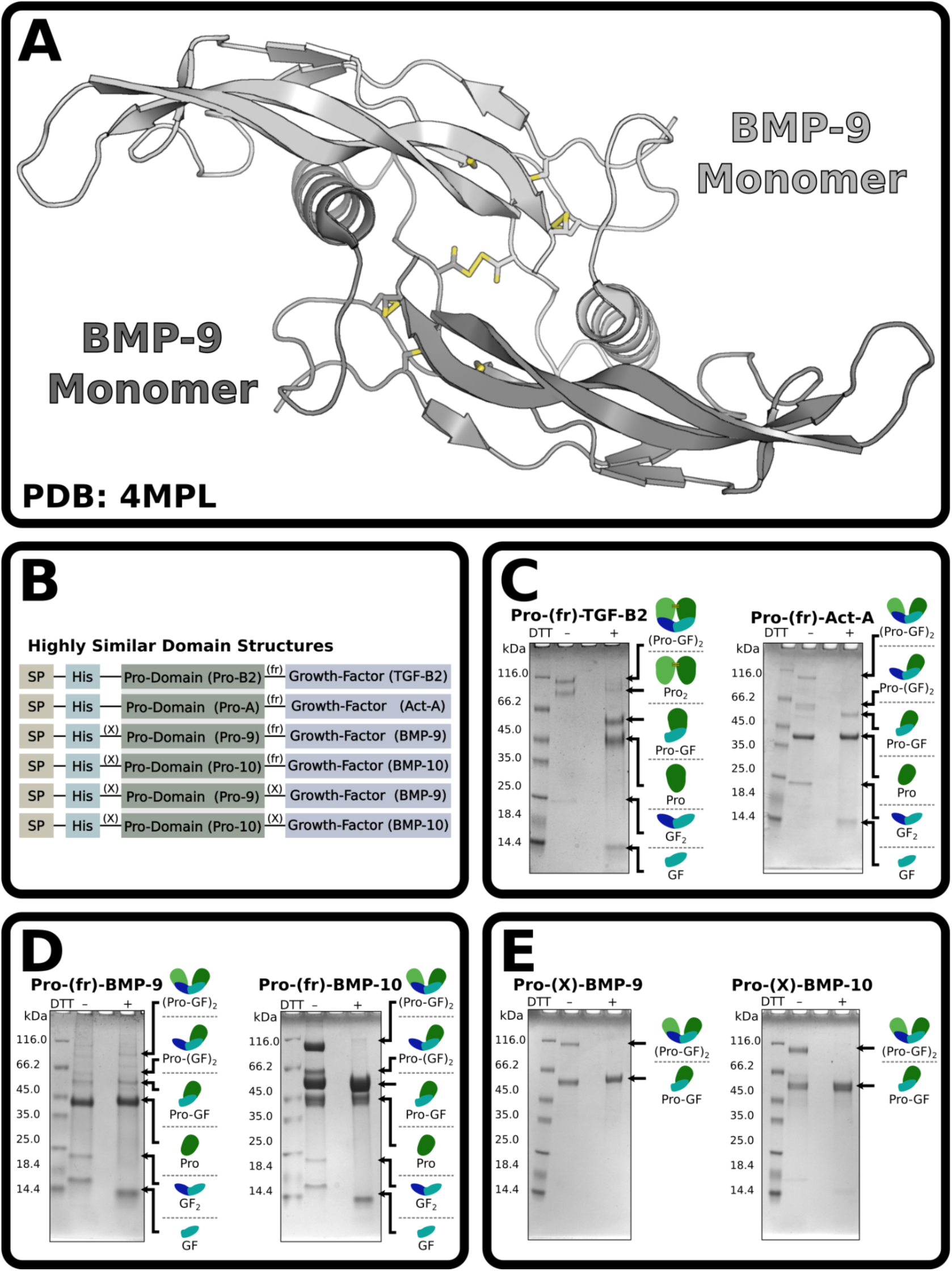
BMP-9 and BMP-10 express as monomers. **(A)** Overall structure of BMP-9 growth factor (PDB: 4MPL). **(B)** Basic domain structure of the constructs used throughout the paper. SP, X, and fr designate signal-peptide, factor X_a_ sites, and furin sites, respectively. **(C-E)** SDS-PAGE of IMAC column purified and either reduced or non-reduced Pro-TGF-β2 and Pro-Act-A representative members of the TGF-β family **(C)**, or wild-type BMP-9 and BMP-10 with unmodified furin processing sites between the prodomain and growth factor domain **(D)**, BMP-9 and BMP-10 with furin processing sites replaced by factor-X processing sites **(E)**. In **(C-E)**, small diagrams of each protein in the mixture are shown on the right of each gel.

Here, experiments are performed with the objective of understanding the poor propensity for covalent homodimer formation in BMP-9 and -10. Size exclusion chromatography with multi-angle light scattering (SEC-MALS) shows that BMP-9 monomers do not self-associate at micromolar or lower concentrations and signaling assays show that the monomers have reduced signaling potency compared to dimers. Mass spectrometry of the purified BMP-9 and -10 monomers, as well as crystallization of the BMP-9 monomer as a noncovalent dimer, show the interchain cysteine is modified by cysteinylation. The crystal structure of the purified BMP-9 dimer reveals the interchain disulfide adopts a highly strained syn-periplanar conformation and is much more sensitive to radiation damage than the intrachain disulfide bonds of the cystine knot. Structural analysis shows that the registration of the monomers in the homodimer complex is non-ideal for disulfide bond formation. Thus, geometric strain is responsible for the poor dimerization propensity, not monomer cysteinylation. Further, we show that the dimerization propensity of BMP-9 is lower than that of BMP-10. Swapping glycine-serine and lysine-arginine residues structurally adjacent to the interchain cysteine in BMP-9 and BMP-10, respectively, generates mutants with swapped dimerization propensities. We discuss the possible implications of these findings for the formation of BMP-9/10 heterodimers.

## Results

### BMP-9 and BMP-10 are produced as monomers

When recombinantly expressed in mammalian cells as full-length proteins, most TGF-β family growth factors are secreted exclusively as covalent dimers. Furin processing between the prodomain and growth factor is also sometimes incomplete. We overexpressed N-terminally histidine-tagged furin processible (fr) Pro-TGF-β2 and Pro-Activin-A (Pro-Act-A) as secreted proteins (**Fig. 1B**) in expi293 suspension cultured HEK293 cells then purified the protein from conditioned medium based on the histidine tag. SDS-PAGE under non-reducing conditions showed both dimer expression and incomplete furin processing (**Fig. 1C**). Under reducing conditions, the dimeric forms are converted to the corresponding monomeric forms. Owing to disulfide linkages in the prodomain of Pro-TGF-β2, but not Pro-Act-A, the prodomain of Pro-TGF-β2 is dimeric under non-reducing conditions, but the prodomain of Pro-Act-A is monomeric (24,25).

Unlike other TGF-β family members, BMP-9 and -10 are secreted as monomer-dimer mixtures (16,17,21,22,26). When overexpressed in expi293 cells with unmodified furin processing sites, Pro-BMP-9 and Pro-BMP-10 are secreted as a partially processed mixture of monomers and dimers. The monomer-dimer mixture is clearly evident by the growth factors which are 12 and 24 kDa in size (**Fig. 1D**). The combination of a monomer-dimer mixture and partial processing leads to more complex results for the high molecular weight forms. To simplify the purification and increase the amount of growth factor that can be obtained, we replaced the furin processing site of Pro-BMP-9 and -10 with an orthogonal factor X_a_ processing site (X) (**Fig. 1B**). Using these constructs, we found that Pro-BMP-9 and -10 are still secreted as monomer-dimer mixtures, but the mixtures are less complex since they express exclusively in an unprocessed form (**Fig. 1E**). In this study, we used this simplified system to obtain fully purified growth factor monomers and dimers by fractionating these from factor X_a_ digests of Pro-BMP-9 or Pro-BMP-10 monomer-dimer mixtures using C18 reverse phase chromatography.

### Monomers are monomeric in solution

Monomers represent a significant portion of secreted BMP-9 and -10, but based on size exclusion chromatography (SEC) they have been reported to form highly-stable noncovalent dimers (23) and signal with potencies that are only slightly different from dimers (21). We sought to quantify these observations with the purified monomers and dimers that we report here. BMP-9 monomer in complex with excess ActRIIa extracellular domain was run on size exclusion chromatography with multiangle light scattering (SEC-MALS) at pH 7.5 (**Fig. 2A**). A complex and receptor peak eluted (**Fig. 2A,B**). At the lowest injected concentration of BMP-9 monomer (6.58 µM), the light scattering of the complex peak is consistent with a solution monomer in complex with a receptor (observed and expected masses, 24.3 and 24.4 kDa, respectively). As the concentration increases (20.98 µM and 88.48 µM), the elution volume of the peak decreases and the average mass increases. This is consistent with dimerization via mass action.

**Figure 2:**
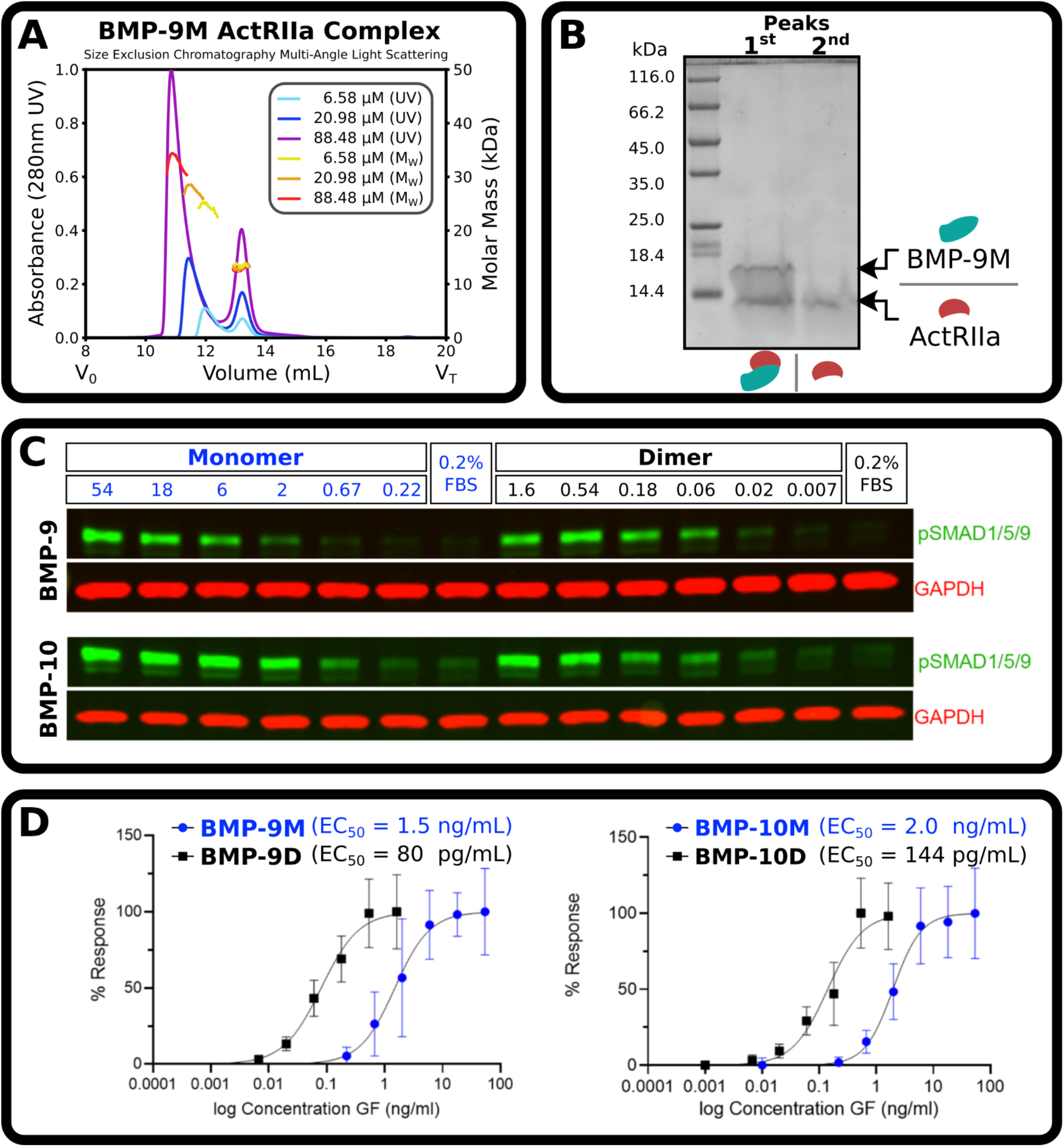
BMP-9 and BMP-10 monomers are solution monomers. **(A)** SEC-MALs of BMP-9 monomer with excess ActRIIa where BMP-9M was at 6.58 µM (UV, cyan; light-scattering, yellow), 20.98 µM (UV, blue; light-scattering, orange), and 88.48 µM (UV, purple; light-scattering, red). **(B)** SDS gel of the first and second peak of the 88.48 µM run. **(C)** Representative western blot, pSMAD1/5/9 and GAPDH. HUVECs were serum starved in 0.2% FBS for 4 hours, stimulated with indicated ligands for 45 minutes, and cell lysates separated by reducing SDS-PAGE and probed for pSMAD1/5/9 (green) and GAPDH (red). **(D)** Quantification of normalized pSMAD1/5/9 intensities from western blots, N = 3 individual experiments with two biological replicates per experiment.

Analysis of BMP-9 monomers by analytical ultracentrifugation (AUC) was performed to determine the noncovalent dimer self-dissociation constant (K_D_). Samples were at pH 3.8 to maintain solubility. However, at concentrations between 1.7 – 60.7 µM an enhanced van Holde-Weischet analysis (27) showed no appreciable shifts in the sedimentation coefficient (**Fig. S2A**). While a genetic algorithm – Monte Carlo (GA-MC) analysis (28,29) revealed the presence of both a monomeric and dimeric species in solution (**Fig. S2B**), mass action was also not observed under this low-pH solution condition. AUC-based hydrodynamic measurements are summarized in **Table S9**.

We sought to determine how the signaling activity of purified monomers compared to purified dimers. To this end, we treated serum-starved human umbilical vein endothelial cells (HUVECs) with ligands for 45 minutes and performed a western blot to measure phosphorylated (p)SMAD 1/5/9 (**Fig. 2C**). The half-maximal effective concentration (EC_50_) of the monomers, 1.5 ng mL^-1^ (124 pM) and 2.0 ng mL^-1^ (164 pM), was roughly 15 times higher by mass than dimers, 80 pg mL^-1^ (3.3 pM) and 144 pg mL^-1^ (5.9 pM), for both BMP-9 and -10, respectively (**Fig. 2D**). Decreased signaling activity of monomers compared to dimers is consistent with observations of solution monomers.

### BMP-9 and BMP-10 monomers are cysteinylated

The integrity of the purified monomers was verified by measuring the intact mass using electrospray ionization time of flight (ESI-TOF) mass spectrometry. The measured mass was 119 Da greater than expected in the absence of reductant but matched the expected mass when treated with tris (2-carboxyethyl) phosphine (TCEP), a reductant (**Fig. S3**). Since the covalent attachment of cysteine would be expected to increase the mass by 119 Da, we hypothesized that the monomers are cysteinylated. Of the seven cysteines in BMP-9 and -10, six form the three intrachain disulfides of the cystine knot that are essential for the folding of the monomer, so we further hypothesized that the cysteinylation resides on the interchain cysteine. If so, this might explain the poor electron densities around the interchain disulfide in several of the published BMP-9 and -10 structures determined from crystals formed from monomer-dimer mixtures (14,16,17,21,22).

The BMP-9 monomer was crystallized at neutral pH with a NaCl precipitant and solved at 1.90 Å resolution to observe the cysteinylation (**Table S1**). Monomers packed together in the same overall manner as previously reported for the BMP-9 dimer and strong density extended from the sulfhydryl group of the interchain cysteine, Cys-392 (**Fig. 3A**). Cysteine was readily modeled into this density, confirming our cysteinylation hypothesis (**Fig. 3A**). The carboxyl of the added cysteine forms a hydrogen-bond with the backbone amide of Cys-392 of the opposing monomer (**Fig. 3B**). Although supported by weaker density, the carboxyl also makes a charge-charge contact with the sidechain ammonium group of the opposing Lys-390 (**Fig. 3B**). In addition, the structure shows noncovalent dimers are not impaired by cysteinylation of Cys-392.

**Figure 3:**
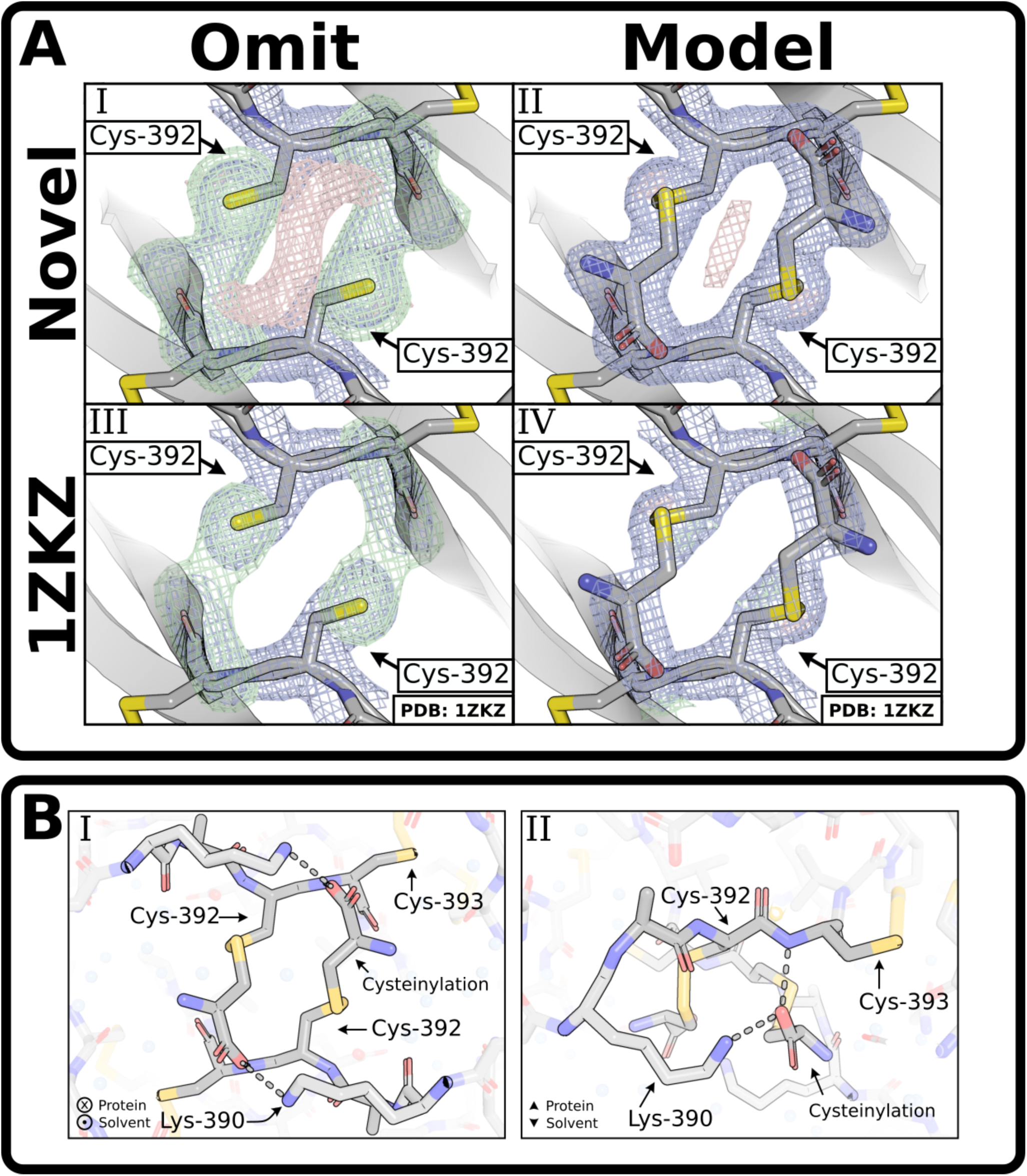
Crystallization of BMP-9 monomer shows cysteinylation. **(A)** Comparison of Cys-392 with **(A.I, A.III)** and without **(A.II, A.IV)** the cysteinylation modeled on both the our novel purified BMP-9 monomer **(A.I, A.II)** and a previously published (PDB: 1ZKZ) structure **(A.III, A.IV)**. In the modeled versions, the BMP-9 monomer structure reported here was modeled with 100% occupancy, while the previously published BMP-9 structure was modeled with 80% occupancy. **(B)** Model showing the cysteinylation’s contacts in dashed lines from perspectives looking into the protein **(B.I)** and along the protein edge **(B.II)**. For images showing density, the direct map is contoured at 1.5σ, and the difference map is contoured at 3.0σ.

Previously published BMP-9 structures were inspected to observe cysteinylation density derived from monomer-dimer mixtures. Due to partial occupancy and several water molecules in the vicinity of the interchain disulfide, the densities of all structures were initially unclear. For one of the structures, PDB: 1ZKZ (22), eliminating the modeled water molecules produced a clear omit map where the cysteinylation was readily modeled with ∼80% occupancy (**Fig. 3A**). Owing to the partial occupancy, the presence of cysteinylation is unclear in the other BMP-9 structures and cannot be reliably modeled.

### Radiation sensitivity of BMP-9’s interchain disulfide

Monomer cysteinylation raises the question of whether cysteinylation directly prevents formation of covalent dimers or is a consequence of the poor covalent dimerization leaving an exposed reactive cysteine. We reasoned that fully dimeric structures derived from pure dimer crystals would reveal the structural basis for inefficient covalent dimer formation. Hence, purified BMP-9 dimers were crystallized under neutral pH conditions similar to those used for the cysteinylated monomers. Contrary to expectations, the initial results revealed monomers with free cysteines rather than an intact interchain disulfide (**Fig. S4A**).

In an attempt to understand these results, the single-crystal dataset was divided into eight equal fractions over time, and it was found that the density of the interchain disulfide rapidly shifts from a monomer-dimer mixture entirely to monomer (**Fig. S4C-J**). The first fraction of this single-crystal analysis had too much monomer to model the dimer structure, so datasets were split into 8, 16, 32, or 64 parts, and the first fraction from nine crystals of BMP-9 dimer were combined. The interchain disulfide remains unbroken in the first fraction of the 32 and 64 splits, but is broken and the cysteines are free as non-cysteinylated monomers in the last fraction of all splits. In contrast, the intrachain disulfides lose some density, but do not change rotamer conformations (**Fig. 4A** and **Table S2**). Hence, there is highly localized radiation damage (i.e. radiation sensitivity) on the interchain disulfide, but not the intrachain disulfides.

**Figure 4:**
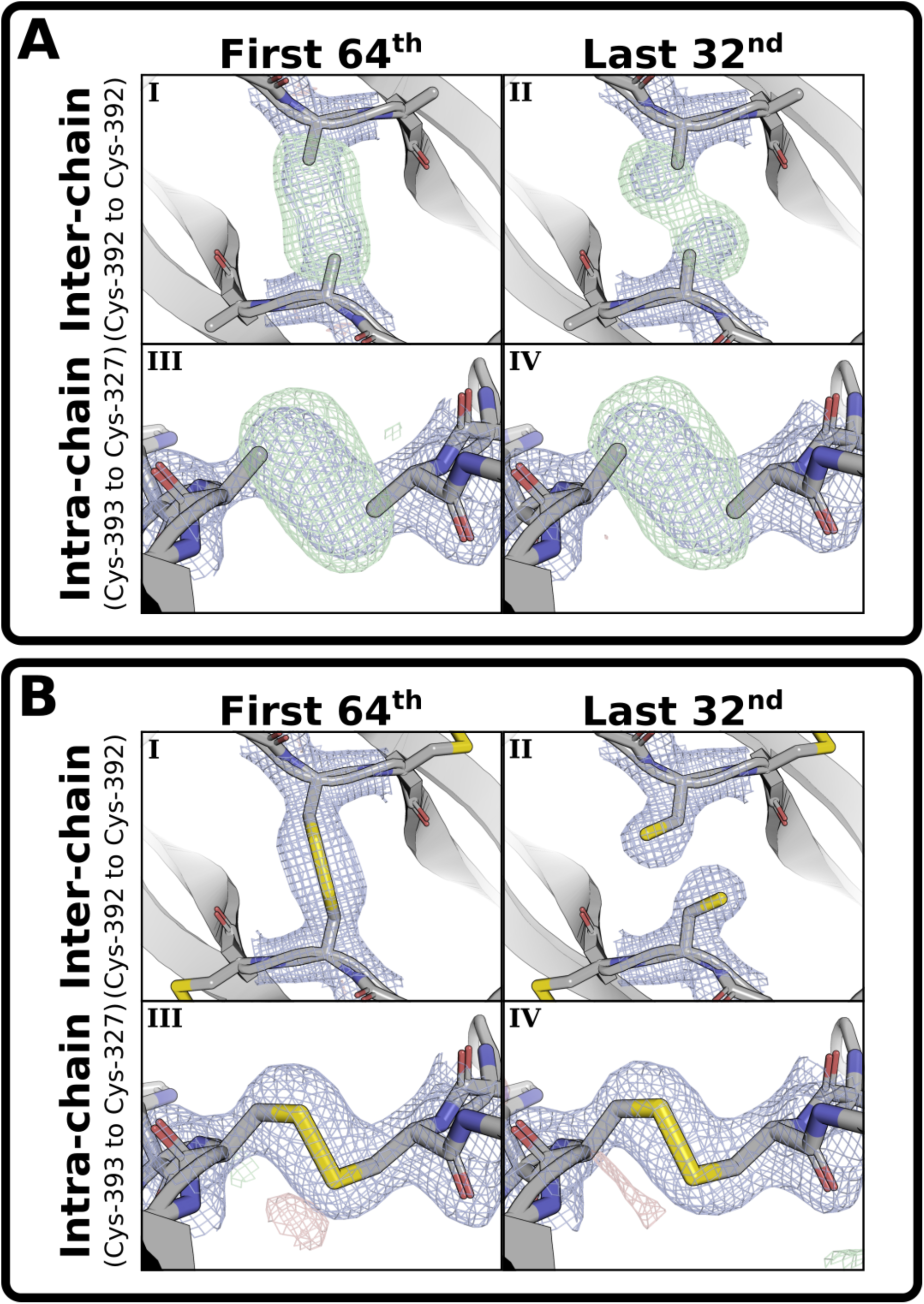
Radiation sensitivity of BMP-9’s interchain disulfide. **(A)** Sulfur-omit maps and **(B)** full models of the **(I,II)** interchain and **(III,IV)** intrachain disulfides for the **(I,III)** first 1/64 and **(II,IV)** last 32/32 blended fractions from nine datasets. The direct map is contoured at 1.5σ, and the difference map is contoured at 3.0σ.

Given the evidence of radiation sensitivity, we checked the reduction potential of BMP-9 and BMP-10 compared to activin-A (Act-A) by a reduction gradient and SDS-PAGE densitometry (**Fig. S6**). While the absolute reduction potential values are not consistent with published values, presumably due to some oxidation of the glutathione stocks, the interchain bond of BMP-9 and BMP-10 both have a more positive reduction potential than Act-A (**Fig. S6B,** legend). We checked for spontaneous monomerization (**Fig. S7**) or dimerization (**Fig. S8**) without reductants or oxidants. However, no evidence of spontaneous bond formation or breakage was found at 37 °C and neutral pH over a period of several days.

### Strain across BMP-9’s interchain disulfide

To model the intact interchain disulfide, we inspected the first fraction of the 32 and 64 splits. In the first fraction of the 32 split, automated fit functions placed the sulfurs in dimer density but well beyond disulfide bonding distance, and manual fit of the interchain cystine produces a perfect syn-periplanar (0°) χ_3_ dihedral angle (C_β_-S-S’-C_β_’), but it lacked full occupancy, both indicative of monomer contamination biasing the density. In the first fraction of the 64 split (**Fig. 4B.I**), automated fit functions still placed the sulfurs beyond disulfide bonding distance, but a manual fit to the omit-map density produces a perfect syn-periplanar (0°) χ_3_ dihedral angle (C_β_-S-S’-C_β_’) with full occupancy. This is illustrated in a tilt-video of the modeled interchain cystine in the omit density (see *Supplementary Material*).

To model the broken interchain disulfide, we inspected the last fraction of the 32 and 64 splits. While finer splitting produces less damage in early fractions, it produces more damage in later fractions. In the last fraction of the 64 split, global radiation damage increases the noise and decreases the resolution producing an overall worse structure with less reliable positioning of the interchain cysteine sulfurs and beta-carbons. In the last fraction of the 32 split (**Fig. 4B.II**), there is less damage and slightly higher resolution producing a better structure with more consistent and reliable positioning of the interchain cysteine’s sulfurs and beta-carbons.

For reference, we compared the dihedral angles of the dimer’s interchain disulfide to dihedral angles reported in the PDB and calculated torsion energies using the AMBER force field (30). While the observed χ_1_ dihedral angle of Cys-392 is both an uncommon and high energy conformation (**Fig. 5A.I**), the χ_3_ dihedral angle (C_β_-S-S’-C_β_’) of the Cys-392-Cys-392’ disulfide is an extreme outlier as the χ_3_ adopts a syn-periplanar conformation placing the C_β_ atoms directly across from each other in the same plane (**Fig. 5A.III**). The intrachain cystines have no outlier dihedral angles (**Table S8**) and all disulfide bond distances, both inter- and intrachain, were acceptable for this resolution. Other geometric parameters, such as cystine screw classification, are not representative of a strained cystine geometry and provide no additional insight (**Fig. S9**).

**Figure 5:**
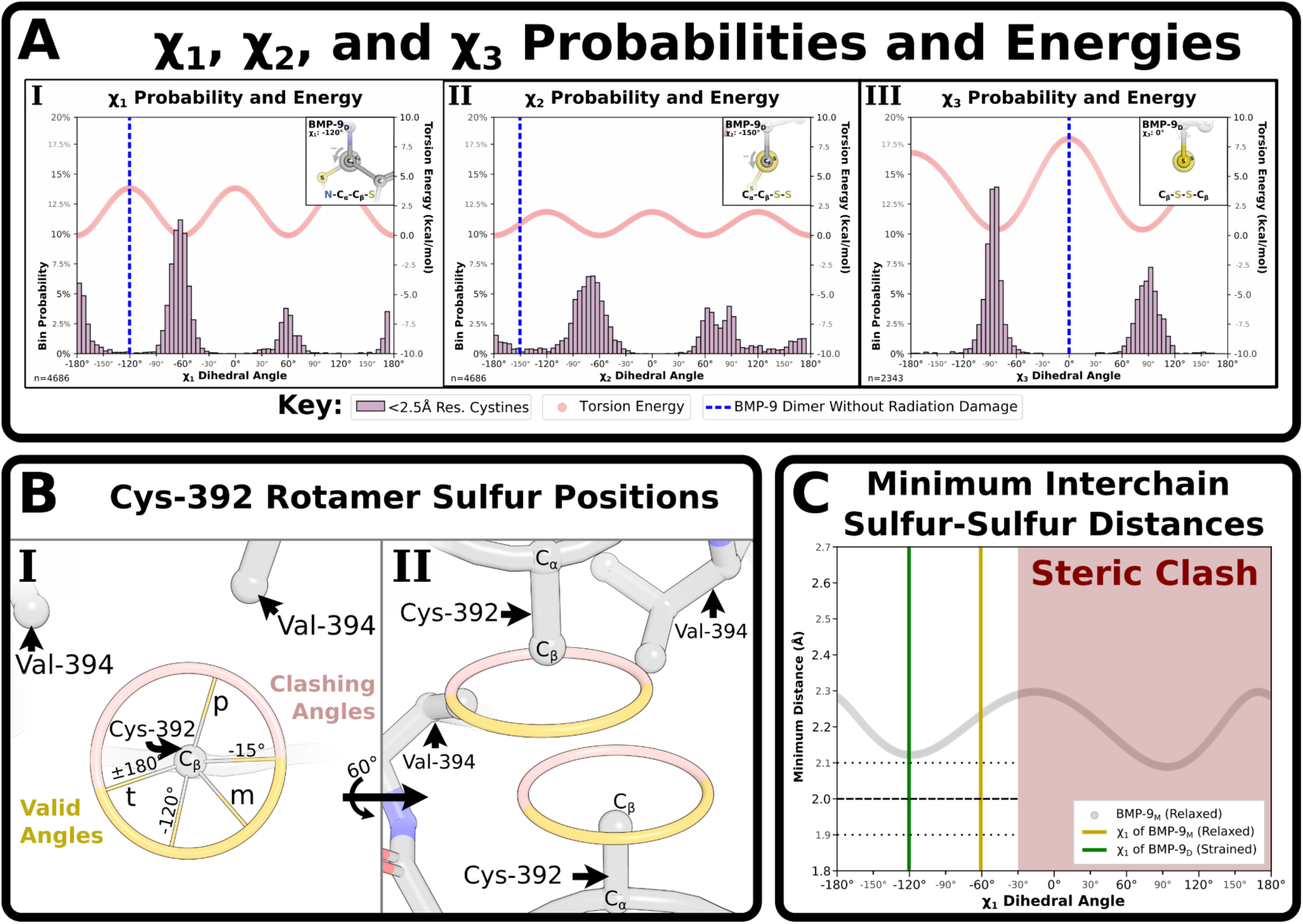
BMP-9 dimer’s Cys-392 dihedral conformations. **(A)** Plot of χ_1_ **(A.I)**, χ_2_ **(A.II)**, and χ_3_ **(A.III)** dihedral angles with frequency of angle in 2.5 Å resolution or better structures and Torsion energy as calculated from the Amber force field. Respective angle of BMP-9’s interchain bond plotted as vertical line and inset as Newman diagram. **(B.I)** Calculated possible sulfur positions in BMP-9’s Cys-392 depending on χ_1_ angles including the standard plus (p), minus (m), and trans (t) conformations, clashing angles in red, valid angles in yellow, edges of valid angles (-15° and ±180°), and the dimer angle -117°. **(B.II)** Calculated possible Cys-392 sulfur positions for each monomer in the dimer complex. **(C)** Calculated minimal possible interchain sulfur-sulfur distances.

### Formation of BMP-9’s interchain disulfide

Statistical mechanics predicts high energy conformations will have low occupancies. The interchain cystine of the BMP-9 dimer has one of the highest-energy conformations possible, (**Fig. 5A.III**), yet this is the dominant conformation prior to radiation damage. This is surprising unless no lower energy conformations are accessible. To better understand why this high energy conformation exists, the disulfide-broken (non-cysteinylated) monomer - as the most relaxed natural structure - was used to assess the conformations of the χ_1_ rotamer angle (N-C_α_-C_β_-S). First, steric clashes with Val-394 eliminate χ_1_ dihedral angles from -15° to 180° (**Fig. 5B.I**). This elimination includes both the plus (+60°) and trans (±180°) conformations, but the minus conformation (-60°) originally occupied structure remains. Next, all Cys-392 sulfur positions in each monomer were calculated (**Fig. 5B.II**). Finally, sulfur-sulfur distances for every opposing position were calculated and the minimum distance was plotted (**Fig. 5C**). All positions except the observed dimer conformation, χ_1_ = -120.3°, had minimum sulfur-sulfur distances greater than the maximum permissible disulfide bond distance (2.0 ± 0.1 Å). Hence, this explains both the high-energy dimer conformation and BMP-9’s poor propensity for covalent dimer formation. An extended analysis of the BMP-9 interchain cystine at neutral pH can be found in **Fig. S5**.

### Radiation sensitivity and strain in published structures

We sought to identify radiation sensitivity in previously published structures of BMP-9 and BMP-10. Only three previously published BMP-9 growth factor structures had sufficient resolution for comparison, PDB: 1ZKZ, 4MPL, and 5I05 (17,21,22). Two structures (PDB: 1ZKZ, 4MPL) lacked dimer density (**Fig. S10A,B**) and one structure (PDB: 5I05) had an equal monomer-dimer mixture (**Fig. S10C**). The first two structures were obtained from crystals grown at pH 7.4-7.5, while the third structure was obtained from crystals grown at pH 3.5. As such, we hypothesized low pH would disrupt the dimer interface, relieve strain, and prevent radiation sensitivity. To verify this hypothesis, pure BMP-9 dimer crystals were grown at pH 3.5. No radiation sensitivity was observed in the single-crystal analysis (**Fig. S10D**); however, mild radiation sensitivity was observed in the multi-crystal analysis (**Fig. S11A-H**). The densities could not be unambiguously modeled as strained or unstrained, thus, an unstrained model was used (**Fig. S11K**).

Most BMP-10 structures lack the resolution for a well-modeled interchain disulfide. Of the remaining structures, one crystallized at pH 5.8 (PDB: 7PPA) and the other co-crystallized with Alk1 bound on the dimer interface at pH 8.5 (PDB: 6SF3). As such, it is unsurprising that neither structure displays radiation sensitivity (**Fig. S12A,B**) or strain (**Fig. S12D-F**). Attempts were made to crystallize BMP-10 and obtain a high-resolution structure. While BMP-10 crystallized in many conditions, none of these crystals diffracted to high resolution. A chimeric BMP-10 with five residues (N357F, Y358F, A374T, Y409L, F411Y) replaced with BMP-9 crystal contacts was purified, crystallized, and solved. Unfortunately, the condition was at pH 6.2 and the crystal contacts disrupted the residues adjacent to the interchain cystine (**Fig. S12C)**, thus neither strain nor radiation sensitivity are observed in chimeric BMP-10. An extended analysis of the BMP-10 interchain cystine can be found in **Fig. S12**. A small sampling of related growth factors (TGF-β, Act-A, BMP-6, and BMP-2) published in the PDB (6XM2, 2ARV, 2QCW, and 2H64, see (31–34)) with neutral pH crystal conditions were inspected for radiation sensitivity, but none was observed (**Fig. S13**).

### Mutagenesis reveals dimerization residues

BMP-9 and BMP-10 growth factors, though not genes (35,36), are often assumed to be identical. However, SDS-PAGE after immobilized metal affinity chromatography (IMAC) purification consistently shows differences in monomer-dimer proportions (**Fig. 6C**). Densitometry shows roughly half as much dimer as monomer in BMP-9 but roughly equal proportions in BMP-10. Since the BMP-9 structural analyses we performed could not be replicated for BMP-10, the structural basis for these dimerization differences are unclear.

**Figure 6:**
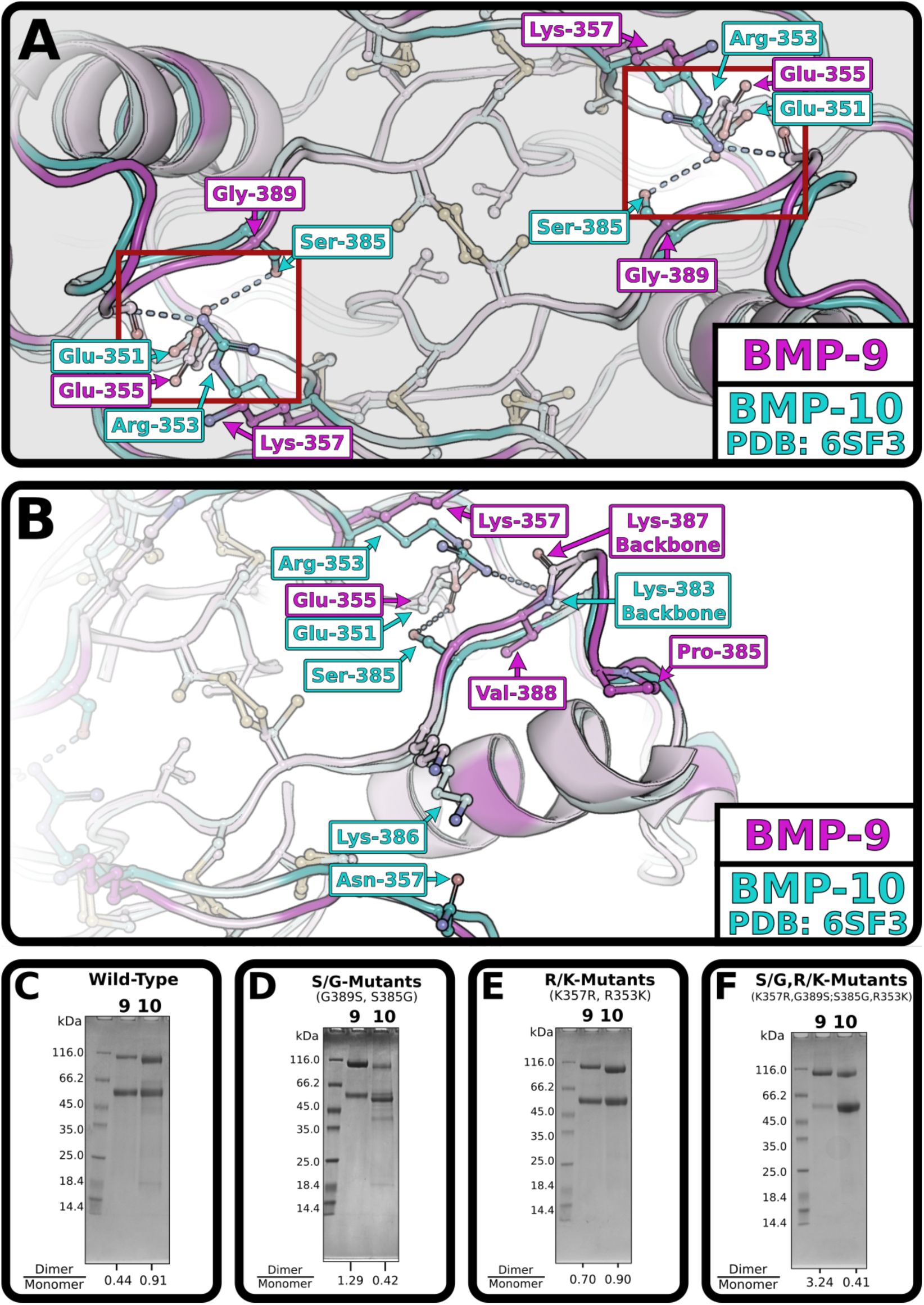
Mutations at interface influence dimerization. **(A,B)** Alignment of BMP-9 (magenta, novel) and BMP-10 (cyan, PDB: 6SF3) with hypothesized interactions, highlighted in the symmetric red boxes, darker residues are unshared in sequence alignment, lighter residues are shared in sequence alignment, shown as a direct view **(A)** or side view **(B)**. **(C, D, E, F)** SDS-PAGE of IMAC column purified Pro-BMP-9 and Pro-BMP-10 wild-type **(C)**, G389S BMP-9 and S385G BMP-10 **(D)**, K357R BMP-9 and R353K BMP-10 **(E)**, and G389S, K357R BMP-9 and S385G, R353K BMP-10 **(F)**. The dimer over monomer ratio for BMP-9 and BMP-10 respectively are shown at the bottom of the gel below the respective lane.

Sequence differences near the interchain cysteine were therefore examined for possible interchain contacts that might explain differences in disulfide bond formation (**Fig. 6A,B**). Two potential contacts present in BMP-10 and absent in BMP-9 were identified. Ser-385 in BMP-10 has its sidechain hydroxyl just beyond hydrogen bonding distance (3.8 Å) from an opposing glutamate’s oxygen (Glu-351 in BMP-10 or Glu-355 in BMP-9). Gly-389 in BMP-9’s equivalent position cannot form any contact. Arg-353 in BMP-10 has its sidechain guanidinium group 4.3 Å from the opposing backbone oxygen (Lys-383 in BMP-10 or Lys387 in BMP-9). Lys-357 in BMP-9’s equivalent position cannot use its ammonium group to form a similar interaction due to its shorter length. Notably, while neither of these contacts are in ideal hydrogen bonding distance in BMP-10, replacing the residues in some BMP-9 structures produces ideal hydrogen bonding distances (not shown). As such, we reasoned that Gly-389 and Ser-385, as well as Lys-357 and Arg-353 in BMP-9 and BMP-10, respectively, may be responsible for the dimerization differences observed by SDS-PAGE.

Mutagenesis was used to swap these dimerization residues, and SDS-PAGE densitometry was used to quantify the difference in dimer propensities. Swapping the identified serine/glycine residues is sufficient to swap dimer propensities (**Fig. 6D**). Swapping the identified lysine/arginine residues improves BMP-9 dimerization slightly but fails to impair BMP-10 dimerization (**Fig. 6E**). Similarly, the double mutants further improve BMP-9 dimerization but fail to further impair BMP-10 dimerization (**Fig. 6F**). These swaps suggest a role for these residues in facilitating dimerization.

### Crystal structures of BMP-9 mutants

To further understand how these residues contribute to growth factor dimerization, BMP-9 G389S and BMP-9 G389S, K357R dimers were crystallized at both neutral and low pH. The contacts in the resultant structures were compared to our BMP-9 and published BMP-10 structures (**Fig. 7A,B**). Where the serine is present, its sidechain hydroxyl is unstructured at neutral pH (**Fig. 7A**). At low pH, the serine contacts the opposing glutamate as predicted in BMP-10, but it is unclear from the omit maps whether the serine in the BMP-9 double-mutant has a water mediated contact to the opposing glutamate or directly contacts the mutated arginine (**Fig. 7B**). Where the arginine is present, the distal guanidino group sits between the serine and predicted backbone oxygen contact at the edge of accepted hydrogen bonding distances (**Fig. 7A,B**). Additionally, Lys-386 in BMP-10 is well structured due to a peripheral contact absent in BMP-9 **(Fig. S15A,B)**.

**Figure 7:**
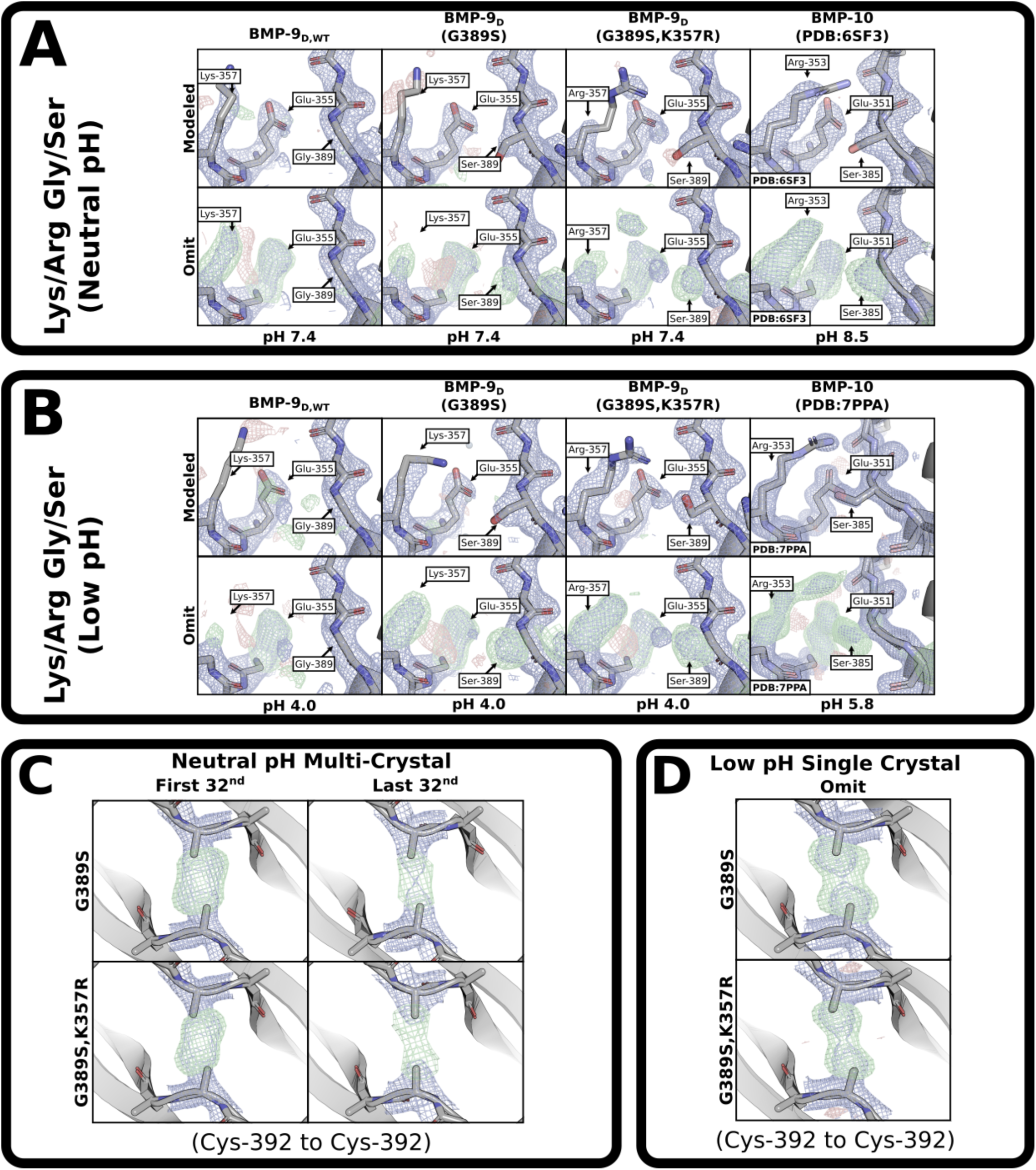
BMP-9 Mutants. **(A,B)** View of the lysine/arginine and glycine/serine dimerization residues in BMP-9 dimer wild-type, G389S, G389S K357R, and BMP-10 at **(A)** neutral pH and **(B)** low pH. Top rows are fully modeled, bottom rows are omits of relevant residues. **(C)** Multi-crystal analyses’ sulfur-omit density maps comparing undamaged (first 1/32) and damaged (last 32/32) BMP-9 G389S and BMP-9 G389S,K357R at neutral pH. **(D)** Single crystal analyses’ sulfur-omit density maps and modeled density maps for BMP-9 G389S and BMP-9 G389S,K357R at low pH. The direct maps are contoured at 1.5σ, and the difference maps are contoured at 3.0σ.

Since wild-type BMP-9 is radiation sensitive, the BMP-9 mutants were checked for radiation sensitivity. At neutral pH, the mutants lost density due to radiation damage but remained dimers (**Fig. 7C**). While neither mutant’s cystine could be unambiguously modeled in a strained conformation (**Fig. 7E**), the radiation sensitivity demonstrates that the bonds are not ideal. At low pH, both mutants were single crystal analyses (**Fig. 7D**). The mutants appeared as a monomer-dimer mixture, likely due to radiation sensitivity, thus the modeled conformations are unreliable (**Fig. 7D**). While it is difficult to make judgements on monomer-dimer mixtures, the low-pH mutants appear to trend towards more well-defined positions and decreased damage with each interaction. While the trend is not clear between mutants at neutral pH, the damage is decreased compared to wild-type. An extended analysis of the BMP-9 mutants’ interchain cystines can be found in **Fig. S16** and **Fig. S17** at neutral and low pH, respectively.

## Discussion

### Solution monomers permit monomer exchange

Our investigation into the BMP-9 and BMP-10 monomers by SEC-MALs revealed BMP-9, and by inference BMP-10, are solution monomers at micromolar and lower concentrations. This includes all biologically relevant concentrations and likely accounts for the reduced signaling activity of BMP-9 and BMP-10 monomers that we observe. This is potentially relevant to heterodimer formation as it shows that dimers can dissociate and exchange one monomer for another.

While mass spectrometry and crystallography show the monomers are cysteinylated, our investigation of dimer structures suggests cysteinylation is incidental. Regardless of whether the cysteinylation is present in animals, or an artifact of the medium used for cell culture, the cysteinylation likely occurs due only to the presence of a free cysteine. The cysteinylation itself serves no distinguishable function, and it is unnecessary for explaining monomers. Rather, the cysteinylation is an incidental consequence of the poor propensity for disulfide-bond formation.

### Strain induced radiation sensitivity

While an in-depth investigation of radiation damage is beyond the scope of this investigation, crystallography of the BMP-9 dimer shows localized radiation damage or radiation sensitivity of the interchain disulfide bond correspondent with extreme strain. There is prior evidence in the literature that suggests strain causes radiation sensitivity (37–41), though to our knowledge nothing is quite as dramatic as what we observe in BMP-9. Factors that could contribute to radiation sensitivity, such as radiation dose, cryoprotectant concentration, solvent exposure, and dynamics are not investigated here, obviating definitive conclusions. Nonetheless, BMP-9 may be an interesting system for studying highly localized radiation damage.

### Molecular basis of dimerization

Analysis of the BMP-9 dimer structure revealed a highly strained syn-periplanar interchain cystine prior to radiation damage, and an unstrained cysteine after the interchain disulfide was broken. Analysis of the unstrained structure by eliminating positions with steric clashes and invalid distances revealed the only accessible dimer is the strained syn-periplanar conformation. As such, BMP-9 homodimers require high-energy low-occupancy states, thus explaining the failure to dimerize. This explains the lack of dimer density in two previous BMP-9 structures with the third explained by reduced radiation damage at low pH, likely due to reduced strain. Unfortunately, an analogous analysis could not be performed on any BMP-10 structure due to both limited resolution and local disruptions (caused by pH, type-I receptors, or crystal contacts) in BMP-10 crystals. As such, the extended analyses of BMP-10 (**Fig. S12**) are limited to disulfide bonded cysteines in published structures, and it is not clear whether differences in these structures are due to deformations caused by the cystine bond, deformations in the overall structure, or fundamental differences in BMP-10.

### Dimerization differences

Our mutagenesis data demonstrated covalent homodimer formation differences between BMP-9 and BMP-10 due to glycine/serine and lysine/arginine residue differences near the interchain cystine. To the best of our knowledge, this observation is the only observed functional difference between the BMP-9 and BMP-10 growth factor domains.

The glycine/serine difference had a pronounced effect on the dimerization of both BMP-9 and BMP-10. Crystal structures with this serine vary by protein and condition with greater distances forming water-mediated interactions, mid-range distances likely forming transient contacts, and the smallest distances allowing for extended hydrogen bonds. It is notable that either an alternative glutamate position or a small shift in the overall interface may facilitate direct contact.

The lysine/arginine difference had a far greater effect on BMP-9 than BMP-10. Crystal structures with this arginine generally place it between the opposing serine and backbone. Notably, the loop with this backbone contact protrudes on BMP-9, but not BMP-10 (**Fig. 6B**). Consequently, the predicted backbone contact is slightly closer to the arginine in BMP-9. This structural asymmetry may explain the dimerization difference in mutagenesis and create the possibility of an asymmetric interface for BMP-9/10 heterodimers.

The interchain cystines on our mutant structures reveal decreased radiation sensitivity with each contact. This decreased damage implies a relief of strain on the interchain cystine bond that in turn explains the increased dimerization. While this decreased strain was not directly observable, the corresponding trend between decreased damage and increased dimerization does suggest the mechanistic relation. This may further imply a decreased strain in the BMP-9/10 heterodimer.

### Heterodimerization via registration shift

BMP-9/10 heterodimers are the predominant circulating form and signaling species in the blood (18,19), but the mechanism for heterodimer formation is not clear. Broadly, the mechanism could be some form of prodomain selection or growth factor complementarity. Based on the data presented here, we propose a registration shift as a form of growth factor complementarity to favor heterodimer formation. Specifically, we propose a shift of the monomers to accommodate a direct interaction between serine-385 on BMP-10 and glutamate-355 on BMP-9. As a result of this shift, the relative positions of the interchain cysteines would shift further apart to create a better geometry for the interchain bond assisted by the pre-existing displacement in BMP-10 (**Fig. S12**). Finally, this shift would further separate the arginine in its current conformation from its backbone contact (**Fig. 6B**), however the arginine can then straighten to contact the protruding loop seen in BMP-9 (caused by valine 388) and form a unique asymmetric contact. In the future, it will be important to determine the structure of the BMP-9/-10 heterodimer to verify these interactions promote heterodimer formation.

### Heterodimerization via mass-action

Regardless of whether some form of growth factor complementarity or registration shift occurs in the heterodimer, the data implies a preference for the BMP-9/10 heterodimer via mass action. Our SEC-MALs observations show that monomers in solution at biological concentrations are monomeric; thus, they are capable of monomer exchange. Our investigation of BMP-9’s interchain disulfide explains its failure to homodimerize; thus, there is an excess of BMP-9 monomers available. Our mutagenesis studies show why this excess of monomers is greater in BMP-9 than BMP-10. These facts together provide strong evidence for heterodimerization via mass-action anywhere BMP-9 and BMP-10 co-express, as in hepatic stellate cells.

## Conclusions

This research has thoroughly explored the molecular basis of interchain disulfide-bond formation in BMP-9 and BMP-10. The results have demonstrated the existence of cysteinylated solution monomers above biologically relevant concentrations, observed radiation sensitivity and strain on the interchain disulfide-bond, and identified dimerization residues responsible for growth factor complementarity. Our discussion extends these observations into an explanation of poor covalent homodimerization propensity, a new system to explore strain and radiation sensitivity, and a mechanistic hypothesis for BMP-9/10 heterodimer formation.

### Methods

### Gene design, cloning, and protein expression

All proteins were harvested five days after expression in suspension cultured HEK293 cells (expi293, Thermo Fisher Waltham, MA, USA) via the pcDNA3.1+ (Thermo Fisher Waltham, MA, USA) plasmid. Unless otherwise stated, coding sequences were obtained by gene synthesis (Twist Biosciences, San Francisco, CA, USA) and inserted downstream of the rat serum albumin signal peptide, a 8xhis-tag, and a factor Xa processing site (IEGR). All constructs were verified over their entire length by sequencing in the forward and reverse directions using primers that bound to the CMV promoter or the BGH terminator, respectively. See supplement for sequences.

*Pro-(fr)-TGF-β2*: Human proTGFβ2 (NCBI: NP_003229.1) residues 19-414 (C24S and P162A) downstream of the mouse Ig signal peptide, a 8xHis tag, and no processing site. The coding sequence was kindly provided by Dr. Peter Sun (NIAID, National Institutes of Health, Rockville, MD).

*Pro-(fr)-Act-A*: Human proINHBA (NCBI: NP_002183.1) residues 27-426 with no tag or processing site was inserted between BamHI and XbaI sites.

*Pro-(fr)-BMP-9* and *Pro-(X)-BMP-9*: Human proBMP9 (NCBI: NP_057288.1) residues 23-429 (A321S) was inserted between NheI and XhoI sites. In *Pro-(X)-BMP-9*, the natural furin processing site (RRKR) was replaced with a factor X processing site (IEGR).

*Pro-(fr)-BMP-10* and *Pro-(X)-BMP-10*: Human proBMP10 (NCBI: NP_055297.1) residue 22-424 was inserted between NheI and ApaI sites. In *Pro-(X)-BMP-10*, the natural furin processing site (AIRIRR) was replaced with a factor X processing site (GIEGR).

*Pro_9_-(X)-BMP-10 Crystal Chimera*: Human proBMP9 (NCBI: NP_057288.1) residues 23-315, a factor Xa cleavage site (IEGRDD), and Human proBMP10 (NCBI: NP_055297.1) residues 320-424 (N357F, Y358F, A374T, Y409L, and F411Y) was inserted between NheI and XhoI sites.

*Albumin-ActRIIb*: Human Albumin (NCBI: NP_000468.1) with mutation V605A and a 6xHis tag inserted after the natural signal peptide, a disordered linker containing a thrombin cut-site (GSTSGSGAQTNASGT ***LVPRGS*** HMLEDPVP), the mouse ActRIIb (NCBI: NP_001300686.1) residues 25-117.

*ActRIIa*: Human activin receptor type-2A (NCBI: NP_001265508.1) residues 23-123 and a disordered linker containing a thrombin cut-site (SSG**LVPRGS**HM) in place of our standard factor Xa cleavage site was inserted between NheI and XhoI sites.

*Mutants*. The mutants were based on *Pro-(X)-BMP-9* and *Pro-(X)-BMP-10*. Mutagenesis was performed with oligos obtained through Integrated DNA Technologies (IDT). The oligo sequences can be found in the supplemental.

### Protein purification

*Pro-BMP-9 or Pro-BMP-10 monomer-dimer mixtures*. After expression, media was concentrated to about 250 mL then dialyzed twice into 4 L of 150 mM NaCl, 8 mM imidazole, 25 mM phosphate, pH 8. Sample was loaded onto an iminodiacetic acid (IDA) resin nickel column and eluted over a 300 mL gradient to 0.5 M imidazole.

*BMP-9 or BMP-10 monomers and dimers*. The monomer-dimer mixture (described above) is dialyzed into 20 mM tris pH 8, 100 mM NaCl, 0.5 mM CaCl, 0.02% Na azide. Activated factor-X (factor-Xa) is added at a ratio of 1:1000 by mass and digested at 37°C over the weekend (∼2 days). The digested sample is dialyzed into 4 L of 4 mM HCl. The supernatant is loaded on a C18 reverse phase column and eluted over a 400 mL gradient from 30%-40% of water with 0.1% trifluoroacetic acid (TFA) versus acetonitrile with 0.1% TFA. The resulting fractions are lyophilized and resuspended in 50 mM acetic acid.

*Pro-BMP-9 or Pro-BMP-10 monomers and dimers*. The monomer-dimer mixture (described above) is dialyzed into 150 mM NaCl 25 mM tris pH 8, concentrated, then 8 M urea 0.5 M NaCl 25 mM tris pH 8 is added to the sample. The sample is run over a HiLoad 26/60 Superdex 200 in 8 M urea 0.5 M NaCl 25 mM tris pH 8 with the respective monomer or dimer peak separated until pure. The final purified monomer or dimer complex is dialyzed into 150 mM NaCl 25 mM tris pH 8.

*Albumin-ActRIIb*: After a nickel column purification (see monomer-dimer mixture), the protein was further cleaned by size exclusion chromatography (HiLoad 26/60 Superdex 200 column, Cytiva) in 150 mM NaCl, 50 mM tris pH 8.

*ActRIIa*: After a nickel column purification (see monomer-dimer mixture), the protein deglycosylated in a 14 hr incubation with PNGAseF at 30°C then cleaned by size exclusion chromatography (HiLoad 26/60 Superdex 75 column, Cytiva) in 150 mM NaCl, 50 mM tris pH 8.

### SDS-PAGE

Sodium dodecyl sulfate polyacrylamide gel electrophoresis (SDS-PAGE) was performed with either gradient gels purchased from GenScript (**Fig. 1 and 6**) or homemade 12% acrylamide gels (all other figures). The homemade 12% gels were made with 11.6% acrylamide, 0.4% bis-acrylamide, 1 M tris pH 8.5, and 10 mM SDS as well as ammonium persulfate (APS) and tetramethylethylenediamine (TEMED) to catalyze the polymerization. All protein markers used were the ThermoFisher Scientific Pierce unstained protein MW marker.

*SDS-PAGE for reduction potential gradients*. Each reductant condition was prepared with 50 mM HEPEs pH 7.5 buffer, 1 mM oxidized glutathione, and a calculated proportion of reduced glutathione for the reduction potentials (-75 mV to -300 mV). 50 uL solutions with 4.93 μM of Alb-ActRIIb and 3 μg, or 2 μg for denatured conditions, of growth factor were prepared. Each condition was allowed to react overnight (∼16 hrs) before 30 uL of 2x running stain (0.81 M tris, 31.6% glycerol, 0.072% SDS, 0.0135% coomassie blue, 0.0045% phenol red) was added. For the denatured conditions, the samples with SDS were placed at 50**°**C for 45 min. Gels were run cooled to 6°C at 80 V over roughly 4 hrs.

*Densitometry analysis*. Quantification of SDS bands was performed with ImageJ by plotting each lane with the uncalibrated optical density conversion setting after processing with the subtract background function. Taking the area under the plotted curve for each band, different monomer forms were summed together (where reduced forms were present) and either a ratio of monomer dimer or fraction of the total summed mass was calculated.

### Size exclusion chromatography multi-angle light scattering

Protein complexes for size exclusion chromatography multi-angle light scattering (SEC-MALS) were prepared from concentrated stocks to 100 μL of 1:1.6 BMP-9M:ActRIIa complexes at the tested concentrations (6.58 µM, 20.98 µM, and 88.48 µM) of BMP-9 Monomer. The running buffer was 150 mM NaCl, 100 mM tris pH 7.5, and the column was a Superdex 75 Increase 10/300 GL column (GE Lifesciences, Piscataway, NJ, USA). SEC-MALS measurements were made using a Waters high-performance liquid chromatography system (Waters, Milford, MA) and a Wyatt DAWN HELEOS-II multiangle light scattering detector and Optilab T-rEX refractive index detector (Wyatt, Santa Barbara, CA, USA). SEC-MALS instrument control was performed with the ASTRA software package (Wyatt, Santa Barbara, CA, USA). UV is normalized globally in ASTRA EASI Graph. Final plot was done in matplotlib.

### Analytical ultracentrifugation

All experiments were performed at the Canadian Center for Hydrodynamics at the University of Lethbridge using a Beckman-Coulter Life Sciences Optima AUC instrument. Cysteinylated BMP-9 was diluted in 15 mM NaPO4, pH 3.8, 100 mM NaCl, and loaded into 2-channel epon centerpieces, fitted with quartz windows. Sedimentation velocity experiments were performed at 50 krpm in an AN50Ti rotor at 20°C. Data was collected for 12.7 hours. All experiments were measured in intensity mode, and analyzed by UltraScan-III (release 7179) (42) according to methods and workflows described in (43).

### Signaling assays

Human umbilical vein endothelial cells (HUVECs, Promocell, Heidelberg, Germany) were cultured in endothelial growth medium 2 (EGM2) with 2% FBS and supplied growth factors. For stimulations, HUVECs (passage 6) were plated in 6-well plates and grown to 90% confluence over 2 days. HUVECs were serum starved in endothelial cell basal medium (Promocell) supplemented with 0.2% FBS for 4 hours prior to treatment with BMP-9 and BMP-10 ligands (monomers: 0.01-54 ng/mL; dimers, 0.001-1.62 ng/mL). Cells were collected at 45 minutes, lysed in RIPA buffer supplemented with HALT protease inhibitor (Thermo Fisher Scientific, Waltham, MA, USA), and frozen at -80°C. Just before use, samples were thawed, sonicated and centrifuged. The cleared supernatants were collected, and protein concentrations were determined using the Pierce BCA Protein Assay (Thermo Fisher Scientific). Ten micrograms of protein were separated by 10% reducing SDS-PAGE and transferred to nitrocellulose membrane (Bio-Rad, Hercules, CA, USA). Membranes were dried for 1 hour, rehydrated in water, and blocked in Intercept (TBS) Blocking Buffer (LI-COR, Lincoln, NE, USA) for 1 hour. Antibodies were diluted in blocking buffer with 0.1% Tween 20. Membranes were probed overnight at 4°C with a 1:1000 dilution of rabbit phospho-SMAD 1/5/9 antibody (#13820, Cell Signaling Technology, Danvers, MA, USA), followed by a 1:10,000 dilution of mouse GAPDH antibody (#ab8245, Abcam, Waltham, MA, USA) for 1 hour at room temperature the next day. Membranes were washed in TBS-Tween and probed with 1:12,000 dilutions of IRDye 800CW donkey anti-rabbit IgG (#925-32213, LI-COR) and IRDye 680LT goat anti-mouse IgG (#926-68020, LI-COR) secondary antibodies for 1 h at room temperature, protected from light. Membranes were washed with TBS-Tween while protected from light and imaged using the Odyssey CLx Imaging System (LI-COR). pSMAD1/5/9 intensities were measured using Image Studio 5.2 software (LI-COR) and normalized to GAPDH, according to manufacturer’s instructions. Data were plotted as log concentration versus mean % maximal response and fit using nonlinear regression with 4-parameter variable slope (Prism 8, GraphPad, San Diego, CA, USA). N = 3 individual experiments with 2 replicates per experiment.

### Mass spectrometry

When probing cysteinylation, purified BMP-9 or BMP-10 monomers were either reduced with TCEP (tris(2-carboxyethyl)phosphine) or not reduced then diluted to 10 μM in mass-spec running buffer (5% acetonitrile,0.01% trifluoroacetic acid, 0.1% formic acid). Each sample was run on a Bruker mass spectrometer instrument and processed with the associated software by restricting the entropy to the 10 kDa to 15 kDa mass range.

### Crystallography

#### Crystallization, integration, and reduction of BMP-10 crystal chimera

Automated screening for crystallization was carried out using the sitting drop vapor-diffusion method with an Art Robbins Instruments Phoenix system in the Structural Biology Core at the University of Texas Health Science Center at San Antonio. Crystals of the BMP-10 Crystal Chimera formed at room temperature in a one-to-one mixture of 5.0 mg mL^-1^ protein and 25% (v/v) 1,2-propanediol, 100 mM sodium phosphate dibasic/potassium phosphate monobasic pH 6.2, 10% (v/v) glycerol (Rigaku Wizard Cryos E11). Crystals were flash-cooled in liquid nitrogen by wicking off excess solution from crystals harvested in nylon cryo-loops and screened on the Structural Biology Core home source X-ray generator. Additional data were acquired at the Advanced Photon Source NE-CAT beamline 24-ID-E (Argonne, IL). Diffraction data were processed using XDS (44). The structure was determined by the molecular replacement method implemented in PHASER (45) using coordinates from PDB entry 6SF3 as the search model. Coordinates were refined using PHENIX (46) including simulated annealing and alternated with manual rebuilding using COOT (47,48). The model was verified using composite omit map analysis. Data collection and refinement statistics are shown in Table S4.

#### Crystallizations of BMP-9

All BMP-9 crystals were formed using hanging drop crystallization in 24-well plates with 300 μL of well-solution and siliconized glass cover slips. Crystals formed within 4 days at 16 °C in drops prepared by mixing 0.4 μL of 5 mg mL^-1^ Protein and 0.4 μL of well-solution. When looped they were transferred to a drop of well-solution adjusted with cryoprotectant, if listed. For the *BMP-9 monomer*, the well solution was 0.9 M NaCl, 133 mM HEPES, 166 mM MES pH 7 and the cryoprotectant was 15% glycerol. For *BMP-9 dimer at neutral pH*, the well solution was 1 M NaCl, 17% glycerol, 0.1 M HEPEs pH 7.4 and the cryoprotectants ranged across 10-15% PEG200 or 10-15% glycerol. For *BMP-9 dimer at acidic pH*, the well solution was 1 M NaCl, 3.5% PEG8K, 0.3 M sodium citrate pH 3.5 and the cryoprotectants ranged across 15-30% glycerol. For *BMP-9 dimer mutants at neutral pH*, the well solution was 1 M NaCl, 22-26% glycerol, 0.1 M HEPEs pH 7.4 and the cryoprotectants ranged across 25-35% glycerol. For *BMP-9 dimer mutants at acidic pH*, the well solution was 1 M NaCl, 0.1 M acetic acid pH 4, with 27% or 32% glycerol for the single and double mutant, respectively. No additional cryoprotectant. Each crystal was mounted in nylon loops. Excess well solution was wicked off and the looped crystals were flash-frozen in liquid nitrogen before being shipped at liquid nitrogen temperature for remote collection.

#### Integration, reduction, and phasing of BMP-9

The diffraction data from the BMP-9 monomer structure was collected at the Stanford Synchrotron Radiation Lightsource (SSRL) BL12-2 beamline then integration and reduction were done in the HKL-3000 suite (49) before conversion to the mtz format. All other BMP-9 structures were collected at the Southeast Regional Collaborative Access Team (SER-CAT) 22-ID beamline at the Advanced Photon Source, Argonne National Laboratory then integration was done with *iMosflm* (50). All BMP-9 crystals crystallized in the I4_1_22 spacegroup as confirmed via *pointless* (51,52). For the crystals where the datasets were split, the integrated diffraction data was split using *rebatch* in the CCP4 suite to take either the first or last 32^nd^ (or first 64^th^ for wild type BMP-9) of diffraction images from each crystal’s 360° datasets. That is the first or last 11.25° collected over 6.75 s or 13.5 s (varied) at 20% transmission with a ring current of 102 mA. *Blend* (53) was used to combine the first and last datasets. Except for the BMP-9 monomer, the data was reduced with *aimless* (54), *ctruncate* (55–59), and the *uniquify* (60) script in the CCP4 software suite (61). For all BMP-9 crystals, *Molrep* (62) was used for molecular replacement using PDB entry 1ZKZ or the structure determined for the first fraction (1/32nd). Several cycles of refinement using phenix refine (46) and model building using *COOT* (47,48) were performed to determine the final structure. Data collection and refinement statistics are shown in **Table S1-S6**.

#### Omit maps and images

All non-omit structures were used as determined or downloaded from the PDB and all direct maps were generated with downloaded phases in *refmac5* (63–70). Omit maps were generated by modifying structures in COOT (47,48) as described then re-calculating phases with zero-step refinements in *refmac5* (63–70). Images of protein structures were generated using Open-Source PyMOL (71). The direct maps (2mF_Obs_ - DF_Cal_) are contoured at 1.5σ, and the difference maps (mF_Obs_ - DF_Cal_) are contoured at 3.0σ.

### Calculations

#### Histograms of dihedral angle distributions

A list of PDB entries was generated from “Cystine-Knot Cytokine” Superfamily (IPR029034), “Snake Toxin-Like” Superfamily (IPR045860), and Zona Pellucida Domains (IPR042235) with resolutions better than 2.5 Å that did not contain BMP-9 or BMP-10. A simple python script was used to compile a list of cystine dihedral angles from those structures. Free cysteines and geometries not immediately processable by PyMOL (non-bonding atoms, too many bound atoms, etc) were filtered out. Standard definitions for each dihedral angle were used: χ_1_ (N-C_α_-C_β_-S), χ_2_ (C_α_-C_β_-S-S’), χ_3_ (C_β_-S-S’-C’_β_). This resulted in 6898 cystines without a resolution filter and 2343 cystines with the 2.5 Å resolution filter. The histograms were binned between -180° and 180° with a 5° bin width for the dihedral angles. The y-axes were normalized. The matplotlib module was used to generate the graphs.

#### Torsion energy calculations

For simplicity’s sake, the original AMBER force field (30) parameters for torsion energy were used to calculate torsion energies of each respective χ dihedral angle. The equation for each disulfide dihedral is included below. The matplotlib module was used to generate the graphs.

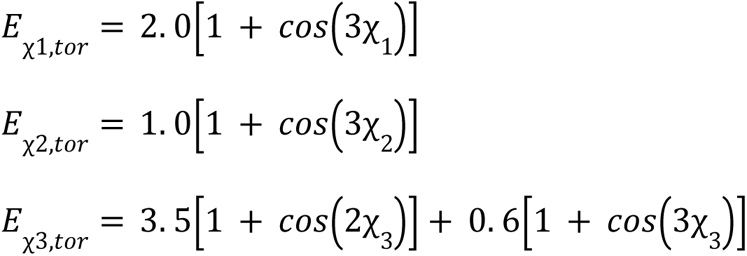

#### Minimum sulfur-sulfur distances calculations

For each χ_1_ dihedral angle a radial and height vector is used to calculate the sulfur position. The radial vector was determined via vectors perpendicular to C_α_-C_β_ and the projection of C_β_-S then combined as linear combinations with the sine and cosine functions all scaled by the C_β_-S distance (1.81 Å) and the sine of π minus the C_α_-C_β_-S angle (114.3°). The height vector was determined by the unit vector of C_α_-C_β_ scaled by the C_β_-S distance (1.81 Å) and the cosine of π minus the C_α_-C_β_-S angle (114.3°).

Circles of possible conformations of each Cys-392 sulfur were calculated in one-degree increments and plotted as pseudoatoms in Open-Source PyMOL (71). Inspection in COOT (47,48) via molprobity (72) was used to identify clashing conformations. For each angle a list of distances to the opposing sulfur at every angle was determined and the minimum of that list was plotted along with vertical markers for the sulfur-sulfur distances. The matplotlib python module was used to generate the graphs.

## Accession numbers

PDB: 9DPM, 9DPN, 9DPO, 9DPP, 9DPQ, 9DPR, 9DPS, 9DPT, 9DPU, 9DPV, 9DPW, 9DPX, 9DPY

## Acknowledgments

We would like to thank Bill Furey for helpful discussions about localized radiation damage, Aina Cohen for helpful suggestions on X-ray data collection and analysis, and Christine Monnie for assistance with the SEC-MALS measurements. This research was supported by the U.S. Department of Defense (FPA W81XWH-17-1-0429 awarded to A.P.H and B.L.R), the NIH (R01 HL133009, R01 HL136566 awarded to B.L.R and R01 GM120600 awarded to B.D.), the Canada 150 Research Chairs Program (C150-2017-00015 awarded to B.D), the Canada Foundation for Innovation (CFI-37589 awarded to B.D.), and NSERC Canada (DG-RGPIN-2019-05637 awarded to B.D.). X-ray diffraction data was collected at the Southeast Regional Collaborative Access Team (SER-CAT) 22-ID and Northeastern Collaborative Access Team (NE-CAT) 24-ID-E beamlines at the Advanced Photon Source, beamline BL12-2 at the Stanford Synchrotron Radiation Lightsource (SSRL) at the SLAC National Accelerator Laboratory, and the Structural Biology Core (SBC), a part of the Institutional Research Cores at the University of Texas Health Science Center at San Antonio (UTHSCSA). SER-CAT is supported by its member institutions and equipment grants (RR25528, RR028976 and OD027000) from the NIH. NE-CAT is supported by Center of Excellence and Equipment grants (GM124165 and S10 OD021527) from the NIH. The research at SER-CAT and NE-CAT used the resources of the Advanced Photon Source, a U.S. Department of Energy (DOE) Office of Science User Facility operated for the DOE Office of Science by Argonne National Laboratory under Contract No. DE-AC02-06CH11357. SSRL is supported by the U.S. Department of Energy, Office of Science, Office of Basic Energy Sciences under Contract No. DE-AC02-76SF00515. The SSRL Structural Molecular Biology Program is supported by the DOE Office of Biological and Environmental Research, and by the National Institutes of Health, National Institute of General Medical Sciences (P30GM133894). UTHSCSA SBC is supported by the Office of the Vice President for Research, Greehey Children’s Cancer Research Institute, the Mays Cancer Center Drug Discovery and Structural Biology Shared Resource (NIH P30 CA054174), and NIH S10 OD030374 (awarded to A.B.T.).

## Notes

### Competing Interest Statement

The authors have declared no competing interest.

