## supplemental data for "Molecular basis of interchain disulfide-bond formation in BMP-9 and BMP-10"

### Table S1

| PDB | BMP-9 WT Monomer, Cysteinylated 9DPM |
|---|---|
| **Data Collection** | |
| X-Ray Source | SSRL BL12-2 |
| Wavelength (Å) | 0.9798 |
| Detector | DECTRIS EIGER2 XE 16M |
| Linear Cell Variation | N/A |
| Number of Crystals | 1 |
| **Data Reduction** | |
| Spacegroup | I $4_1$ 2 2 (98) |
| a, b, c (Å) | 71.86, 71.86, 144.61 |
| α. β, γ (°) | 90.00, 90.00, 90.00 |
| Completeness (%) | 100.0 (99.9) |
| Resolution (Å) | 29.46-1.90 (1.94-1.90) |
| $R_{meas}$ | 0.139 (3.093) |
| $R_{pim}$ | 0.028 (0.643) |
| $\langle I/\sigma(I) \rangle$ | 11.6 (0.7) |
| Redundancy or Multiplicity | 23.78 |
| $CC_{1/2}$ | 0.998 (0.598) |
| CC* | 0.999 (0.865) |
| $\chi^2$ | 0.687 (0.544) |
| Proteins/ASU | 1 |
| Percent Solvent (%) | 68.3 |
| Matthews Coefficient | 3.88 |
| Wilson B-Factor (Å$^2$) | 39.23 |
| No. Observations | 366054 |
| No. Reflections | 15391 |
| **Refinement** | |
| Resolution (Å) | 29.46 (1.90) |
| $R_{Work}$ / $R_{Free}$ | 0.2046 / 0.2417 |
| R.M.S. Deviations | |
| Bond-Lengths (Å) | 0.0076 |
| Bond-Angles (°) | 1.4093 |
| **Numbers** | |
| Residues | 107 |
| Atoms / Non-Hydrogens | 1767 / 936 |
| Protein: Overall / Heavy / Backbone | 1673 / 842 / 430 |
| Ions and Ligands: Overall / Heavy | 6 / 6 |
| Waters | 88 |
| B-factors | |
| Protein: Overall / Heavy / Backbone | 73.29 / 66.15 / 60.89 |
| Ions and Ligands: Overall / Heavy | 58.77 / 58.77 |
| Waters | 57.91 |
| **MolProbity** | |
| Clash Score | 0.00 |
| MolProbity Score | 0.52 |
| Ramachandran | |
| Allowed | 100 % |
| Favored | 99.04 % |
| Outliers | 0 % |
| Rotamers | |
| Allowed | 98.95 % |
| Favored | 98.95 % |
| Outliers | 1.05 % |
| Clashes | 0 |

### Table S2:

| PDB | BMP-9 WT Dimer, First 64th 9DPN | BMP-9 WT Dimer, Last 32nd 9DPO |
|---|---|---|
| **Data Collection** | | |
| X-Ray Source | APS 22-ID | APS 22-ID |
| Wavelength (Å) | 1 | 1 |
| Detector | DECTRIS EIGER X 16M | DECTRIS EIGER X 16M |
| Linear Cell Variation | 0.91 % | 0.91 % |
| Number of Crystals | 9 | 9 |
| **Data Reduction** | | |
| Spacegroup | I $4_1$ 2 2 (98) | I $4_1$ 2 2 (98) |
| a, b, c (Å) | 71.20, 71.20, 146.54 | 71.20, 71.20, 146.52 |
| α. β, γ (°) | 90.00, 90.00, 90.00 | 90.00, 90.00, 90.00 |
| Completeness (%) | 95.9 (96.7) | 99.8 (100.0) |
| Resolution (Å) | 41.50-2.24 (2.31-2.24) | 41.49-2.34 (2.42-2.34) |
| $R_{meas}$ | 0.198 (1.058) | 0.206 (1.305) |
| $R_{pim}$ | 0.103 (0.558) | 0.077 (0.478) |
| $\langle I/\sigma(I) \rangle$ | 4.3 (1.6) | 6.0 (1.8) |
| Redundancy or Multiplicity | 3.2 (3.3) | 7.1 (7.3) |
| $CC_{1/2}$ | 0.970 (0.471) | 0.984 (0.638) |
| CC* | 0.992 (0.8) | 0.996 (0.883) |
| $\chi^2$ | 0.93 (0.85) | 1.01 (1.02) |
| Proteins/ASU | 1 | 1 |
| Percent Solvent (%) | 68.7 | 68.7 |
| Matthews Coefficient | 3.93 | 3.93 |
| Wilson B-Factor (Å$^2$) | 36.29 | 43.21 |
| No. Observations | 28789 (2646) | 58722 (5794) |
| No. Reflections | 8929 (800) | 8300 (796) |
| **Refinement** | | |
| Resolution (Å) | 41.50 (2.240) | 41.528 (2.340) |
| $R_{Work}$ / $R_{Free}$ | 0.1953 / 0.2061 | 0.1989 / 0.2446 |
| R.M.S. Deviations | | |
| Bond-Lengths (Å) | 0.0093 | 0.0073 |
| Bond-Angles (°) | 1.3532 | 1.3381 |
| **Numbers** | | |
| Residues | 105 | 105 |
| Atoms / Non-Hydrogens | 1727 / 908 | 1715 / 894 |
| Protein: Overall / Heavy / Backbone | 1625 / 822 / 420 | 1627 / 822 / 420 |
| Ions and Ligands: Overall / Heavy | 37 / 21 | 37 / 21 |
| Waters | 65 | 51 |
| B-factors | | |
| Protein: Overall / Heavy / Backbone | 61.54 / 55.45 / 51.04 | 66.58 / 60.24 / 55.23 |
| Ions and Ligands: Overall / Heavy | 48.54 / 49.09 | 51.99 / 52.81 |
| Waters | 55.82 | 57.15 |
| **MolProbity** | | |
| Clash Score | 0.00 | 0.00 |
| MolProbity Score | 0.53 | 0.53 |
| Ramachandran | | |
| Allowed | 99.03 % | 100 % |
| Favored | 99.03 % | 99.03 % |
| Outliers | 0.97 % | 0 % |
| Rotamers | | |
| Allowed | 98.91 % | 98.91 % |
| Favored | 98.91 % | 97.83 % |
| Outliers | 1.09 % | 1.09 % |
| Clashes | 0 | 0 |

**Table S3:**

| PDB | BMP-9 WT Dimer, Acidic 9DPP | BMP-9 WT Dimer, Acid, First 32nd 9DPQ | BMP-9 WT Dimer, Acid, Last 32nd 9DPR |
|---|---|---|---|
| **Data Collection** | | | |
| X-Ray Source | APS 22-ID | APS 22-ID | APS 22-ID |
| Wavelength (Å) | 1 | 1 | 1 |
| Detector | DECTRIS EIGER X 16M | DECTRIS EIGER X 16M | DECTRIS EIGER X 16M |
| Linear Cell Variation | N/A | 0.55% | 0.55% |
| Number of Crystals | 1 | 7 | 7 |
| **Data Reduction** | | | |
| Spacegroup | $I\,4_1\,2\,2$ (98) | $I\,4_1\,2\,2$ (98) | $I\,4_1\,2\,2$ (98) |
| a, b, c (Å) | 70.94, 70.94, 145.86 | 71.04, 71.04, 145.57 | 71.04, 71.04, 145.56 |
| α. β, γ (°) | 90.00, 90.00, 90.00 | 90.00, 90.00, 90.00 | 90.00, 90.00, 90.00 |
| Completeness (%) | 100.0 (100.0) | 97.9 (98.3) | 98.0 (97.9) |
| Resolution (Å) | 35.47-2.12 (2.18-2.12) | 41.34-2.35 (2.43-2.35) | 41.34-2.61 (2.73-2.61) |
| $R_{meas}$ | 0.163 (2.711) | 0.249 (1.789) | 0.354 (3.076) |
| $R_{pim}$ | 0.033 (0.587) | 0.107 (0.803) | 0.144 (1.286) |
| $\langle I/\sigma(I) \rangle$ | 14.0 (2.0) | 4.8 (1.7) | 4.9 (1.7) |
| Redundancy or Multiplicity | 24.6 (20.9) | 5.0 (4.6) | 5.4 (5.5) |
| $CC_{1/2}$ | 0.999 (0.526) | 0.982 (0.311) | 0.953 (0.265) |
| CC* | 1.0 (0.83) | 0.995 (0.689) | 0.988 (0.647) |
| $\chi^2$ | 0.95 (0.98) | 1.00 (1.00) | 1.08 (0.93) |
| Proteins/ASU | 1 | 1 | 1 |
| Percent Solvent (%) | 68 | 68.1 | 68.1 |
| Matthews Coefficient | 3.85 | 3.85 | 3.85 |
| Wilson B-Factor ($Å^2$) | 44.6 | 39.55 | 49.21 |
| No. Observations | 270013 (17875) | 38986 (3538) | 31400 (3740) |
| No. Reflections | 10969 (857) | 7868 (763) | 5804 (684) |
| **Refinement** | | | |
| Resolution | 35.47 (2.12) | 41.34 (2.350) | 41.34 (2.61) |
| $R_{Work}$ / $R_{Free}$ | 0.2199 / 0.2577 | 0.2000 / 0.2489 | 0.2122 / 0.2662 |
| R.M.S. Deviations | | | |
| Bond-Lengths | 0.0159 | 0.0067 | 0.0088 |
| Bond-Angles | 1.6861 | 1.2615 | 1.3173 |
| **Numbers** | | | |
| Residues | 106 | 106 | 106 |
| Atoms / Non-Hydrogens | 1760 / 918 | 1767 / 928 | 1734 / 911 |
| Protein: Overall / Heavy / Backbone | 1652 / 834 / 428 | 1633 / 826 / 424 | 1633 / 826 / 424 |
| Ions and Ligands: Overall / Heavy | 56 / 32 | 72 / 40 | 41 / 25 |
| Waters | 52 | 62 | 60 |
| B-factors | | | |
| Protein: Overall / Heavy / Backbone | 73.18 / 66.47 / 62.1 | 67.14 / 60.64 / 55.05 | 70.52 / 64.1 / 59.65 |
| Ions and Ligands: Overall / Heavy | 50.88 / 53.91 | 41.86 / 46.7 | 54.63 / 56.42 |
| Waters | 62.01 | 60.39 | 55.25 |
| **MolProbity** | | | |
| Clash Score | 0.00 | 0.00 | 0.00 |
| MolProbity Score | 0.52 | 0.50 | 0.50 |
| Ramachandran | | | |
| Allowed | 100 % | 100 % | 100 % |
| Favored | 99.04 % | 99.04 % | 99.04 % |
| Outliers | 0 % | 0 % | 0 % |
| Rotamers | | | |
| Allowed | 98.92 % | 100 % | 100 % |
| Favored | 98.92 % | 100 % | 98.91 % |
| Outliers | 1.08 % | 0 % | 0 % |
| Clashes | 0 | 0 | 0 |

### Table S4:

| | Dimer BMP-10 Dimer, Chimera |
|---|---|
| **PDB** | **9DPY** |
| **Data Collection** | |
| X-Ray Source | APS 24-ID-E |
| Wavelength (Å) | 0.97918 |
| Detector | DECTRIS EIGER X 16M |
| Linear Cell Variation | N/A |
| Number of Crystals | 1 |
| **Data Reduction** | |
| Spacegroup | C 2 2 $2_1$ (20) |
| a, b, c (Å) | 42.75, 123.04, 120.56 |
| α. β, γ (°) | 90.00, 90.00, 90.00 |
| Completeness (%) | 98.8 (97.2) |
| Resolution (Å) | 61.52-1.77 (1.87-1.77) |
| $R_{meas}$ | 0.086 (2.713) |
| $R_{pim}$ | 0.029 (0.889) |
| <I/σ(I)> | 12.2 (1.0) |
| Redundancy or Multiplicity | 8.9 (8.9) |
| $CC_{1/2}$ | 0.998 (0.531) |
| CC* | 0.999 (0.833) |
| $\chi^2$ | 1.05 (0.84) |
| Proteins/ASU | 2 |
| Percent Solvent (%) | 63.4 |
| Matthews Coefficient | 3.36 |
| Wilson B-Factor ($Å^2$) | 35.76 |
| No. Observations | 276852 (39065) |
| No. Reflections | 31025 (4391) |
| **Refinement** | |
| Resolution (Å) | 61.52 (1.77) |
| $R_{Work}$ / $R_{Free}$ | 0.2141 / 0.2512 |
| R.M.S. Deviations | |
| Bond-Lengths (Å) | 0.0102 |
| Bond-Angles (°) | 1.4773 |
| **Numbers** | |
| Residues | 210 |
| Atoms / Non-Hydrogens | 3538 / 1875 |
| Protein: Overall / Heavy / Backbone | 3327 / 1664 / 840 |
| Ions and Ligands: Overall / Heavy | 67 / 67 |
| Waters | 144 |
| B-factors | |
| Protein: Overall / Heavy / Backbone | 67.14 / 60.14 / 55.39 |
| Ions and Ligands: Overall / Heavy | 61.93 / 61.93 |
| Waters | 61.2 |
| **MolProbity** | |
| Clash Score | 0 |
| MolProbity Score | 0.53 |
| Ramachandran | |
| Allowed | 99.52 % |
| Favored | 98.06 % |
| Outliers | 0.49 % |
| Rotamers | |
| Allowed | 98.9 % |
| Favored | 97.8 % |
| Outliers | 1.1 % |
| Clashes | 0 |

**Table S5:**

| PDB | BMP-9 G389S Dimer, First 32nd 9DPS | BMP-9 G389S Dimer, Last 32nd 9DPT | BMP-9 G389S Dimer, Acid 9DPU |
|---|---|---|---|
| **Data Collection** | | | |
| X-Ray Source | APS 22-ID | APS 22-ID | APS 22-ID |
| Wavelength (Å) | 1 | 1 | 1 |
| Detector | DECTRIS EIGER X 16M | DECTRIS EIGER X 16M | DECTRIS EIGER X 16M |
| Linear Cell Variation | 1.18 % | 1.18 % | N/A |
| Number of Crystals | 8 | 8 | 1 |
| **Data Reduction** | | | |
| Spacegroup | I $4_1$ 2 2 (98) | I $4_1$ 2 2 (98) | I $4_1$ 2 2 (98) |
| a, b, c (Å) | 70.77, 70.77, 145.52 | 70.77, 70.77, 145.52 | 71.59, 71.59, 146.72 |
| α. β, γ (°) | 90.00, 90.00, 90.00 | 90.00, 90.00, 90.00 | 90.00, 90.00, 90.00 |
| Completeness (%) | 99.5 (99.9) | 99.7 (98.9) | 100.0 (100.0) |
| Resolution (Å) | 63.65-2.06 (2.12-2.06) | 50.04-2.49 (2.59-2.49) | 41.66-2.10 (2.16-2.10) |
| $R_{meas}$ | 0.129 (1.142) | 0.464 (3.782) | 0.159 (1.489) |
| $R_{pim}$ | 0.052 (0.458) | 0.192 (1.720) | 0.032 (0.287) |
| $\langle I/\sigma(I)\rangle$ | 7.2 (2.1) | 5.1 (1.6) | 13.9 (4.4) |
| Redundancy or Multiplicity | 5.8 (6.0) | 5.9 (5.1) | 25.6 (26.9) |
| $CC_{1/2}$ | 0.993 (0.794) | 0.828 (0.252) | 0.997 (0.970) |
| CC* | 0.998 (0.941) | 0.952 (0.634) | 0.999 (0.992) |
| $\chi^2$ | 1.09 (1.21) | 0.91 (0.74) | 0.89 (0.85) |
| Proteins/ASU | 1 | 1 | 1 |
| Percent Solvent (%) | 68.1 | 68.1 | 69.1 |
| Matthews Coefficient | 3.86 | 3.86 | 3.98 |
| Wilson B-Factor (Å$^2$) | 38.13 | 25.38 | 39.19 |
| No. Observations | 68104 (5382) | 39927 (3722) | 295471 (25019) |
| No. Reflections | 11744 (903) | 6799 (733) | 11543 (929) |
| **Refinement** | | | |
| Resolution | 31.82 (2.06) | 50.04 (2.49) | 41.66 (2.10) |
| $R_{Work}$ / $R_{Free}$ | 0.2163 / 0.2490 | 0.2392 / 0.2791 | 0.1916 / 0.2236 |
| R.M.S. Deviations | | | |
| Bond-Lengths | 0.0094 | 0.0084 | 0.0091 |
| Bond-Angles | 1.4591 | 1.4242 | 1.4151 |
| **Numbers** | | | |
| Residues | 105 | 105 | 105 |
| Atoms / Non-Hydrogens | 1797 / 959 | 1771 / 941 | 1790 / 957 |
| Protein: Overall / Heavy / Backbone | 1648 / 834 / 424 | 1630 / 824 / 420 | 1636 / 827 / 421 |
| Ions and Ligands: Overall / Heavy | 52 / 28 | 51 / 27 | 48 / 24 |
| Waters | 97 | 90 | 106 |
| B-factors | | | |
| Protein: Overall / Heavy / Backbone | 63.38 / 57.65 / 55.09 | 37.84 / 34.73 / 31.79 | 72.42 / 65.52 / 58.66 |
| Ions and Ligands: Overall / Heavy | 57.66 / 54.46 | 40.86 / 37.52 | 73.54 / 65.1 |
| Waters | 58.9 | 37.87 | 66.31 |
| **MolProbity** | | | |
| Clash Score | 0.00 | 0.00 | 0.00 |
| MolProbity Score | 0.75 | 0.52 | 0.52 |
| Ramachandran | | | |
| Allowed | 99.03 % | 100 % | 100 % |
| Favored | 99.03 % | 99.03 % | 99.03 % |
| Outliers | 0.97 % | 0 % | 0 % |
| Rotamers | | | |
| Allowed | 97.87 % | 98.92 % | 98.94 % |
| Favored | 96.81 % | 97.85 % | 97.87 % |
| Outliers | 2.13 % | 1.08 % | 1.06 % |
| Clashes | 0 | 0 | 0 |

**Table S6:**

| PDB | BMP-9 G389S K357R Dimer, First 32nd 9DPV | Dimer BMP-9 G389S K357R Dimer, Last 32nd 9DPW | BMP-9 G389S K357R Dimer, Acid 9DPX |
|---|---|---|---|
| **Data Collection** | | | |
| X-Ray Source | APS 22-ID | APS 22-ID | APS 22-ID |
| Wavelength (Å) | 1 | 1 | 1 |
| Detector | DECTRIS EIGER X 16M | DECTRIS EIGER X 16M | DECTRIS EIGER X 16M |
| Linear Cell Variation | 0.73 % | 0.73 % | N/A |
| Number of Crystals | 10 | 10 | 1 |
| **Data Reduction** | | | |
| Spacegroup | I $4_1$ 2 2 (98) | I 41 2 2 (98) | I $4_1$ 2 2 (98) |
| a, b, c (Å) | 71.11, 71.11, 145.55 | 71.10, 71.10, 145.55 | 71.29, 71.29, 146.38 |
| α. β, γ (°) | 90.00, 90.00, 90.00 | 90.00, 90.00, 90.00 | 90.00, 90.00, 90.00 |
| Completeness (%) | 98.8 (86.7) | 99.6 (99.6) | 100.0 (100.0) |
| Resolution (Å) | 41.37-1.99 (2.04-1.99) | 40.08-2.71 (2.84-2.71) | 64.09-2.10 (2.16-2.10) |
| $R_{meas}$ | 0.167 (1.693) | 0.494 (4.951) | 0.274 (3.484) |
| $R_{pim}$ | 0.062 (0.659) | 0.181 (1.779) | 0.055 (0.678) |
| $<I/\sigma(I)>$ | 6.4 (1.2) | 4.2 (1.4) | 8.5 (2.2) |
| Redundancy or Multiplicity | 7.0 (5.6) | 6.9 (7.4) | 24.9 (26.2) |
| $CC_{1/2}$ | 0.988 (0.645) | 0.948 (0.309) | 0.938 (0.909) |
| CC* | 0.997 (0.886) | 0.987 (0.687) | 0.984 (0.976) |
| $\chi^2$ | 1.00 (0.95) | 0.97 (1.04) | 1.01 (1.05) |
| Proteins/ASU | 1 | 1 | 1 |
| Percent Solvent (%) | 68.4 | 68.4 | 68.5 |
| Matthews Coefficient | 3.89 | 3.89 | 3.9 |
| Wilson B-Factor (Å$^2$) | 36.55 | 51.45 | 41.17 |
| No. Observations | 91713 (4637) | 37180 (5191) | 284389 (23875) |
| No. Reflections | 13055 (832) | 5360 (706) | 11416 (913) |
| **Refinement** | | | |
| Resolution | 35.55 (1.99) | 40.08 (2.71) | 64.09 (2.10) |
| $R_{Work}$ / $R_{Free}$ | 0.2238 / 0.2649 | 0.2235 / 0.2986 | 0.2275 / 0.2675 |
| R.M.S. Deviations | | | |
| Bond-Lengths | 0.0086 | 0.0094 | 0.0167 |
| Bond-Angles | 1.3556 | 1.4325 | 1.7516 |
| **Numbers** | | | |
| Residues | 105 | 105 | 106 |
| Atoms / Non-Hydrogens | 1756 / 934 | 1733 / 902 | 1765 / 929 |
| Protein: Overall / Heavy / Backbone | 1632 / 826 / 420 | 1633 / 826 / 420 | 1645 / 833 / 425 |
| Ions and Ligands: Overall / Heavy | 37 / 21 | 54 / 30 | 51 / 27 |
| Waters | 87 | 46 | 69 |
| B-factors | | | |
| Protein: Overall / Heavy / Backbone | 60.4 / 55.46 / 52.14 | 73.03 / 66.84 / 61.74 | 69.18 / 63.02 / 59.53 |
| Ions and Ligands: Overall / Heavy | 33.0 / 37.94 | 63.16 / 62.22 | 50.91 / 51.17 |
| Waters | 34.49 | 60.11 | 44.06 |
| **MolProbity** | | | |
| Clash Score | 0.00 | 0.00 | 0.00 |
| MolProbity Score | 0.52 | 0.52 | 0.75 |
| Ramachandran | | | |
| Allowed | 100 % | 100 % | 99.04 % |
| Favored | 99.03 % | 99.03 % | 99.04 % |
| Outliers | 0 % | 0 % | 0.96 % |
| Rotamers | | | |
| Allowed | 98.92 % | 98.92 % | 97.87 % |
| Favored | 97.85 % | 98.92 % | 97.87 % |
| Outliers | 1.08 % | 1.08 % | 2.13 % |
| Clashes | 0 | 0 | 0 |

**Table S7:** Oligos

| | |
|---|---|
| F9-G389S | AGGTGTCCAAGGCCTGCTGTGTGCCCACC |
| R9-G389S | GCCTTGGACACCTTTGTGGGGAACTTGAGATGC |
| F10-S385G | AAGCTGGCAAAGCCTGCTGTGTGCCCAC |
| R10-S385G | GCTTTGCCAGCTTTCTGGGAATTCTTGAGGTGG |
| F9-K357R | AGTGTCGTGGCGGCTGCTTCTTCCCCTTGGCT |
| R9-K357R | CCGCCACGACACTCGTAGGCTTCATACTCCTTGGG |
| F10-R353K | AATGCAAGGGTGTTTGTAACTACCCCCTGG |
| R10-R353K | ACACCCTTGCATTCATAGGCTTCGTATCCAGG |

**Table S8:** Intra-chain Disulfide Dihedrals

| | Cys-327 | Cys-356 | Cys-360 | Cys-393 | Cys-426 | Cys-428 |
|---|---|---|---|---|---|---|
| $\chi_1$ (±60°±15°, ±180°±15°) | -172° | -64° | -61° | 176° | -63° | -72° |
| $\chi_2$ (±90°±30°) | 71° | -66° | -85° | 67° | -74° | -78° |
| $\chi_3$ (±90°±15°) | 79° | -86° | -89° | 79° | -86° | -89° |

**Table S9:** Hydrodynamic measurements and partial concentrations of each oligomeric species and concentration and according to the genetic algorithm - Monte Carlo analysis. Values in parentheses are 95 % confidence intervals.

| **Sample:** | **Monomer %:** | **Dimer %:** | **Tetramer %:** |
|---|---|---|---|
| Green: 1.7 µM (215 nm) | 26.5 | 62.4 | 11.1 |
| Red: 6.2 µM (230nm) | 25.8 | 68.8 | 5.4 |
| Blue: 17.6 µM (280nm) | 37.0 | 51.3 | 11.7 |
| Magenta: 60.5 µM (295nm) | 42.9 | 57.1 | Not detected |
| Frictional Ratio, f/f0 | 1.33 (1.0 – 1.38) | 1.46 (1.21 – 1.71) | 1.74 (1.52 – 1.96) |
| Sedimentation Coefficient (s) | 1.25 (1.11 – 1.39) | 2.12 (1.92 – 2.33) | 2.75 (2.32 – 3.19) |
| Experimental molar mass (kDa) | 10.3 (7.2 – 13.4) | 26.0 (22.2 – 29.7) | 50.0 (44.7 – 55.2) |
| Theoretical molar mass (kDa) | 12.1 | 24.2 | 48.4 |

### AlbSigPep-His-(X)-Pro-(X)-BMP9 Construct

```
atg aag tgg gta acc ttt ctc ctc ctc ctc ttc atc tcc ggt tct gcc ttt tct gcg gcc gca aac gct cat cac
 M   K   W   V   T   F   L   L   L   L   F   I   S   G   S   A   F   S   A   A   A   N   A   H   H
cat cac cac cat cat cat att gaa gga cgt aag cca ctg cag agc tgg gaa cag ggg tct gct ggg gga aac gcc
 H   H   H   H   H   H   I   E   G   R   K   P   L   Q   S   W   E   Q   G   S   A   G   G   N   A
cac agc cca ctg ggg gtg cct gga ggt ggg ctg cct gag cac acc ttc aac ctg aag atg ttt ctg gag aac gtg
 H   S   P   L   G   V   P   G   G   G   L   P   E   H   T   F   N   L   K   M   F   L   E   N   V
aag gtg gat ttc ctg cgc agc ctt aac ctg agt ggg gtc cct tcg cag gac aaa acc agg gtg gag ccg ccg cag
 K   V   D   F   L   R   S   L   N   L   S   G   V   P   S   Q   D   K   T   R   V   E   P   P   Q
tac atg att gac ctg tac aac agg tac acg tcc gat aag tcg act acg cca gcg tcc aac att gtg cgg agc ttc
 Y   M   I   D   L   Y   N   R   Y   T   S   D   K   S   T   T   P   A   S   N   I   V   R   S   F
agc atg gaa gat gcc atc tcc ata act gcc aca gag gac ttc ccc ttc cag aag cac atc ttg ctc ttc aac atc
 S   M   E   D   A   I   S   I   T   A   T   E   D   F   P   F   Q   K   H   I   L   L   F   N   I
tcc att cct agg cat gag cag atc acc aga gct gag ctc cga ctc tat gtc tcc tgt caa aat cac gtg gac ccc
 S   I   P   R   H   E   Q   I   T   R   A   E   L   R   L   Y   V   S   C   Q   N   H   V   D   P
tct cat gac ctg aaa gga agc gtg gtc att tat gat gtt ctg gat gga aca gat gcc tgg gat agt gct aca gag
 S   H   D   L   K   G   S   V   V   I   Y   D   V   L   D   G   T   D   A   W   D   S   A   T   E
acc aag acc ttc ctg gtg tcc cag gac att cag gat gag ggc tgg gag acc ttg gaa gtg tcc agc gcc gtg aag
 T   K   T   F   L   V   S   Q   D   I   Q   D   E   G   W   E   T   L   E   V   S   S   A   V   K
cgc tgg gtc cgg tcc gac tcc acc aag agc aaa aat aag ctg gaa gtg act gtg gag agc cac agg aag ggc tgc
 R   W   V   R   S   D   S   T   K   S   K   N   K   L   E   V   T   V   E   S   H   R   K   G   C
gac acg ctg gac atc agt gtc ccc cca ggt tcc aga aac ctg ccc ttc ttt gtt gtc ttc tcc aat gac cac agc
 D   T   L   D   I   S   V   P   P   G   S   R   N   L   P   F   F   V   V   F   S   N   D   H   S
agt ggg acc aag gag acc agg ctg gag ctg agg gag atg atc agc cat gaa caa gag agc gtg ctc aag aag ctg
 S   G   T   K   E   T   R   L   E   L   R   E   M   I   S   H   E   Q   E   S   V   L   K   K   L
tcc aag gac ggc tcc aca gag gca ggt gag agc agt cac gag gag gac acg gat ggc cac gtg gct gcg ggg tcg
 S   K   D   G   S   T   E   A   G   E   S   S   H   E   E   D   T   D   G   H   V   A   A   G   S
act tta gcc atc gag ggg cgg agc tcc ggg gct ggc agc cac tgt caa aag acc tcc ctg cgg gta aac ttc gag
 T   L   A   I   E   G   R   S   S   G   A   G   S   H   C   Q   K   T   S   L   R   V   N   F   E
gac atc ggc tgg gac agc tgg atc att gca ccc aag gag tat gaa gcc tac gag tgt aag ggc ggc tgc ttc ttc
 D   I   G   W   D   S   W   I   I   A   P   K   E   Y   E   A   Y   E   C   K   G   G   C   F   F
ccc ttg gct gac gat gtg acg ccg acg aaa cac gct atc gtg cag acc ctg gtg cat ctc aag ttc ccc aca aag
 P   L   A   D   D   V   T   P   T   K   H   A   I   V   Q   T   L   V   H   L   K   F   P   T   K
gtg ggc aag gcc tgc tgt gtg ccc acc aaa ctg agc ccc atc tcc gtc ctc tac aag gat gac atg ggg gtg ccc
 V   G   K   A   C   C   V   P   T   K   L   S   P   I   S   V   L   Y   K   D   D   M   G   V   P
acc ctc aag tac cat tac gag ggc atg agc gtg gca gag tgt ggg tgc agg tag ctc gag
 T   L   K   Y   H   Y   E   G   M   S   V   A   E   C   G   C   R   -
```

### AlbSigPep-His-(X)-Pro-(fr)-BMP9 Construct

```
atg aag tgg gta acc ttt ctc ctc ctc ctc ttc atc tcc ggt tct gcc ttt tct gcg gcc gca cat cac cat cac
 M   K   W   V   T   F   L   L   L   L   F   I   S   G   S   A   F   S   A   A   A   H   H   H   H
cac cat cat cat att gaa gga cgt aag cca ctg cag agc tgg gga cga ggg tct gct ggg gga aac gcc cac agc
 H   H   H   H   I   E   G   R   K   P   L   Q   S   W   G   R   G   S   A   G   G   N   A   H   S
cca ctg ggg gtg cct gga ggt ggg ctg cct gag cac acc ttc aac ctg aag atg ttt ctg gag aac gtg aag gtg
 P   L   G   V   P   G   G   G   L   P   E   H   T   F   N   L   K   M   F   L   E   N   V   K   V
gat ttc ctg cgc agc ctt aac ctg agt ggg gtc cct tcg cag gac aaa acc agg gtg gag ccg ccg cag tac atg
 D   F   L   R   S   L   N   L   S   G   V   P   S   Q   D   K   T   R   V   E   P   P   Q   Y   M
att gac ctg tac aac agg tac acg tcc gat aag tcg act acg cca gcg tcc aac att gtg cgg agc ttc agc atg
 I   D   L   Y   N   R   Y   T   S   D   K   S   T   T   P   A   S   N   I   V   R   S   F   S   M
gaa gat gcc atc tcc ata act gcc aca gag gac ttc ccc ttc cag aag cac atc ttg ctc ttc aac atc tcc att
 E   D   A   I   S   I   T   A   T   E   D   F   P   F   Q   K   H   I   L   L   F   N   I   S   I
cct agg cat gag cag atc acc aga gct gag ctc cga ctc tat gtc tcc tgt caa aat cac gtg gac ccc tct cat
 P   R   H   E   Q   I   T   R   A   E   L   R   L   Y   V   S   C   Q   N   H   V   D   P   S   H
gac ctg aaa gga agc gtg gtc att tat gat gtt ctg gat gga aca gat gcc tgg gat agt gct aca gag acc aag
 D   L   K   G   S   V   V   I   Y   D   V   L   D   G   T   D   A   W   D   S   A   T   E   T   K
acc ttc ctg gtg tcc cag gac att cag gat gag ggc tgg gag acc ttg gaa gtg tcc agc gcc gtg aag cgc tgg
 T   F   L   V   S   Q   D   I   Q   D   E   G   W   E   T   L   E   V   S   S   A   V   K   R   W
gtc cgg tcc gac tcc acc aag agc aaa aat aag ctg gaa gtg act gtg gag agc cac agg aag ggc tgc gac acg
 V   R   S   D   S   T   K   S   K   N   K   L   E   V   T   V   E   S   H   R   K   G   C   D   T
ctg gac atc agt gtc ccc cca ggt tcc aga aac ctg ccc ttc ttt gtt gtc ttc tcc aat gac cac agc agt ggg
 L   D   I   S   V   P   P   G   S   R   N   L   P   F   F   V   V   F   S   N   D   H   S   S   G
acc aag gag acc agg ctg gag ctg agg gag atg atc agc cat gaa caa gag agc gtg ctc aag aag ctg tcc aag
 T   K   E   T   R   L   E   L   R   E   M   I   S   H   E   Q   E   S   V   L   K   K   L   S   K
gac ggc tcc aca gag gca ggt gag agc agt cac gag gag gac acg gat ggc cac gtg gct gcg ggg tcg act tta
 D   G   S   T   E   A   G   E   S   S   H   E   E   D   T   D   G   H   V   A   A   G   S   T   L
gcc agg cgg aaa agg agc gcc ggg gct ggc agc cac tgt caa aag acc tcc ctg cgg gta aac ttc gag gac atc
 A   R   R   K   R   S   A   G   A   G   S   H   C   Q   K   T   S   L   R   V   N   F   E   D   I
ggc tgg gac agc tgg atc att gca ccc aag gag tat gaa gcc tac gag tgt aag ggc ggc tgc ttc ttc ccc ttg
 G   W   D   S   W   I   I   A   P   K   E   Y   E   A   Y   E   C   K   G   G   C   F   F   P   L
gct gac gat gtg acg ccg acg aaa cac gct atc gtg cag acc ctg gtg cat ctc aag ttc ccc aca aag gtg ggc
 A   D   D   V   T   P   T   K   H   A   I   V   Q   T   L   V   H   L   K   F   P   T   K   V   G
aag gcc tgc tgt gtg ccc acc aaa ctg agc ccc atc tcc gtc ctc tac aag gat gac atg ggg gtg ccc acc ctc
 K   A   C   C   V   P   T   K   L   S   P   I   S   V   L   Y   K   D   D   M   G   V   P   T   L
aag tac cat tac gag ggc atg agc gtg gca gag tgt ggg tgc agg tag taa tct aga
 K   Y   H   Y   E   G   M   S   V   A   E   C   G   C   R   -   -
```

### AlbSigPep-His-(X)-Pro-(X)-BMP10 Construct

```
atg aag tgg gta acc ttt ctc ctc ctc ctc ttc atc tcc ggt tct gcc ttt tct gcg gcc gca cat cac cat cac
 M   K   W   V   T   F   L   L   L   L   F   I   S   G   S   A   F   S   A   A   A   H   H   H   H
cac cat cat cat att gaa gga cgt agt agc ccc atc atg aac cta gag cag tct cct ctg gaa gaa gat atg tcc
 H   H   H   H   I   E   G   R   S   S   P   I   M   N   L   E   Q   S   P   L   E   E   D   M   S
ctc ttt ggt gat gtt ttc tca gag caa gac ggt gtc gac ttt aac aca ctg ctc cag agc atg aag gat gag ttt
 L   F   G   D   V   F   S   E   Q   D   G   V   D   F   N   T   L   L   Q   S   M   K   D   E   F
ctt aag aca cta aac ctc tct gac atc ccc acg cag gat tca gcc aag gtg gac cca cca gag tac atg ttg gaa
 L   K   T   L   N   L   S   D   I   P   T   Q   D   S   A   K   V   D   P   P   E   Y   M   L   E
ctc tac aac aaa ttt gca aca gat cgg acc tcc atg ccc tct gcc aac atc att agg agt ttc aag aat gaa gat
 L   Y   N   K   F   A   T   D   R   T   S   M   P   S   A   N   I   I   R   S   F   K   N   E   D
ctg ttt tcc cag ccg gtc agt ttt aat ggg ctc cga aaa tac ccc ctc ctc ttc aat gtg tcc att cct cac cat
 L   F   S   Q   P   V   S   F   N   G   L   R   K   Y   P   L   L   F   N   V   S   I   P   H   H
gaa gag gtc atc atg gct gaa ctt agg cta tac aca ctg gtg caa agg gat cgt atg ata tac gat gga gta gac
 E   E   V   I   M   A   E   L   R   L   Y   T   L   V   Q   R   D   R   M   I   Y   D   G   V   D
cgg aaa att acc att ttt gaa gtg ctg gag agc aaa ggg gat aat gag gga gaa aga aac atg ctg gtc ttg gtg
 R   K   I   T   I   F   E   V   L   E   S   K   G   D   N   E   G   E   R   N   M   L   V   L   V
tct ggg gag ata tat gga acc aac agt gag tgg gag act ttt gat gtc aca gat gcc atc aga cgt tgg caa aag
 S   G   E   I   Y   G   T   N   S   E   W   E   T   F   D   V   T   D   A   I   R   R   W   Q   K
tca ggc tca tcc acc cac cag ctg gag gtc cac att gag agc aaa cac gat gaa gct gag gat gcc agc agt gat
 S   G   S   S   T   H   Q   L   E   V   H   I   E   S   K   H   D   E   A   E   D   A   S   S   D
acc cta gaa ata gat acc agt gcc cag aat aag cat aac cct ttg ctc atc gtg ttt tct gat gac caa agc agt
 T   L   E   I   D   T   S   A   Q   N   K   H   N   P   L   L   I   V   F   S   D   D   Q   S   S
gac aag gag agg aag gag gaa ctg aat gaa atg att tcc cat gag caa ctt cca gag ctg gac aac ttg ggc ctg
 D   K   E   R   K   E   E   L   N   E   M   I   S   H   E   Q   L   P   E   L   D   N   L   G   L
gat agc ttt tcc agt gga cct ggg gaa gag gct ttg ttg cag atg aga tca aac atc atc tat gac tcc act ggc
 D   S   F   S   S   G   P   G   E   E   A   L   L   Q   M   R   S   N   I   I   Y   D   S   T   G
att gaa gga cgt aac gcc aaa gga aac tac tgt aag agg acc ccg ctc tac atc gac ttc aag gag att ggg tgg
 I   E   G   R   N   A   K   G   N   Y   C   K   R   T   P   L   Y   I   D   F   K   E   I   G   W
gac tcc tgg atc atc gct ccg cct gga tac gaa gcc tat gaa tgc cgt ggt gtt tgt aac tac ccc ctg gca gag
 D   S   W   I   I   A   P   P   G   Y   E   A   Y   E   C   R   G   V   C   N   Y   P   L   A   E
cat ctc aca ccc aca aag cat gca att atc cag gcc ttg gtc cac ctc aag aat tcc cag aaa gct tcc aaa gcc
 H   L   T   P   T   K   H   A   I   I   Q   A   L   V   H   L   K   N   S   Q   K   A   S   K   A
tgc tgt gtg ccc aca aag cta gag ccc atc tcc atc ctc tat tta gac aaa ggc gtc gtc acc tac aag ttt aaa
 C   C   V   P   T   K   L   E   P   I   S   I   L   Y   L   D   K   G   V   V   T   Y   K   F   K
tac gaa ggc atg gcc gtc tcc gaa tgt ggc tgt aga tag
 Y   E   G   M   A   V   S   E   C   G   C   R   -
```

### AlbSigPep-His-(X)-Pro-(fr)-BMP10 Construct

```
atg aag tgg gta acc ttt ctc ctc ctc ctc ttc atc tcc ggt tct gcc ttt tct gcg gcc gca cat cac cat cac
 M   K   W   V   T   F   L   L   L   L   F   I   S   G   S   A   F   S   A   A   A   H   H   H   H
cac cat cat cat att gaa gga cgt agt agc ccc atc atg aac cta gag cag tct cct ctg gaa gaa gat atg tcc
 H   H   H   H   I   E   G   R   S   S   P   I   M   N   L   E   Q   S   P   L   E   E   D   M   S
ctc ttt ggt gat gtt ttc tca gag caa gac ggt gtc gac ttt aac aca ctg ctc cag agc atg aag gat gag ttt
 L   F   G   D   V   F   S   E   Q   D   G   V   D   F   N   T   L   L   Q   S   M   K   D   E   F
ctt aag aca cta aac ctc tct gac atc ccc acg cag gat tca gcc aag gtg gac cca cca gag tac atg ttg gaa
 L   K   T   L   N   L   S   D   I   P   T   Q   D   S   A   K   V   D   P   P   E   Y   M   L   E
ctc tac aac aaa ttt gca aca gat cgg acc tcc atg ccc tct gcc aac atc att agg agt ttc aag aat gaa gat
 L   Y   N   K   F   A   T   D   R   T   S   M   P   S   A   N   I   I   R   S   F   K   N   E   D
ctg ttt tcc cag ccg gtc agt ttt aat ggg ctc cga aaa tac ccc ctc ctc ttc aat gtg tcc att cct cac cat
 L   F   S   Q   P   V   S   F   N   G   L   R   K   Y   P   L   L   F   N   V   S   I   P   H   H
gaa gag gtc atc atg gct gaa ctt agg cta tac aca ctg gtg caa agg gat cgt atg ata tac gat gga gta gac
 E   E   V   I   M   A   E   L   R   L   Y   T   L   V   Q   R   D   R   M   I   Y   D   G   V   D
cgg aaa att acc att ttt gaa gtg ctg gag agc aaa ggg gat aat gag gga gaa aga aac atg ctg gtc ttg gtg
 R   K   I   T   I   F   E   V   L   E   S   K   G   D   N   E   G   E   R   N   M   L   V   L   V
tct ggg gag ata tat gga acc aac agt gag tgg gag act ttt gat gtc aca gat gcc atc aga cgt tgg caa aag
 S   G   E   I   Y   G   T   N   S   E   W   E   T   F   D   V   T   D   A   I   R   R   W   Q   K
tca ggc tca tcc acc cac cag ctg gag gtc cac att gag agc aaa cac gat gaa gct gag gat gcc agc agt gat
 S   G   S   S   T   H   Q   L   E   V   H   I   E   S   K   H   D   E   A   E   D   A   S   S   D
acc cta gaa ata gat acc agt gcc cag aat aag cat aac cct ttg ctc atc gtg ttt tct gat gac caa agc agt
 T   L   E   I   D   T   S   A   Q   N   K   H   N   P   L   L   I   V   F   S   D   D   Q   S   S
gac aag gag agg aag gag gaa ctg aat gaa atg att tcc cat gag caa ctt cca gag ctg gac aac ttg ggc ctg
 D   K   E   R   K   E   E   L   N   E   M   I   S   H   E   Q   L   P   E   L   D   N   L   G   L
gat agc ttt tcc agt gga cct ggg gaa gag gct ttg ttg cag atg aga tca aac atc atc tat gac tcc act gcc
 D   S   F   S   S   G   P   G   E   E   A   L   L   Q   M   R   S   N   I   I   Y   D   S   T   A
cga atc aga agg aac gcc aaa gga aac tac tgt aag agg acc ccg ctc tac atc gac ttc aag gag att ggg tgg
 R   I   R   R   N   A   K   G   N   Y   C   K   R   T   P   L   Y   I   D   F   K   E   I   G   W
gac tcc tgg atc atc gct ccg cct gga tac gaa gcc tat gaa tgc cgt ggt gtt tgt aac tac ccc ctg gca gag
 D   S   W   I   I   A   P   P   G   Y   E   A   Y   E   C   R   G   V   C   N   Y   P   L   A   E
cat ctc aca ccc aca aag cat gca att atc cag gcc ttg gtc cac ctc aag aat tcc cag aaa gct tcc aaa gcc
 H   L   T   P   T   K   H   A   I   I   Q   A   L   V   H   L   K   N   S   Q   K   A   S   K   A
tgc tgt gtg ccc aca aag cta gag ccc atc tcc atc ctc tat tta gac aaa ggc gtc gtc acc tac aag ttt aaa
 C   C   V   P   T   K   L   E   P   I   S   I   L   Y   L   D   K   G   V   V   T   Y   K   F   K
tac gaa ggc atg gcc gtc tcc gaa tgt ggc tgt aga tag
 Y   E   G   M   A   V   S   E   C   G   C   R   -
```

### AlbSigPep-His8-(X)-Pro9-(X)-BMP10 Crystal Chimera

```
atg aag tgg gta acc ttt ctc ctc ctc ctc ttc atc tcc ggt tct gcc ttt tct gcg gcc gca aac gct cat cac
 M   K   W   V   T   F   L   L   L   L   F   I   S   G   S   A   F   S   A   A   A   N   A   H   H
cat cac cac cat cat cat att gaa gga cgt aag cca ctg cag agc tgg gaa cag ggg tct gct ggg gga aac gcc
 H   H   H   H   H   H   I   E   G   R   K   P   L   Q   S   W   E   Q   G   S   A   G   G   N   A
cac agc cca ctg ggg gtg cct gga ggt ggg ctg cct gag cac acc ttc aac ctg aag atg ttt ctg gag aac gtg
 H   S   P   L   G   V   P   G   G   G   L   P   E   H   T   F   N   L   K   M   F   L   E   N   V
aag gtg gat ttc ctg cgc agc ctt aac ctg agt ggg gtc cct tcg cag gac aaa acc agg gtg gag ccg ccg cag
 K   V   D   F   L   R   S   L   N   L   S   G   V   P   S   Q   D   K   T   R   V   E   P   P   Q
tac atg att gac ctg tac aac agg tac acg tcc gat aag tcg act acg cca gcg tcc aac att gtg cgg agc ttc
 Y   M   I   D   L   Y   N   R   Y   T   S   D   K   S   T   T   P   A   S   N   I   V   R   S   F
agc atg gaa gat gcc atc tcc ata act gcc aca gag gac ttc ccc ttc cag aag cac atc ttg ctc ttc aac atc
 S   M   E   D   A   I   S   I   T   A   T   E   D   F   P   F   Q   K   H   I   L   L   F   N   I
tcc att cct agg cat gag cag atc acc aga gct gag ctc cga ctc tat gtc tcc tgt caa aat cac gtg gac ccc
 S   I   P   R   H   E   Q   I   T   R   A   E   L   R   L   Y   V   S   C   Q   N   H   V   D   P
tct cat gac ctg aaa gga agc gtg gtc att tat gat gtt ctg gat gga aca gat gcc tgg gat agt gct aca gag
 S   H   D   L   K   G   S   V   V   I   Y   D   V   L   D   G   T   D   A   W   D   S   A   T   E
acc aag acc ttc ctg gtg tcc cag gac att cag gat gag ggc tgg gag acc ttg gaa gtg tcc agc gcc gtg aag
 T   K   T   F   L   V   S   Q   D   I   Q   D   E   G   W   E   T   L   E   V   S   S   A   V   K
cgc tgg gtc cgg tcc gac tcc acc aag agc aaa aat aag ctg gaa gtg act gtg gag agc cac agg aag ggc tgc
 R   W   V   R   S   D   S   T   K   S   K   N   K   L   E   V   T   V   E   S   H   R   K   G   C
gac acg ctg gac atc agt gtc ccc cca ggt tcc aga aac ctg ccc ttc ttt gtt gtc ttc tcc aat gac cac agc
 D   T   L   D   I   S   V   P   P   G   S   R   N   L   P   F   F   V   V   F   S   N   D   H   S
agt ggg acc aag gag acc agg ctg gag ctg agg gag atg atc agc cat gaa caa gag agc gtg ctc aag aag ctg
 S   G   T   K   E   T   R   L   E   L   R   E   M   I   S   H   E   Q   E   S   V   L   K   K   L
tcc aag gac ggc tcc aca gag gca ggt gag agc agt cac gag gag gac acg gat ggc cac gtg gct gcg ggg tcg
 S   K   D   G   S   T   E   A   G   E   S   S   H   E   E   D   T   D   G   H   V   A   A   G   S
act tta gcc atc gag ggg cgg gga gac gat gga aac tac tgt aag agg acc ccg ctc tac atc gac ttc aag gag
 T   L   A   I   E   G   R   G   D   D   G   N   Y   C   K   R   T   P   L   Y   I   D   F   K   E
att ggg tgg gac tcc tgg atc atc gct ccg cct gga tac gaa gcc tat gaa tgc cgt ggt gtt tgt ttc ttc ccc
 I   G   W   D   S   W   I   I   A   P   P   G   Y   E   A   Y   E   C   R   G   V   C   F   F   P
ctg gca gag cat ctc aca ccc aca aag cat gca att atc cag acc ttg gtc cac ctc aag aat tcc cag aaa gct
 L   A   E   H   L   T   P   T   K   H   A   I   I   Q   T   L   V   H   L   K   N   S   Q   K   A
tcc aaa gcc tgc tgt gtg ccc aca aag cta gag ccc atc tcc atc ctc tat tta gac aaa ggc gtc gtc acc ctc
 S   K   A   C   C   V   P   T   K   L   E   P   I   S   I   L   Y   L   D   K   G   V   V   T   L
aag tac aaa tac gaa ggc atg gcc gtc tcc gaa tgt ggc tgt aga tag
 K   Y   K   Y   E   G   M   A   V   S   E   C   G   C   R   -
```

### AlbSigPep-His-Albumin-(thr)-ActRIIb

```
atg gtc tgg gtg acc ttc atc agc ctg ctg ttt ctg ttc agc agc gcc tac agc aga ggc gtg ttc aga cgg gac
 M   V   W   V   T   F   I   S   L   L   F   L   F   S   S   A   Y   S   R   G   V   F   R   R   D
gcc cat cac cat cac cat cac aag agc gag gtg gcc cac aga ttc aag gac ctg ggc gag gaa aac ttc aag gcc
 A   H   H   H   H   H   H   K   S   E   V   A   H   R   F   K   D   L   G   E   E   N   F   K   A
ctg gtg ctg atc gcc ttc gcc cag tat ctg cag cag tgc ccc ttc gag gac cac gtg aag ctg gtc aac gaa gtg
 L   V   L   I   A   F   A   Q   Y   L   Q   Q   C   P   F   E   D   H   V   K   L   V   N   E   V
acc gag ttc gcc aag acc tgt gtg gcc gac gag agc gcc gag aac tgc gac aag agc ctg cac acc ctg ttc ggc
 T   E   F   A   K   T   C   V   A   D   E   S   A   E   N   C   D   K   S   L   H   T   L   F   G
gac aag ctg tgc acc gtg gcc acc ctg aga gaa acc tac ggc gag atg gcc gac tgc tgc gcc aag cag gaa cct
 D   K   L   C   T   V   A   T   L   R   E   T   Y   G   E   M   A   D   C   C   A   K   Q   E   P
gag aga aat gaa tgc ttc ttg caa cac aaa gat gac aac cca aac ctc ccc cga ttg gtg aga cca gag gtt gat
 E   R   N   E   C   F   L   Q   H   K   D   D   N   P   N   L   P   R   L   V   R   P   E   V   D
gtg atg tgc act gct ttt cat gac aat gaa gag aca ttt ttg aaa aaa tac tta tat gaa att gcc aga aga cat
 V   M   C   T   A   F   H   D   N   E   E   T   F   L   K   K   Y   L   Y   E   I   A   R   R   H
cct tac ttt tat gcc ccg gaa ctc ctt ttc ttt gct aaa agg tat aaa gct gct ttt aca gaa tgt tgc caa gct
 P   Y   F   Y   A   P   E   L   L   F   F   A   K   R   Y   K   A   A   F   T   E   C   C   Q   A
gct gat aaa gct gcc tgc ctg ttg cca aag ctc gat gaa ctt cgg gat gaa ggg aag gct tcg tct gcc aaa cag
 A   D   K   A   A   C   L   L   P   K   L   D   E   L   R   D   E   G   K   A   S   S   A   K   Q
aga ctc aag tgt gcc agt ctc caa aaa ttt gga gaa aga gct ttc aaa gca tgg gca gta gct cgc ctg agc cag
 R   L   K   C   A   S   L   Q   K   F   G   E   R   A   F   K   A   W   A   V   A   R   L   S   Q
aga ttt ccc aaa gct gag ttt gca gaa gtt tcc aag tta gtg aca gat ctt acc aaa gtc cac acg gaa tgc tgc
 R   F   P   K   A   E   F   A   E   V   S   K   L   V   T   D   L   T   K   V   H   T   E   C   C
cat gga gat ctg ctt gaa tgt gct gat gac agg gcg gac ctt gcc aag tat atc tgt gaa aat caa gat tcg atc
 H   G   D   L   L   E   C   A   D   D   R   A   D   L   A   K   Y   I   C   E   N   Q   D   S   I
tcc agt aaa ctg aag gaa tgc tgt gaa aaa cct ctg ttg gaa aaa tcc cac tgc att gcc gaa gtg gaa aat gat
 S   S   K   L   K   E   C   C   E   K   P   L   L   E   K   S   H   C   I   A   E   V   E   N   D
gag atg cct gct gac ttg cct tca tta gct gct gat ttt gtt gaa agt aag gat gtt tgc aaa aac tat gct gag
 E   M   P   A   D   L   P   S   L   A   A   D   F   V   E   S   K   D   V   C   K   N   Y   A   E
gca aag gat gtc ttc ctg ggc atg ttt ttg tat gaa tat gca aga agg cat cct gat tac tct gtc gtg ctg ctg
 A   K   D   V   F   L   G   M   F   L   Y   E   Y   A   R   R   H   P   D   Y   S   V   V   L   L
ctg aga ctt gcc aag aca tat gaa acc act cta gag aag tgc tgt gcc gct gca gat cct cat gaa tgc tat gcc
 L   R   L   A   K   T   Y   E   T   T   L   E   K   C   C   A   A   A   D   P   H   E   C   Y   A
aaa gtg ttc gat gaa ttt aaa cct ctt gtg gaa gag cct cag aat tta atc aaa caa aat tgt gag ctt ttt gag
 K   V   F   D   E   F   K   P   L   V   E   E   P   Q   N   L   I   K   Q   N   C   E   L   F   E
cag ctt gga gag tac aaa ttc cag aat gcg cta tta gtt cgt tac acc aag aaa gta ccc caa gtg tca act cca
 Q   L   G   E   Y   K   F   Q   N   A   L   L   V   R   Y   T   K   K   V   P   Q   V   S   T   P
act ctt gta gag gtc tca aga aac cta gga aaa gtg ggc agc aaa tgt tgt aaa cat cct gaa gca aaa aga atg
 T   L   V   E   V   S   R   N   L   G   K   V   G   S   K   C   C   K   H   P   E   A   K   R   M
ccc tgt gca gaa gac tat cta tcc gtg gtc ctg aac cag tta tgt gtg ttg cat gag aaa acg cca gta agt gac
 P   C   A   E   D   Y   L   S   V   V   L   N   Q   L   C   V   L   H   E   K   T   P   V   S   D
aga gtc acc aaa tgc tgc aca gaa tcc ttg gtg aac agg cga cca tgc ttt tca gct ctg gaa gtc gat gaa aca
 R   V   T   K   C   C   T   E   S   L   V   N   R   R   P   C   F   S   A   L   E   V   D   E   T
tac gtt ccc aaa gag ttt aat gct gaa aca ttc acc ttc cat gca gat ata tgc aca ctt tct gag aag gag aga
 Y   V   P   K   E   F   N   A   E   T   F   T   F   H   A   D   I   C   T   L   S   E   K   E   R
caa atc aag aaa caa act gca ctt gtt gag ctc gtg aaa cac aag ccc aag gca aca aaa gag caa ctg aaa gct
 Q   I   K   K   Q   T   A   L   V   E   L   V   K   H   K   P   K   A   T   K   E   Q   L   K   A
gtt atg gat gat ttc gca gct ttt gta gag aag tgc tgc aag gct gac gat aag gag acc tgc ttt gcc gag gag
 V   M   D   D   F   A   A   F   V   E   K   C   C   K   A   D   D   K   E   T   C   F   A   E   E
ggt aaa aag ctg gtc gcc gcc tct cag gtt gct ctg gga ctc ggt tca act agt ggt tct ggt gcg cag act aat
 G   K   K   L   V   A   A   S   Q   V   A   L   G   L   G   S   T   S   G   S   G   A   Q   T   N
gcg agt ggt acc ctg gtg cct aga ggc agc cac atg ctg gaa gat ccc gtg cct gag aca aga gag tgc atc tac
 A   S   G   T   L   V   P   R   G   S   H   M   L   E   D   P   V   P   E   T   R   E   C   I   Y
tac aac gcc aac tgg gag ctt gag cgg acc aac cag agc ggc ctg gaa aga tgt gaa ggc gag cag gac aag cgg
 Y   N   A   N   W   E   L   E   R   T   N   Q   S   G   L   E   R   C   E   G   E   Q   D   K   R
ctg cac tgt tac gcc tct tgg aga aac agc agc ggc acc atc gag ctg gtc aag aaa ggc tgc tgg ctg gac gac
 L   H   C   Y   A   S   W   R   N   S   S   G   T   I   E   L   V   K   K   G   C   W   L   D   D
ttc aac tgc tac gac cgg caa gag tgc gtg gcc acc gaa gag aat ccc cag gtg tac ttc tgc tgc tgc gag ggc
 F   N   C   Y   D   R   Q   E   C   V   A   T   E   E   N   P   Q   V   Y   F   C   C   C   E   G
aac ttc tgc aac gag cgg ttc acc cat ctg cct tga
 N   F   C   N   E   R   F   T   H   L   P   -
```

### AlbSigPep-His-Thr-ActRII

```
atg aag tgg gta acc ttt ctc ctc ctc ctc ttc atc tcc ggt tct gcc ttt tct gcg gcc gca ggc agc tct cac
 M   K   W   V   T   F   L   L   L   L   F   I   S   G   S   A   F   S   A   A   A   G   S   S   H
cac cac cat cac cat agc tct ggc ctg gtg cct aga ggc tct cac atg ggc aga tcc gag aca caa gag tgc ctg
 H   H   H   H   H   S   S   G   L   V   P   R   G   S   H   M   G   R   S   E   T   Q   E   C   L
ttc ttc aac gcc aac tgg gag aaa gac cgg acc aac cag aca ggc gtg gaa ccc tgt tac ggc gac aag gac aag
 F   F   N   A   N   W   E   K   D   R   T   N   Q   T   G   V   E   P   C   Y   G   D   K   D   K
cgg aga cac tgc ttc gcc acc tgg aag aac atc agc ggc agc atc gag atc gtg aag caa ggc tgc tgg ctg gac
 R   R   H   C   F   A   T   W   K   N   I   S   G   S   I   E   I   V   K   Q   G   C   W   L   D
gac atc aac tgc tac gac aga acc gac tgc gtg gaa aag aaa gac agc ccc gag gtg tac ttc tgc tgc tgc gag
 D   I   N   C   Y   D   R   T   D   C   V   E   K   K   D   S   P   E   V   Y   F   C   C   C   E
ggc aac atg tgc aac gag aag ttc agc tac ttc ccc gag atg gaa gtg acc tga
 G   N   M   C   N   E   K   F   S   Y   F   P   E   M   E   V   T   -
```

**Figure S1:**

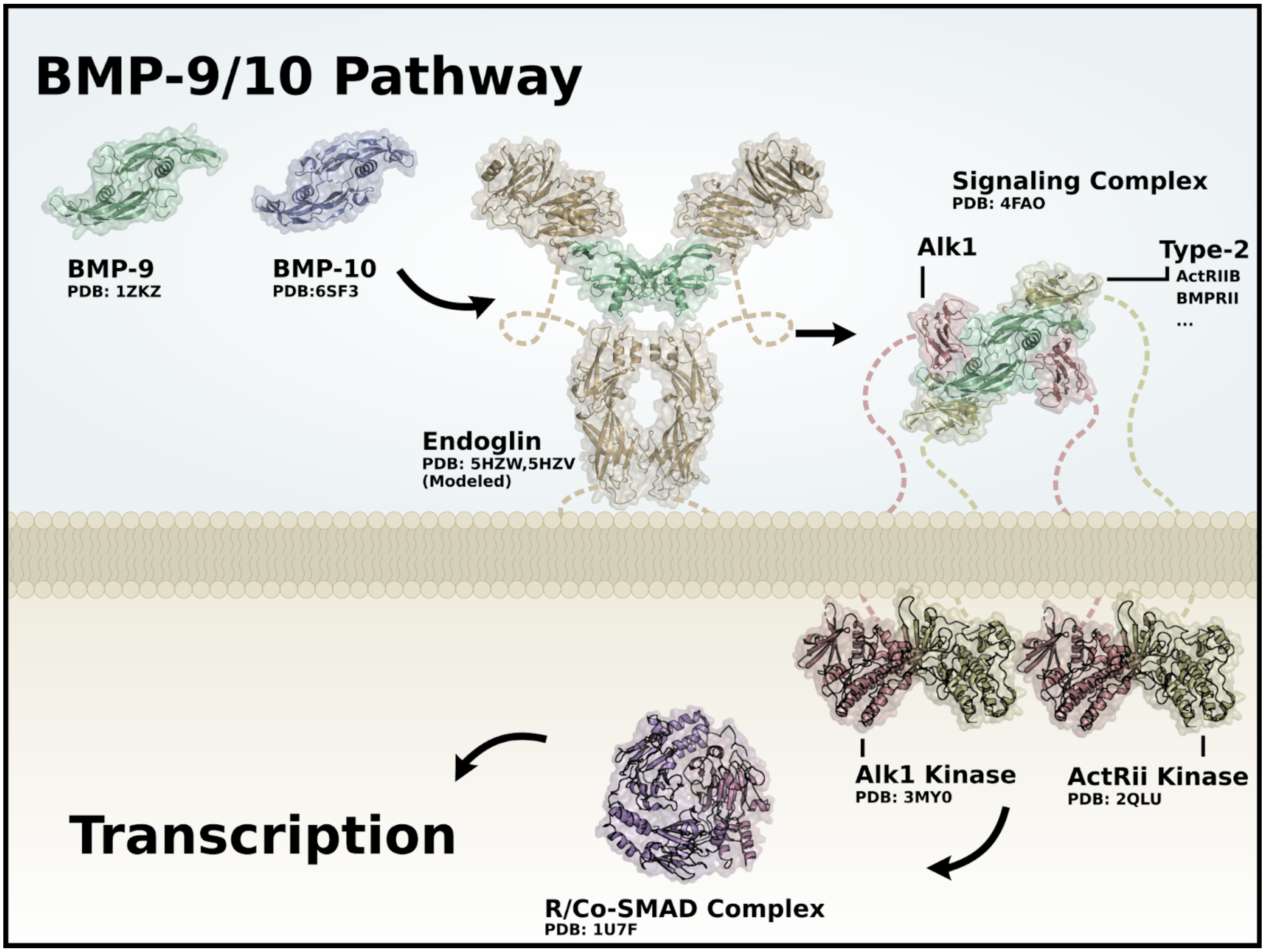


**Figure S1: Model of the BMP-9/10 signaling pathway.** This is a simplified model of the BMP-9/10 signaling pathway. It skips over many specific aspects of the BMP-9/10 pathway such as dimerization, pro-complex release, binding orders, colocalizations, phosphorylation details, conformational shifts, nuclear localization, recruitment and cooperation of transcription factors, and recruitment RNA polymerase.

**Figure S2:**

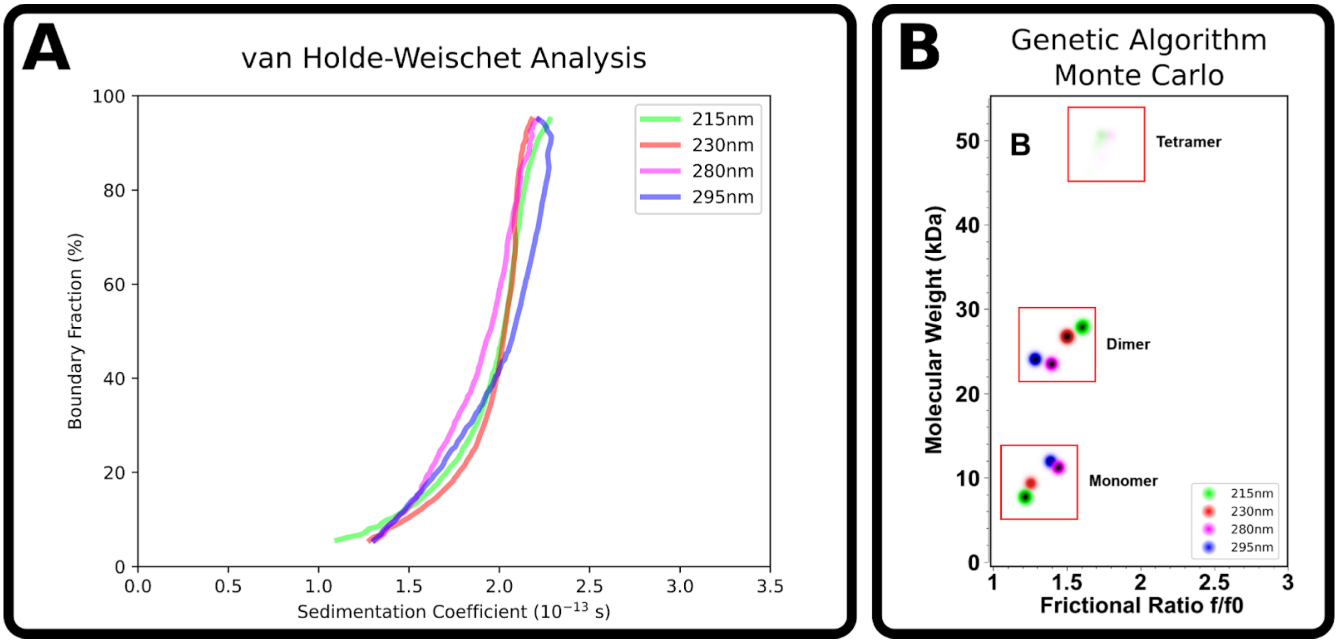


**Figure S2: Analytical ultracentrifugation of cysteinylated BMP-9 monomers.** **(A)** Diffusion-corrected integral sedimentation coefficient distributions of cysteinylated BMP-9 as determined by the enhanced van Holde – Weischet method (1), showing sedimentation coefficients ranging between 1.0-2.5 S. Different concentrations were measured at varying wavelengths. Green: 1.7 µM (215nm), red: 6.2 µM (230nm), blue: 17.6 µM (280nm), magenta: 60.5 µM (295nm). **(B)** Molar mass distribution of cysteinylated BMP-9 as analyzed by genetic algorithm – Monte Carlo analysis (2,3) at the same concentrations and wavelengths as in (A), showing monomer, dimer and tetramer species. Colors are identical to (A), and color intensities reflect partial concentrations for each species.

**Figure S3:**

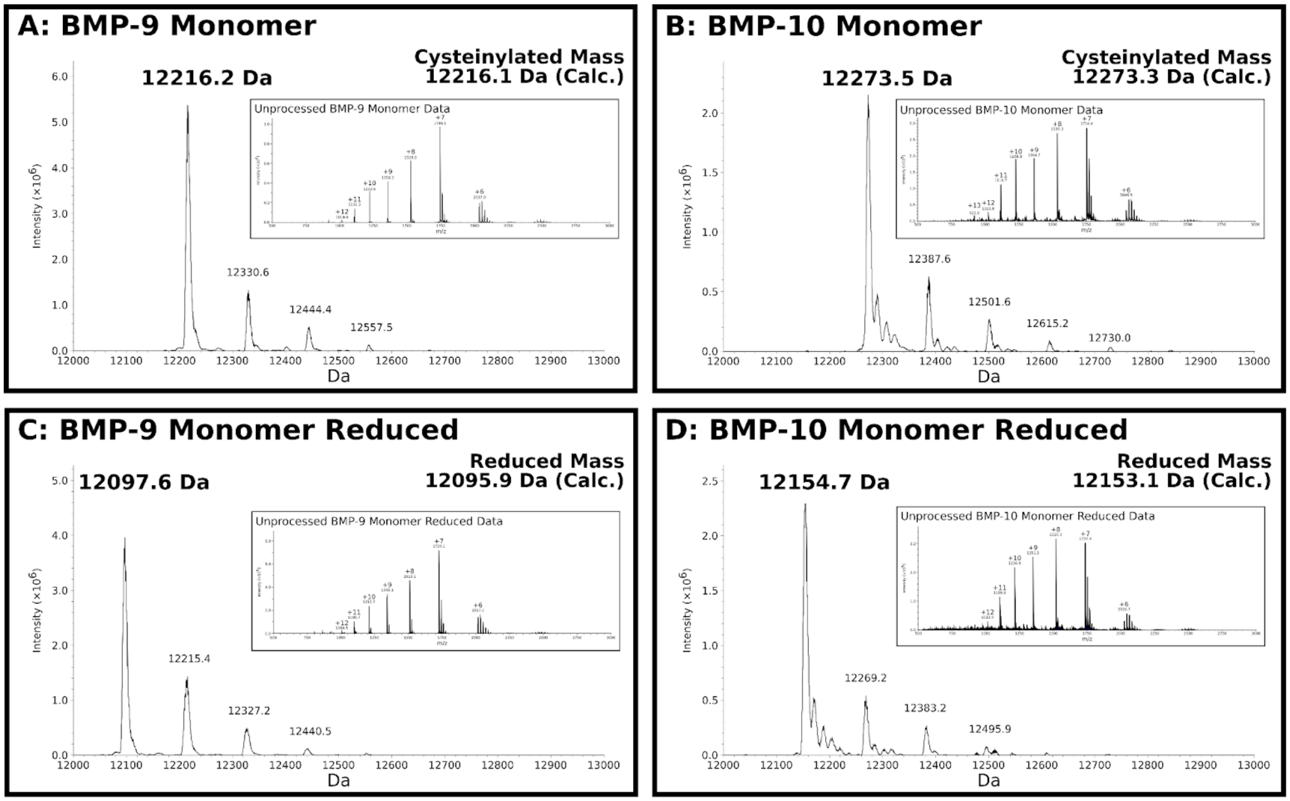


**Figure S3: Mass spectrometry of BMP-9 and BMP-10 monomers is consistent with cysteinylation. (A, B)** Deconvoluted mass spectra of the BMP-9 **(A)** and BMP-10 **(B)** monomers with experimental masses noted above the respective peaks and the calculated cysteinylation mass noted in the upper-right corner of the sub-figures. ( **C, D**) Deconvoluted mass spectra of reduced BMP-9 **(C)** and BMP-10 **(D)** monomers with experimental masses noted above the respective peaks and the calculated reduced mass noted in the upper-right corner of the sub-figures. Insets are the unprocessed ESI-TOF mass spectrometry results.

*BMP-9*: The reduced masses were consistent with a partial reduction where only the solvent exposed Cys-392 was reduced. The (partially) "reduced mass" of our BMP-9 construct (i.e. all cysteines intact, without seven hydrogens) is calculated to be 12095.9229Da. To account for the cysteinylation, one must add the mass of a cysteine (121.16 Da) without one hydrogen (120.16Da) for the total "cysteinylated mass" of 12216.0829 Da. Monomeric BMP-9 will have at least one additional hydrogen present on the inter-chain cysteine (12096.9229Da), plus two hydrogens for each additional reduction of an intra-chain disulfide.

*BMP-10*: Analogous math is performed for BMP-10. The (partially) "reduced mass" of our BMP-10 construct (i.e. all cysteines intact, without seven hydrogens) is calculated to be 12153.1367Da. To account for the cysteinylation, one must add the mass of a cysteine (121.16 Da) without one hydrogen (120.16Da) for the total "cysteinylated mass" of 12273.2967Da. Monomeric BMP-10 will have at least one additional hydrogen present on the inter-chain cysteine (12154.1367Da), plus two hydrogens for each additional reduction of an intra-chain disulfide.

**Figure S4:**

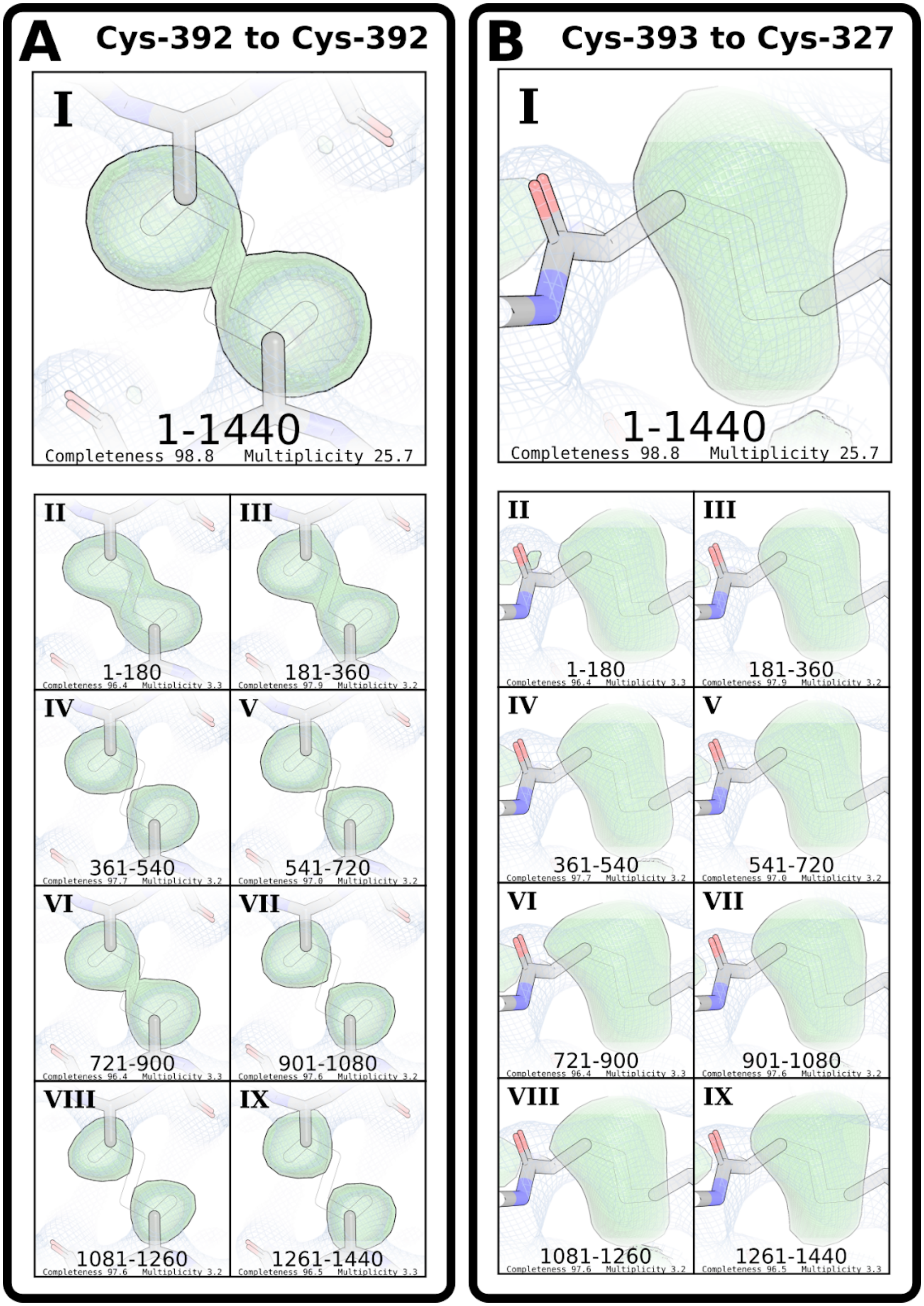


**Figure S4: Single crystal analyses of BMP-9 dimer at neutral pH.** **(A,B)** Sulfur-omit maps of BMP-9 dimer crystallized at a neutral pH with faint outlines showing possible conformations of the inter-chain cysteine Cys-392 **(A)** or intra-chain cystine Cys393-Cys327 **(B)**. In both, full 360° datasets **(I)** are compared to each eighth of the total diffraction images **(II-IX)** as indicated by the large numbers at the bottom of each sub-figure. The direct map is contoured at 1.5σ, and the difference map is contoured at 3.0σ.

### Figure S5:

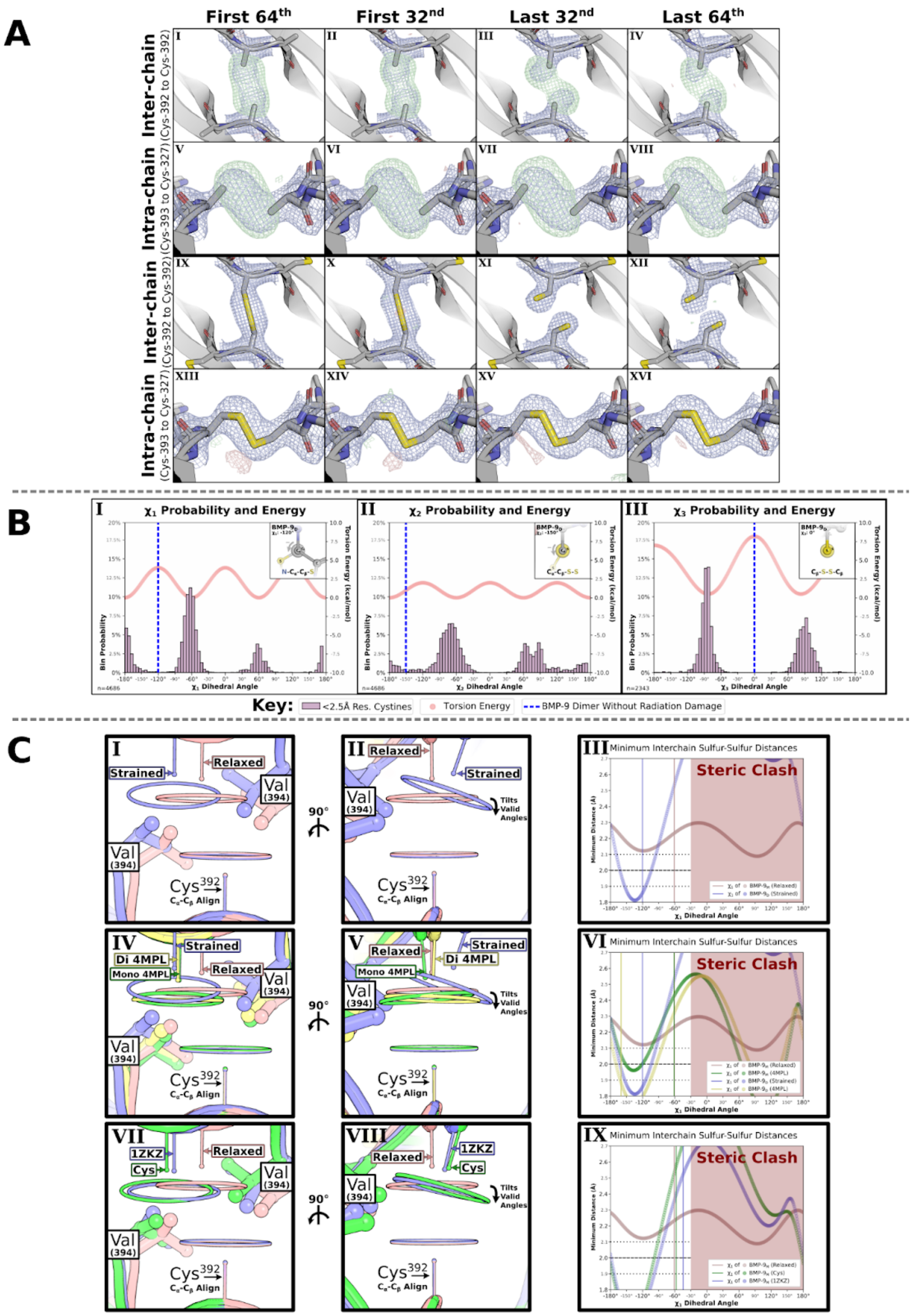

**Figure S5: Extended analysis of neutral-pH Wild-Type BMP-9.** **(A)** Cysteine density analysis using sulfur-omit density maps **(I-VIII)** and modeled density maps **(IX-XVI)** of BMP-9 dimer at neutral pH where the direct map is contoured at 1.5σ, and the difference map is contoured at 3.0σ. **(B)** Plot of $\chi_1$ **(I)**, $\chi_2$ **(II)**, and $\chi_3$ **(III)** dihedral angles with frequency of angle in 2.5Å resolution or better structures and torsion energy as calculated from the AMBER force field with respective dihedral angle plotted as vertical lines for unbroken dimer (strained) dimer. **(C)** Distance analyses with model of interchain cysteine and all rotamer sulfur positions as a halo from two perspectives and the minimal sulfur-sulfur distances as a function of dihedral angle. Comparison of our unbroken dimer (strained) to our relaxed (broken-cystine or non-cysteinylated) monomer structure **(I-III)**. Comparison of cysteinylated monomers, PDB: 1ZKZ and ours to our relaxed monomer **(IV-VI)**. Comparison of PDB: 4MPL conformations to our strained dimer and relaxed monomer **(VII-IX)**. Some components of this figure repeated in Figure 4 and 5.

**Figure S6:**

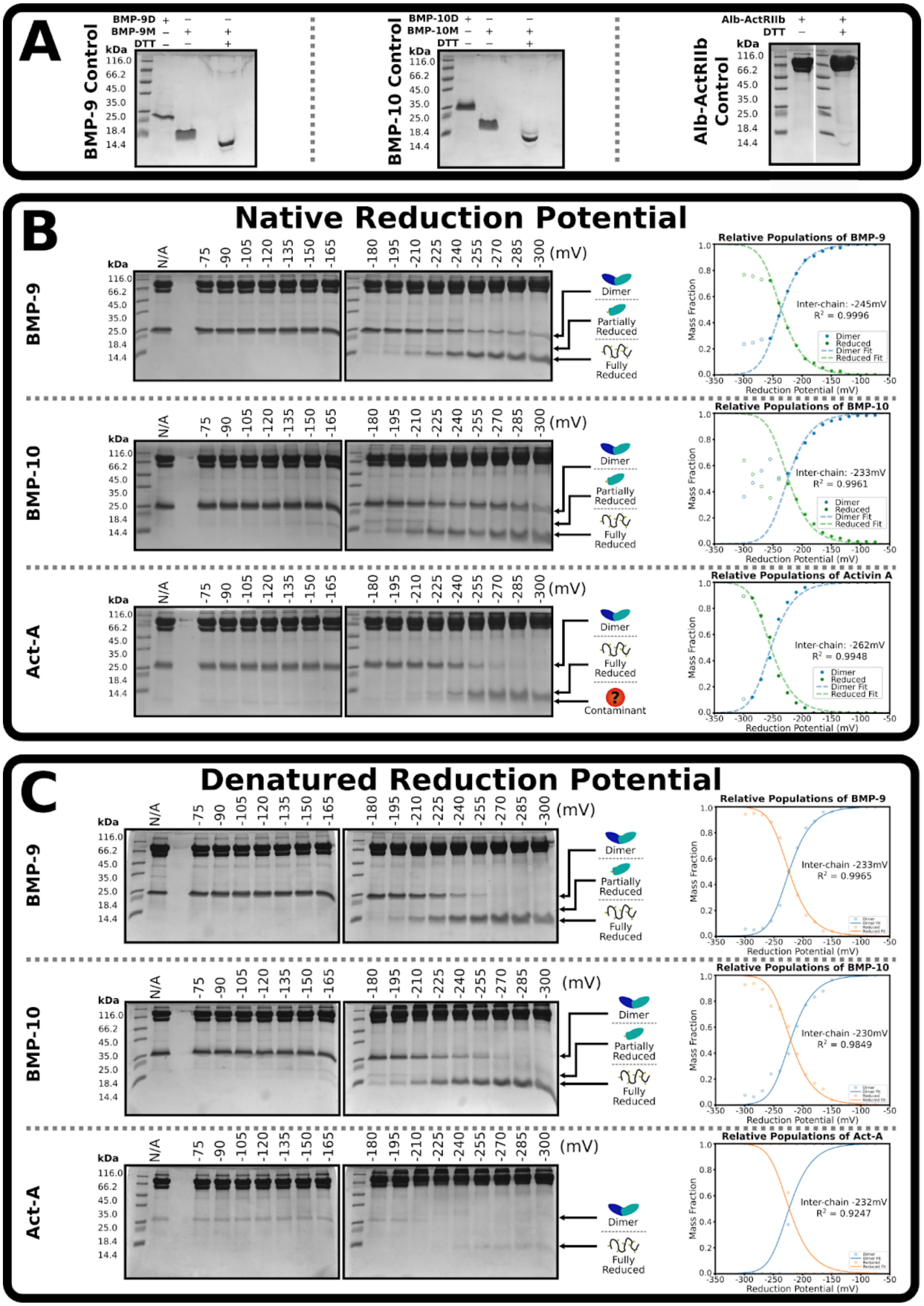


**Figure S6: BMP-9 and BMP-10 have a greater reduction potential than Act-A.** **(A)** Reference of purified dimeric, monomeric, and reduced BMP-9, BMP-10 or Albumin-ActRIIb fusion. **(B)** SDS-PAGE at varying reduction potentials with densitometry of each growth-factor species plotted and fit to Nernst equation. Inter-chain reductions potentials fit to -245mV for BMP-9, -233mV for BMP-10, and -262mV for Act-A. **(C)** Same as **(B)** but reduced with SDS as a denaturant during the reaction. Inter-chain reduction potentials fit to -233mV for BMP-9, -230mV for BMP-10, and -232mV for Act-A.

**Figure S7:**

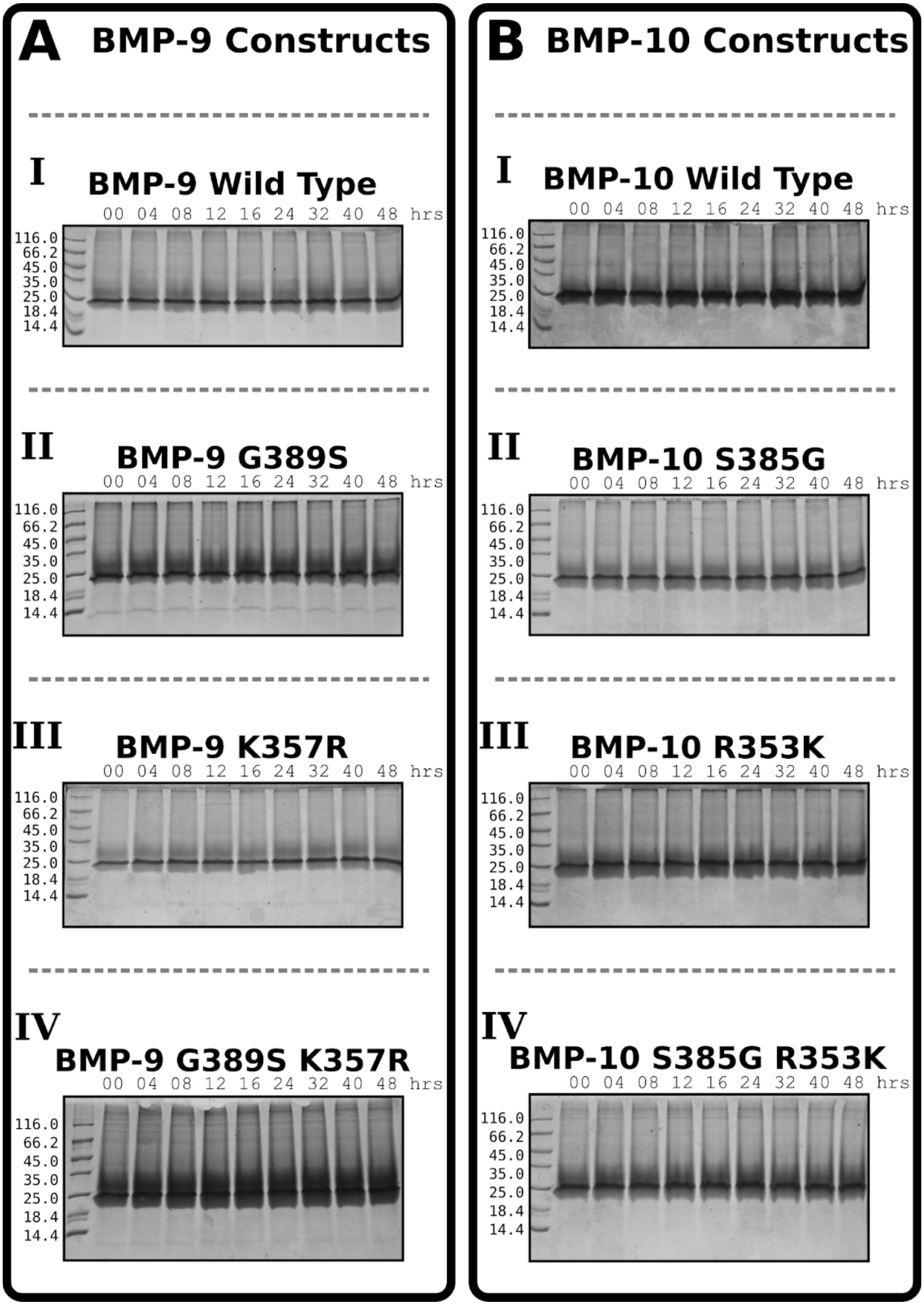


**Figure S7: Evaluation of spontaneous monomerization.** **(A,B)** SDS-PAGE at different time-points over 48hrs where BMP-9 variants **(A)** and BMP-10 variants **(B)** were incubated at pH 8.0 with BMPRII. For both the variants are wild-type **(I)**, glycine/serine mutants **(II)**, lysine/arginine mutants **(III)**, double mutants **(IV)**.

**Figure S8:**

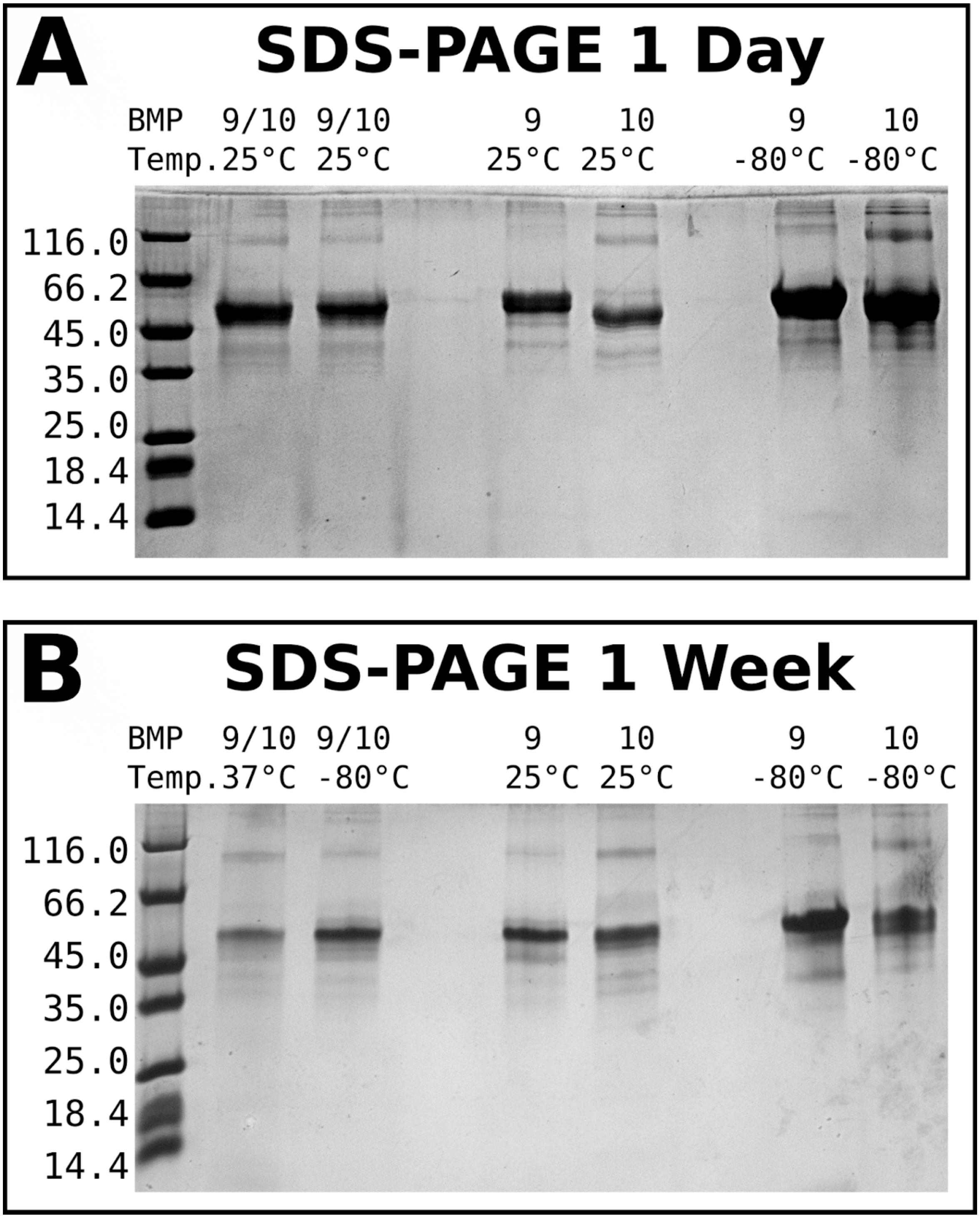


**Figure S8: Evaluation of spontaneous dimerization.** **(A)** Samples of (lane 4) monomeric Pro-BMP-9, (lane 5) monomeric Pro-BMP-10, or (lane 1 and 2) mixtures of monomeric Pro-BMP-9 and monomeric Pro-BMP-10 were left at room temperature for a day and compared to (lanes 7 and 8) -80°C stocks. **(B)** Same samples after a week, but one of the mixtures (lane 1) was placed at 37°C and the other (lane 2) was placed at -80°C.

**Figure S9:**

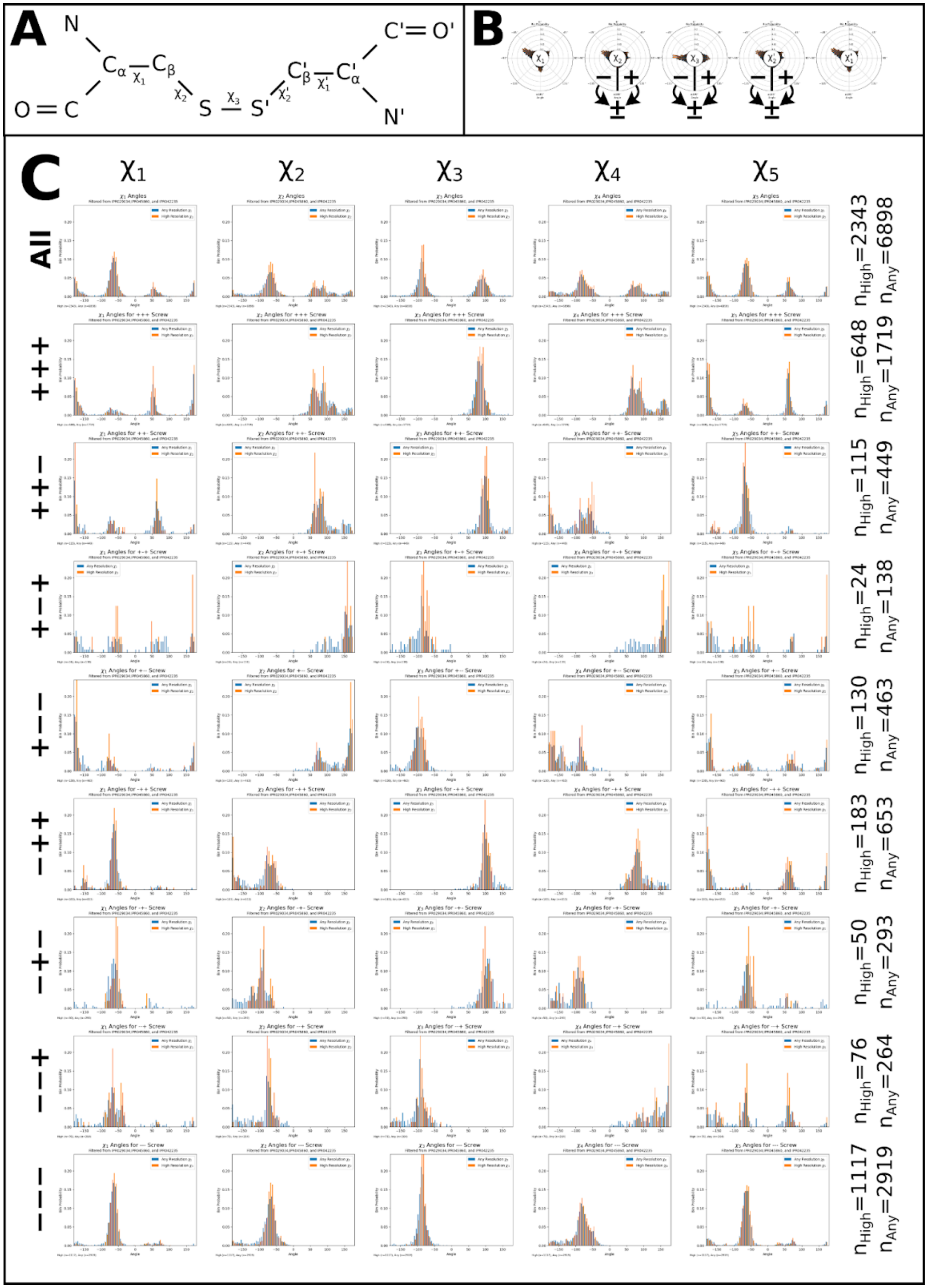


**Figure S9: Cystine screw analysis.** **(A)** Reference diagram of cystine. **(B)** Radial histogram of dihedral angles at low and high resolutions. **(C)** Table of dihedral angle distributions for different cystine screws. Columns are the dihedrals (where $\chi_5$=$\chi_1$', $\chi_4$=$\chi_2$'). Rows are the cystine screw. Right column of labels indicates the number of cystines in histogram.

**Figure S10:**

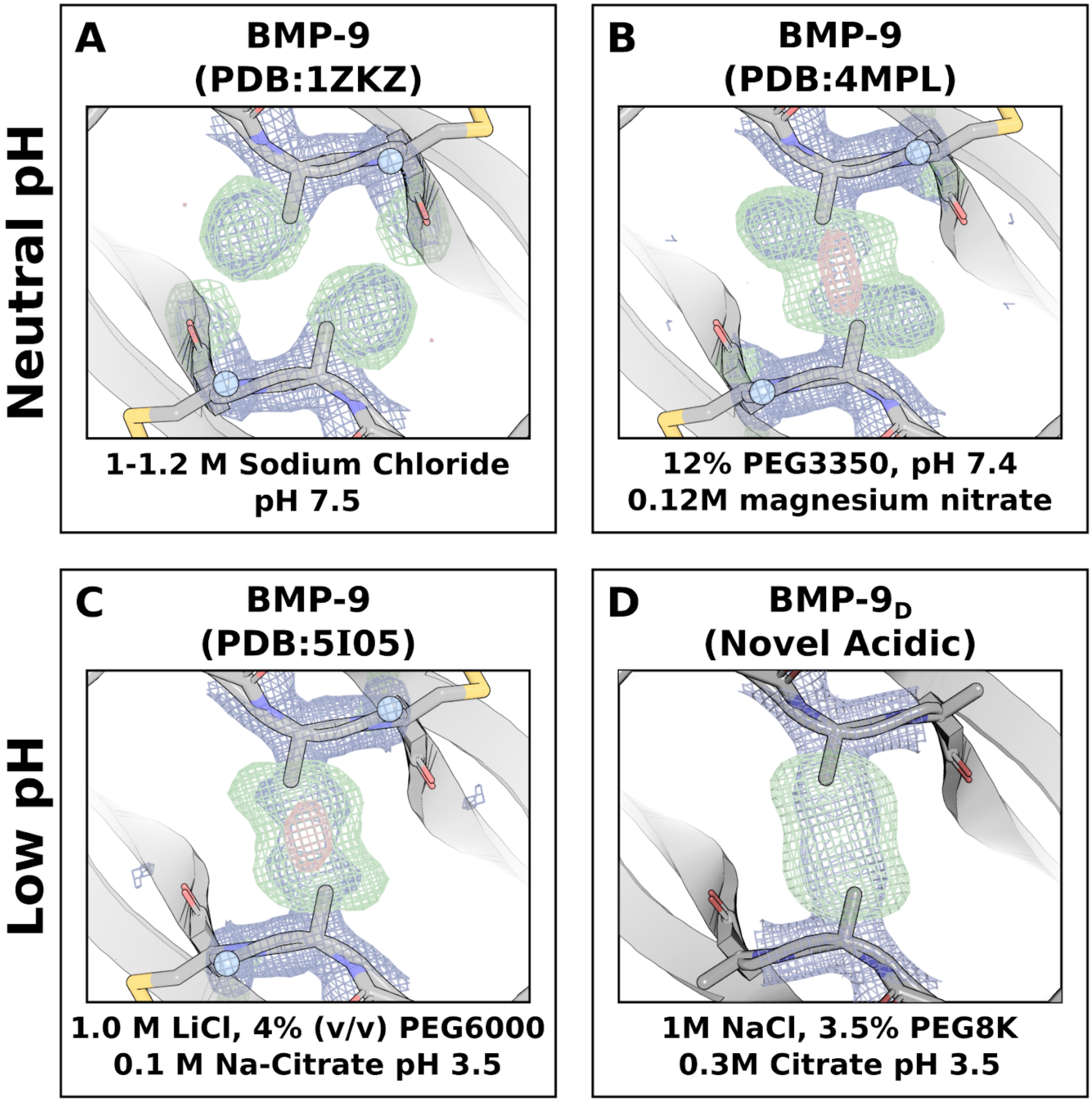


**Figure S10: pH dependence of radiation sensitivity.** (**A-D**) Omit maps of previously published BMP-9 structures' inter-chain bond PDB:1ZKZ **(A)**, PDB: 4MPL **(B)**, PDB: 5I05 **(C)**, and novel low pH BMP-9 dimer structure **(D)** where direct map is contoured at 1.5σ, and the difference map is contoured at 3.0σ. Note that the negative densities in the center of these are above the region of interest.

**Figure S11:**

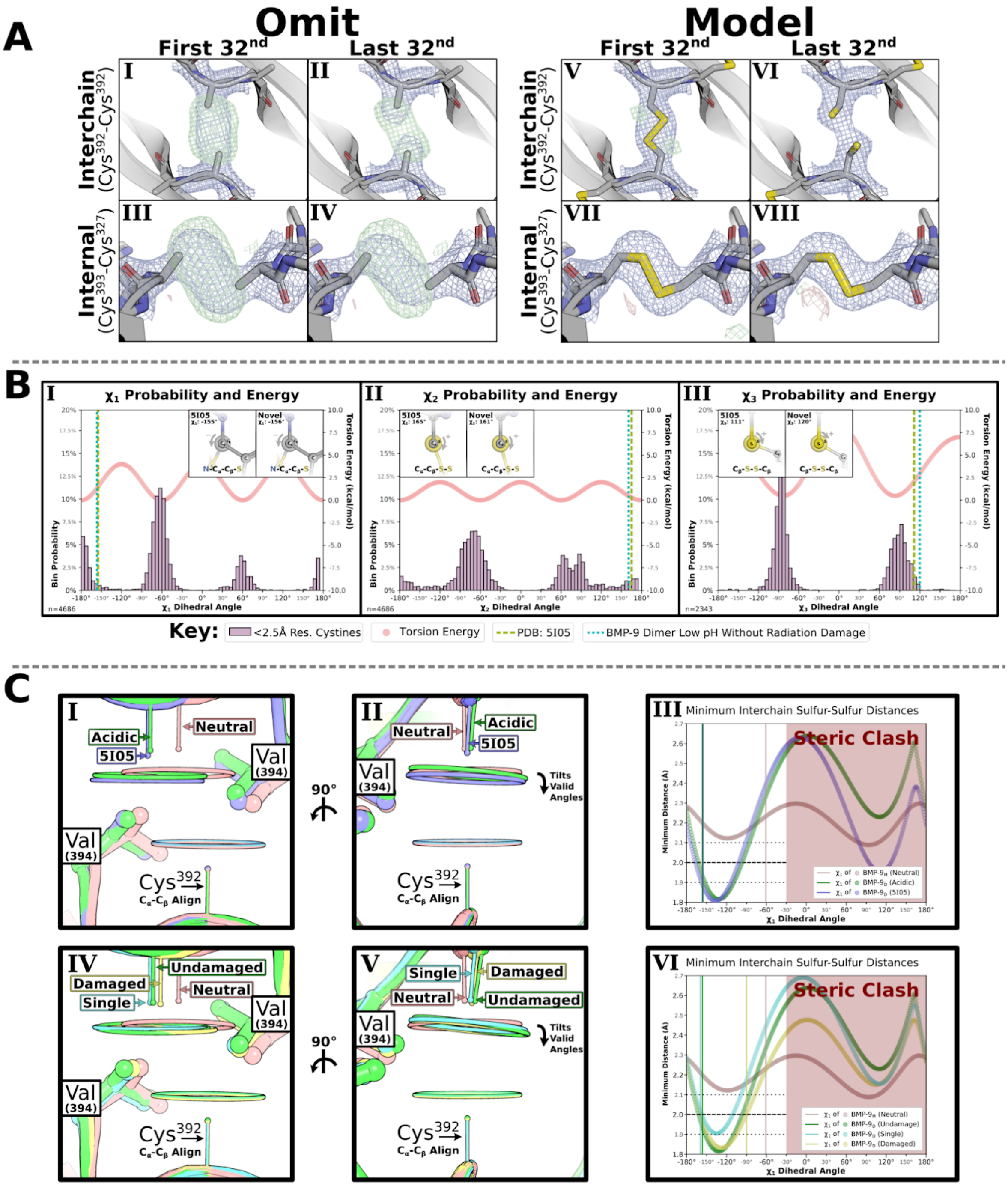


**Figure S11: Extended analysis of low-pH wild type BMP-9 dimer.** **(A)** Cysteine density analysis using sulfur-omit density maps **(I-IV)** and modeled density maps **(V-VIII)** of BMP-9 dimer at low pH where the direct map is contoured at 1.5σ, and the difference map is contoured at 3.0σ. **(B)** Plot of $\chi_1$ **(I)**, $\chi_2$ **(II)**, and $\chi_3$ **(III)** dihedral angles with frequency of angle in 2.5Å resolution or better structures and torsion energy as calculated from the AMBER force field with respective dihedral angle plotted as vertical lines for PDB:5I05 and our our undamaged acidic dimer. **(C)** Distance analyses with model of interchain cysteine and all rotamer sulfur positions as a halo from two perspectives and the minimal sulfur-sulfur distances as a function of dihedral angle. Comparison of PDB:5I05 and our undamaged acidic dimer to our relaxed (broken-cystine or non-cysteinylated) monomer structure **(I-III)**. Comparison of our single-crystal, undamaged, and damaged low pH BMP-9 dimers to our relaxed monomer **(IV-VI)**.

**Figure S12:**

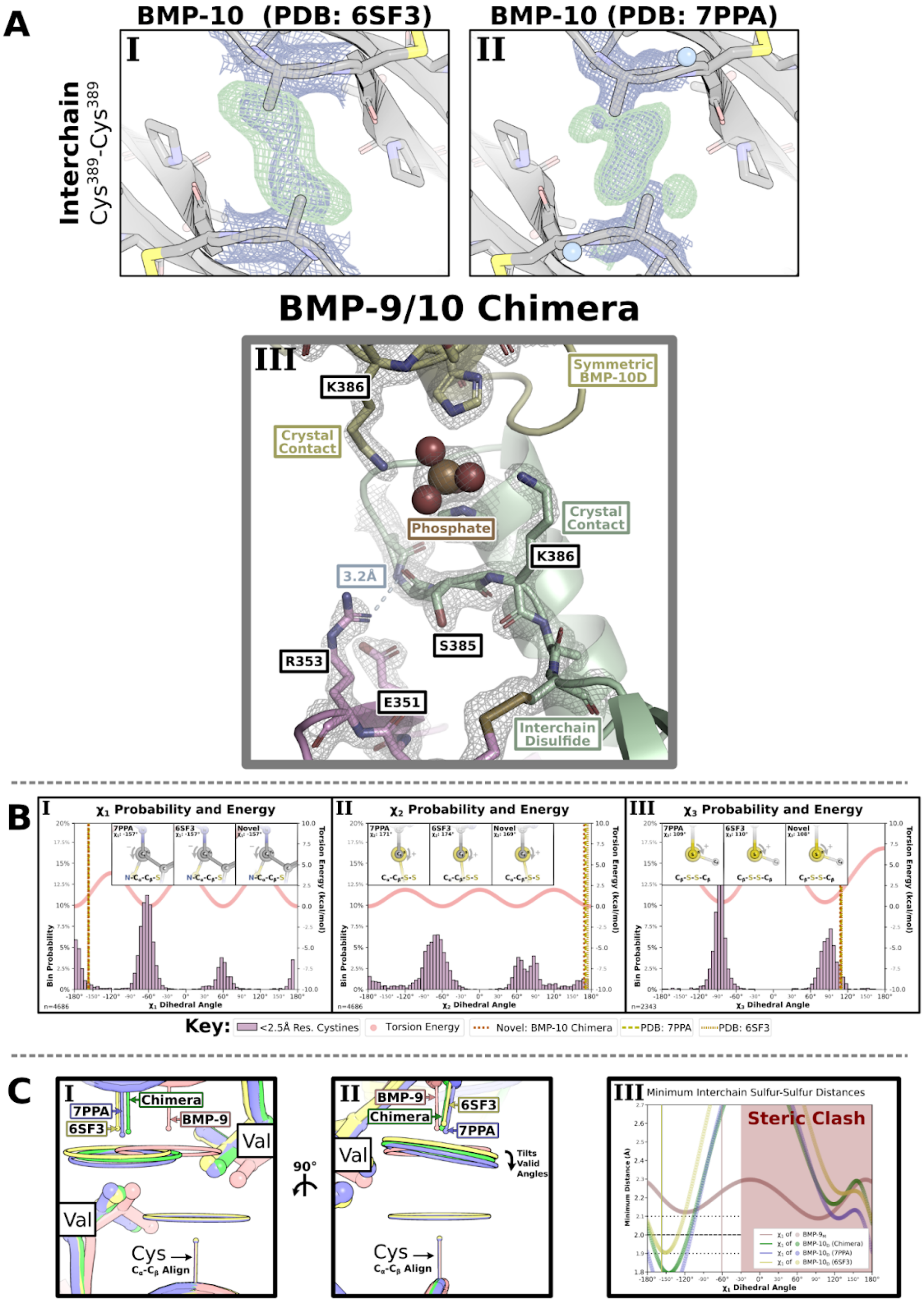


**Figure S12: Extended analysis of BMP-10.** **(A)** Interchain cystine as sulfur omit maps of published BMP-10 PDB: 6SF3 **(I)** and PDB: 7PPA **(II)** and our fully modeled chimeric BMP-9/10 **(III)** where the direct map is contoured at 1.5σ, and the difference map is contoured at 3.0σ. **(B)** Plot of $\chi_1$ **(I)**, $\chi_2$ **(II)**, and $\chi_3$ **(III)** dihedral angles with frequency of angle in 2.5Å resolution or better structures and torsion energy as calculated from the AMBER force field with respective dihedral angle plotted as vertical lines fo PDB:7PPA, PDB:6SF3, and our chimeric BMP-9/10 dimer. **(C)** Distance analyses with model of interchain cysteine and all rotamer sulfur positions as a halo from two perspectives and the minimal sulfur-sulfur distances as a function of dihedral angle that compares PDB:7PPA, PDB:6SF3, and our chimeric BMP-9/10 dimer to our relaxed (broken-cystine or non-cysteinylated) monomer structure.

**Figure S13:**

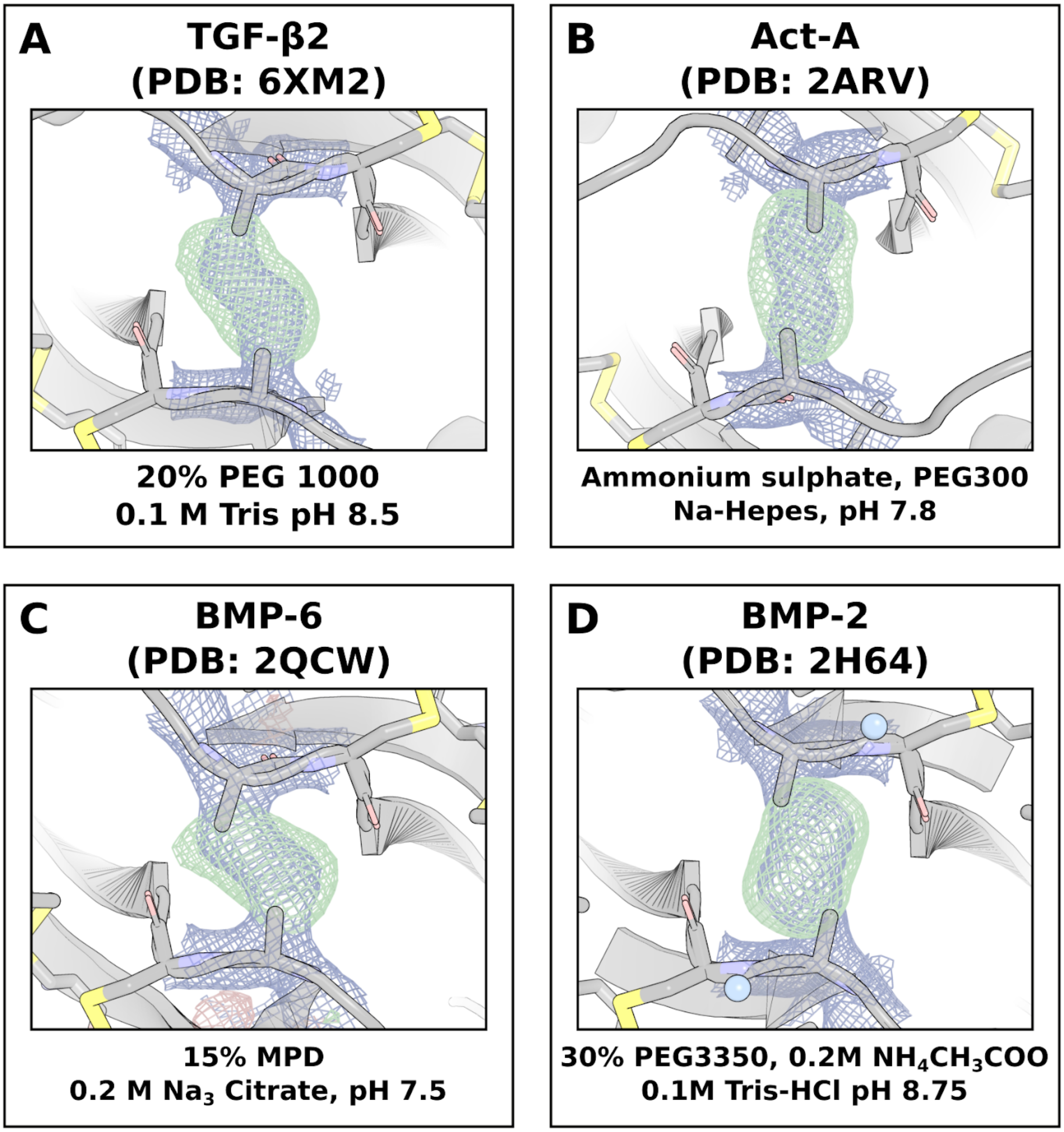


**Figure S13: Published structures of related growth-factors display no radiation sensitivity.** (**A-D**) Omit maps of the inter-chain bond of TGF-β2 PDB:6XM2 **(A)**, Act-A PDB: 2ARV **(B)**, BMP-6 PDB: 2QCW **(C)**, and BMP-2 PDB: 2H64 **(D)**, where the direct map is contoured at 1.5σ and the difference map is contoured at 3.0σ. All maps show fully dimeric protein not indicative of radiation damage.

### Figure S14:

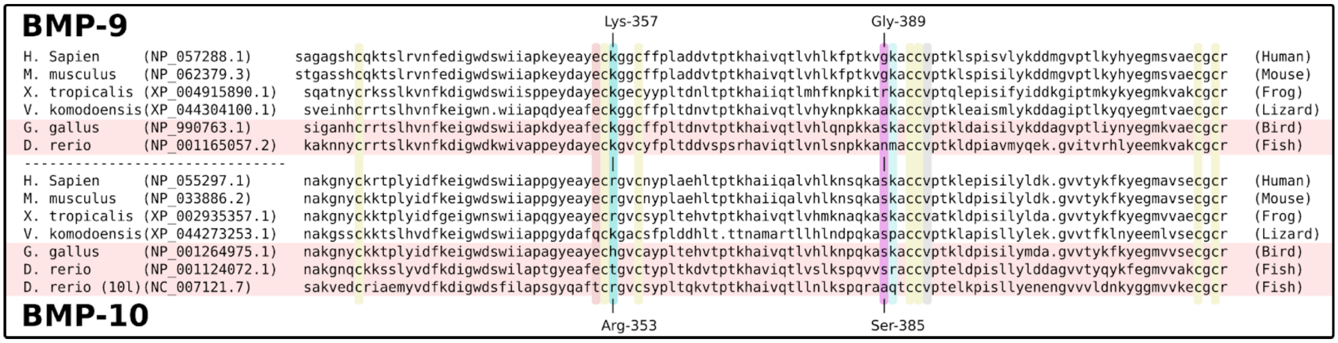


**Figure S14: Aligned BMP-9 and BMP-10 sequences.** Aligned BMP-9 and BMP-10 amino acid sequences from various species. Teal highlights the lysine/arginine mutant. Magenta highlights the glycine/serine mutant. Yellow highlights the cysteines. Gray highlights the valine sterically restricting the interchain cysteine conformations. Light blue highlights Lys-390/386 discussed in S15C,D. Red highlights chicken and zebrafish species as divergent on the glycine/serine residue.

**Figure S15:**

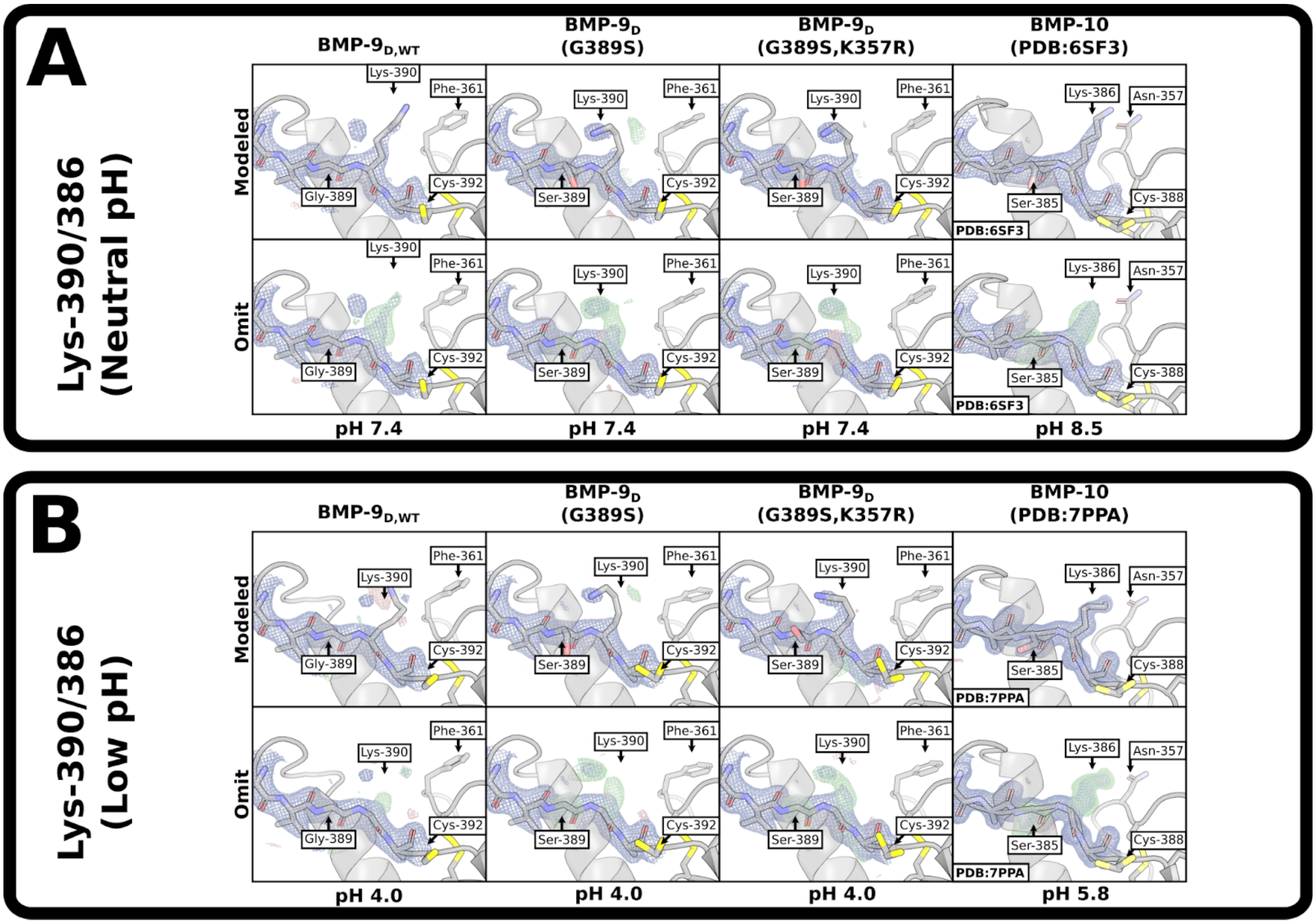


**Figure S15: Residue differences and mutant positions in BMP-9 and BMP-10.** (**A-B**) View of the Lys-390/386 residue position on BMP-9 dimer wild-type, G389S, G389S K357R, and BMP-10 at neutral pH **(A)** and low pH **(B)**. Top rows are fully modeled, bottom rows are omits of relevant residues (Lys-390/386). Direct map is contoured at 1.5σ, and the difference map is contoured at 3.0σ.

**Figure S16:**

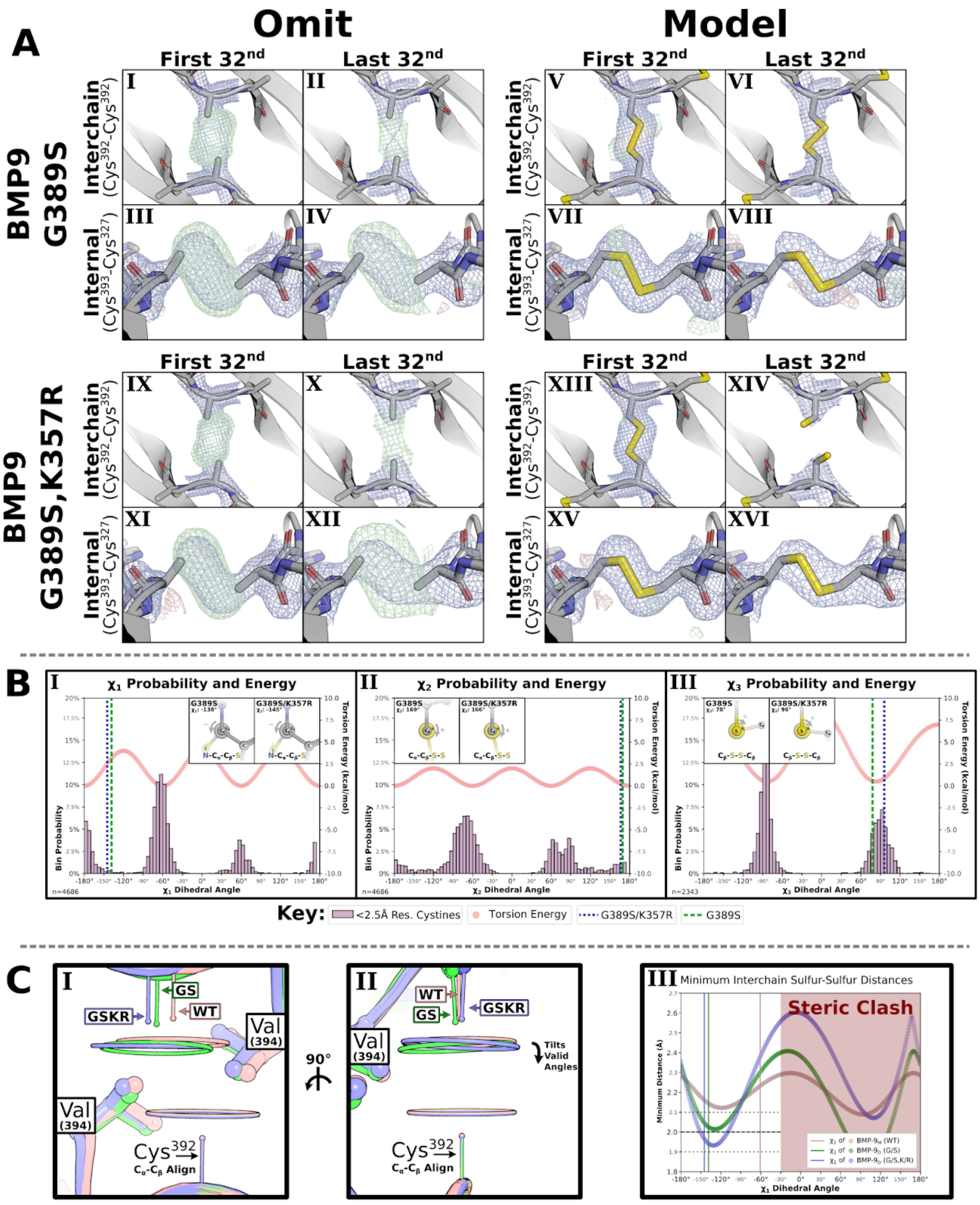


**Figure S16: Extended analysis of neutral-pH mutant BMP-9 dimers.** **(A)** Cysteine density analysis using sulfur-omit density maps (**I-IV,IX-XII**) and modeled density maps **(V-VIII,XIII-XVI)** of BMP-9 G389S dimer **(I-VIII)** and BMP-9 G389S, K357R dimer **(IX-XVI)** at neutral pH where the direct map is contoured at 1.5σ, and the difference map is contoured at 3.0σ. **(B)** Plot of $\chi_1$ **(I)**, $\chi_2$ **(II)**, and $\chi_3$ **(III)** dihedral angles with frequency of angle in 2.5Å resolution or better structures and torsion energy as calculated from the AMBER force field with respective dihedral angle plotted as vertical lines for BMP-9 G389S and BMP-9 G389S, K357R at neutral pH. **(C)** Distance analyses with model of interchain cysteine and all rotamer sulfur positions as a halo from two perspectives and the minimal sulfur-sulfur distances as a function of dihedral angle that compares BMP-9 G389S and BMP-9 G389S, K357R at neutral pH to our relaxed (broken-cystine or non-cysteinylated) monomer structure. Some components of this figure repeated in Figure 7.

**Figure S17:**

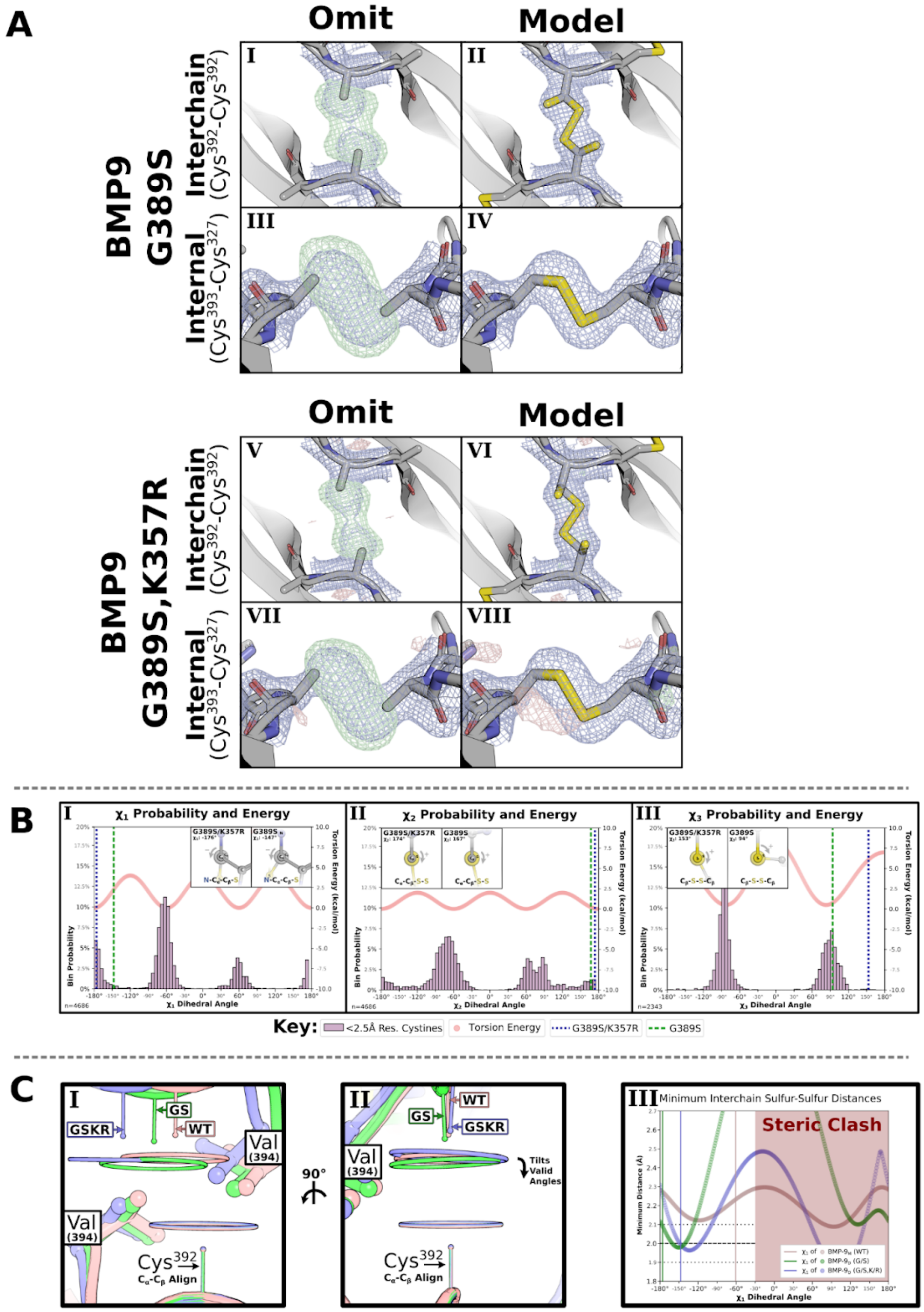


**Figure S17: Extended analysis of low-pH mutant BMP-9 dimers.** **(A)** Interchain cystine analysis of BMP-9 G389S dimer **(I-IV)** and BMP-9 G389S, K357R dimer **(V-VIII)** at low pH using sulfur omit maps and modeled density maps where the direct map is contoured at 1.5σ, and the difference map is contoured at 3.0σ. **(B)** Plot of $\chi_1$ **(I)**, $\chi_2$ **(II)**, and $\chi_3$ **(III)** dihedral angles with frequency of angle in 2.5Å resolution or better structures and torsion energy as calculated from the AMBER force field with respective dihedral angle plotted as vertical lines for BMP-9 G389S and BMP-9 G389S, K357R at low pH. **(C)** Distance analyses with model of interchain cysteine and all rotamer sulfur positions as a halo from two perspectives and the minimal sulfur-sulfur distances as a function of dihedral angle that compares BMP-9 G389S and BMP-9 G389S, K357R at low pH to our relaxed (broken-cystine or non-cysteinylated) monomer structure. Some components of this figure repeated in Figure 7.
